# Protein semisynthesis underscores role of a conserved lysine in activation and desensitization of acid-sensing ion channels

**DOI:** 10.1101/2023.01.23.525296

**Authors:** Debayan Sarkar, Iacopo Galleano, Stephanie Andrea Heusser, Sofie Yuewei Ou, Gül Refika Uzun, Keith K. Khoo, Gerbrand Jan van der Heden van Noort, Joseph Scott Harrison, Stephan Alexander Pless

**Affiliations:** Department of Drug Design and Pharmacology, University of Copenhagen, 2100 Copenhagen, Denmark; Department of Cell and Chemical Biology, Leiden University Medical Centre, Leiden, The Netherlands; Department of Chemistry, University of the Pacific, Stockton, California, USA

**Author notes:** Co-corresponding author: Stephan Pless, Department of Drug Design and Pharmacology, University of Copenhagen, Jagtvej 160, 2100 Copenhagen, Denmark, Mail. These authors contributed equally (Debayan Sarkar, Iacopo Galleano).

## Abstract

Acid-sensing ion channels (ASICs) are trimeric ion channels that open a cation-conducting pore in response to proton binding. Excessive ASIC activation during prolonged acidosis in conditions such as inflammation and ischemia is linked to pain and stroke. A conserved lysine in the extracellular domain (Lys211 in mASIC1a) is suggested to play a key role in ASIC function. However, the precise contributions are difficult to dissect with conventional mutagenesis, as replacement of Lys211 with naturally occurring amino acids invariably changes multiple physico-chemical parameters. Here, we study the contribution of Lys211 to mASIC1a function using tandem protein-trans splicing (tPTS) to incorporate non-canonical lysine analogs. We conduct optimization efforts to improve splicing and functionally interrogate semisynthetic mASIC1a. In combination with molecular modeling, we show that Lys211 charge and side chain length are crucial to activation and desensitization, thus emphasizing that tPTS can enable atomic-scale interrogations of membrane proteins in live cells.

## INTRODUCTION

Acid-sensing ion channels (ASICs) are trimeric ligand-gated ion channels that open a cation-selective pore in response to proton binding to their extracellular domain (ECD). ^1^ They contribute to fast synaptic neurotransmission and have been associated with pathological conditions of both the peripheral nervous system (e.g. pain, diabetic neuropathy) and the central nervous system (e.g. stroke, fear, neurodegenerative diseases).^2-4^ Consequently, ASICs are emerging as possible drug targets for both small molecules and peptide-/animal toxin-derived lead molecules.^5^

Structural evidence has confirmed that each ASIC subunit contains intracellular N- and C-terminal domains and a transmembrane domain (TMD) encompassing two transmembrane helices that are linked to the large ECD.^6^ Binding of protons to the ECD triggers a series of conformational changes that lead to opening of the channel pore.^6-9^ This transient pore opening is typically followed by a process termed fast desensitization, during which the channel enters a non-conductive state in the continued presence of high proton concentrations.^10^ Alternatively, exposure to less elevated proton levels can evoke conformational changes that evoke a non-conductive state without prior pore opening. This process is termed steady-state desensitization (SSD).^10^

Previous functional and structural work has identified Lys211 (mASIC1a numbering) to play an important role in channel gating in different ASIC isoforms, as well as across ASICs from different species ^6,11-13^, likely because it forms a H-bond network with Leu351 and Asp355 of the neighboring subunit (Figure 1A). Additionally, Lys211 has been suggested to contribute to a state-dependent binding site for Cl-ions.^14^ Together with Arg309 and Glu313 from the adjacent subunit (Figure 1A), this anion binding site was implicated in the process of fast desensitization and had been suggested to affect acid-induced cell death.^12,13^

**Figure 1:**
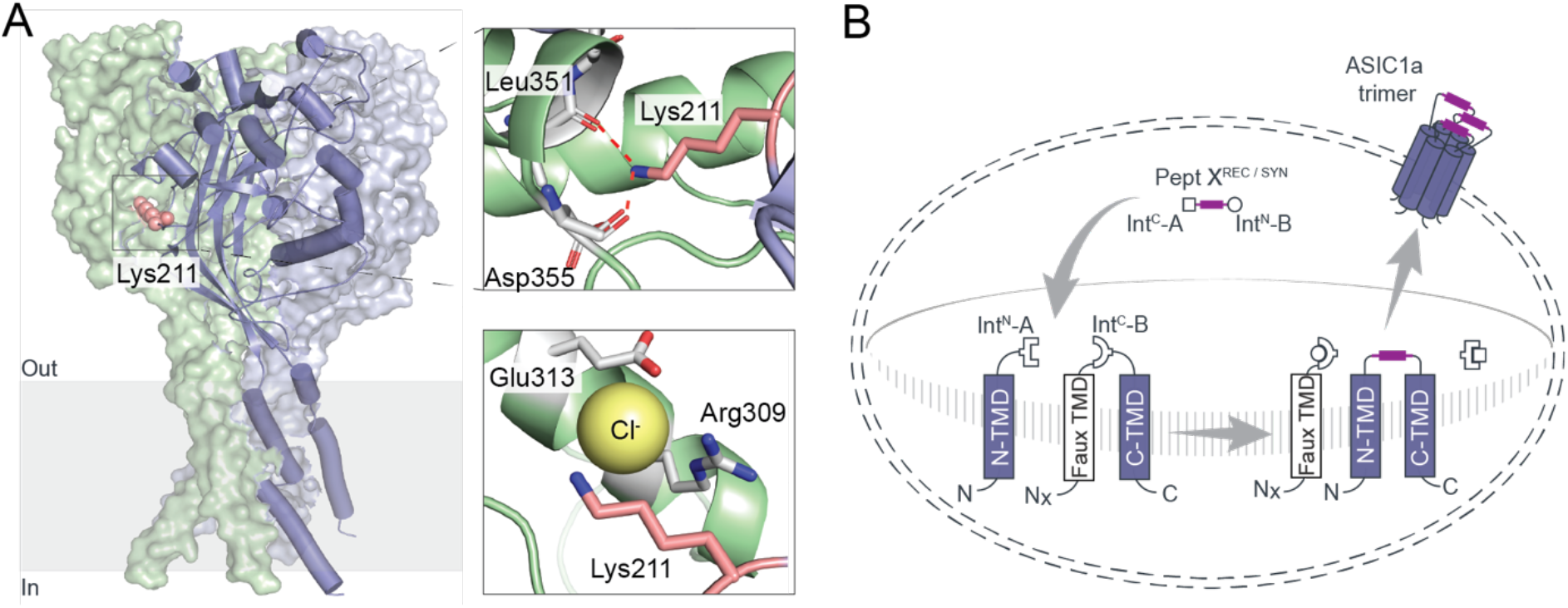
A semisynthetic strategy to probe the contributions of Lys211 to the Cl--binding site in the ASIC ECD. (A) Side view of a cASIC1 channel (PDB:2qts). Subunit in front is shown in violet cartoon representation with the Lys211 side chain highlighted as salmon-colored spheres and the two remaining subunits are depicted with light grey and light green surfaces. The top inset shows the proposed H-bond network between the Lys211 amine and the main chain carbonyl oxygen of Leu351 and side chain carboxylate of Asp355 ^6,11^. The bottom inset shows the Cl--binding pocket in cASIC1 (PDB: 2qts), in which the Cl-ion is predicted to be coordinated by the side chains of Lys211, Arg309 and Glu313. (B) Schematic overview of the strategy to reconstitute full-length mASIC1a. This is achieved by recombinantly expressing two separate mASIC1a fragments comprising the first transmembrane helix and the N-terminal part of the ECD (N-TMD) and the C-terminal part of the ECD and the second transmembrane helix (C-TMD; note the *faux* transmembrane domain (Faux TMD) at its N-terminus to ensure correct topology) in the presence of a recombinantly expressed or synthetically prepared peptide X in *Xenopus laevis* oocytes (Pept X^REC^ and Pept X^SYN^, respectively). Tandem protein trans-splicing in the presence of peptide X results in formation of full-length mASIC1a monomers that trimerize to form functional ion channels. Inteins A and B are indicated by square and round symbols, respectively.

However, conventionally employed mutagenic approaches, such as side chain deletion or substitution by alanine, are likely to cause global disruptions and can therefore complicate data interpretation. This is especially true for side chains situated at the functionally important subunit interface of ASICs or the related epithelial sodium channels, ENaCs.^15-18^ A high-resolution dissection of a possible contribution to channel activation and fast desensitization of Lys211 therefore requires the incorporation of Lys analogs with subtle changes in side chain length or pKa.

A widely applied approach to incorporate such non-canonical amino acid (ncAA) analogs into membrane proteins is by genetic code expansion.^19,20^ However, protein yields, non-specific incorporation or lack of sufficiently effective discrimination by the amino-acyl tRNA synthetase between ncAA side chains with similar physico-chemical properties can be challenging. In some cases, these issues can be overcome by alternative approaches such as protein semisynthesis using protein trans-splicing (PTS).^21-24^PTS involves formation of a peptide bond between two proteins or protein fragments (termed as exteins) by means of traceless native chemical ligation mediated by a split intein. Split inteins consist of an N-terminal part (Int^N^) and a C-terminal part (Int^C^), which spontaneously recombine to form an active intein.^25-27^ If the site of interest is situated far from the N-or C-terminus of the protein, the approach can be expanded to tandem protein trans-splicing (tPTS), in which two orthogonal split inteins are used to insert a synthetic peptide into a protein.^19,28,29^ This strategy has been successfully used to incorporate ncAAs in the ECD of the ATP-gated P2X2 receptor, in addition to incorporating post-translational modifications in intracellular linkers of the cardiac sodium channel Na_v_1.5.^29,30^ tPTS requires the N- and C-terminal fragments of the ion channel to be expressed in a cell by recombinant means, while the synthetically produced peptide (Pept X^SYN^) can be introduced into the cell by micro-injection or cell squeezing, depending on cell type.^29^

Here, we aim to insert a recombinant (Pept X^REC^) or a synthetic peptide (Pept X^SYN^) into the mASIC1a ECD using the strategy outlined in Figure 1B. In the recombinantly expressed N-terminal and C-terminal transmembrane constructs (N-TMD and C-TMD), split intein Int^N^-A is placed at the C-terminus of the N-TMD fragment, whereas the orthogonal Int^C^-B is positioned at the N-terminus of the C-TMD fragment. The latter contains a *faux* transmembrane helix (faux-TMD) that is introduced to preserve the correct topology of the C-TMD fragment in the membrane during protein trafficking (^28,29^, Figure 1B). To reconstitute full-length mASIC1a using tPTS, Int^C^-A is situated at the N-terminus of peptide X, while Int^N^-B is attached to its C-terminus. For further details on the approach see. ^28,29^

With a goal to establish and optimize tPTS to incorporate ncAA Lys analogs at position 211 in mASIC1a, we systematically screened 11 orthogonal split intein pairs across two different splice sites, conducted extein sequence optimization, and evaluated the potential of covalently linking N-TMD and C-TMD constructs. Our results, based on reconstitution of WT mASIC1a (i.e. using Pept X^REC^), establish the orthogonal *Cfa*DnaE-*Ssp*DnaB^M86^ split intein pair as ideal for tPTS in the mASIC1a ECD. We then generated semisynthetic mASIC1a variants by inserting four ncAA analogs of Lys via Pept X^SYN^ at position 211, complemented by molecular modeling to assess chloride binding and conformational dynamics. Together, we provide direct evidence for the importance of side chain charge and length for chloride binding and thus activation and fast desensitization properties of mASIC1a. Overall, we demonstrate the potential of tPTS for the evaluation of functional properties of ASICs with atomic resolution.

## RESULTS

### Optimizing splice site location and choice of split intein pairs

Previous studies with orthogonal split inteins used for protein trans-splicing have indicated that the splicing efficiency and yield depend on several factors. These include presence of nucleophilic Cys, Ser or Thr side chains, location of splice site, amino acids sequence present in close proximity to the splice sites, and orthogonality among split intein pairs.^25,28,31,32^ We therefore set out to identify the optimal combination of splice site, orthogonal split intein pair and sequence context in order to replace Lys211 in mASIC1a.

First, we evaluated the sequence context around Lys211 for the presence of nucleophilic Cys, Ser or Thr side chains that would enable splicing with no or minimal alterations of the native mASIC1a sequence, depending on the nucleophile preference of different split inteins. This led to the identification of two similar but distinct splice site combinations: For splice site combination #1, we chose the N-TMD, peptide X (including Lys211) and C-TMD fragments to span mASIC1a residues 1-193, 194-213 and 214-526, respectively; those chosen for splice site combination #2 spanned residues 1-179, 180-213, and 214-526, respectively (Figure 2A). This enabled us to take advantage of the presence of either a Cys (Cys194) (splice site combination #1) or a Ser (Ser180) (splice site combination #2) at the N terminus of peptide X, to accommodate split inteins with either Cys- and Ser-dependent splicing mechanisms. The C-terminal splice junction at Thr214 was kept identical for both splice site combination #1 and #2 because the next Ser/Cys residues in mASIC1a sequence are situated over 25 amino acids further downstream, which would have necessitated a much longer peptide X. However, due to concerns over cross-reactivity with Thr-based split inteins such as *Tvo*VMA ^33^, this required us to mutate Thr214 to Cys or Ser (depending on the split intein in use) to ensure efficient splicing.

**Figure 2:**
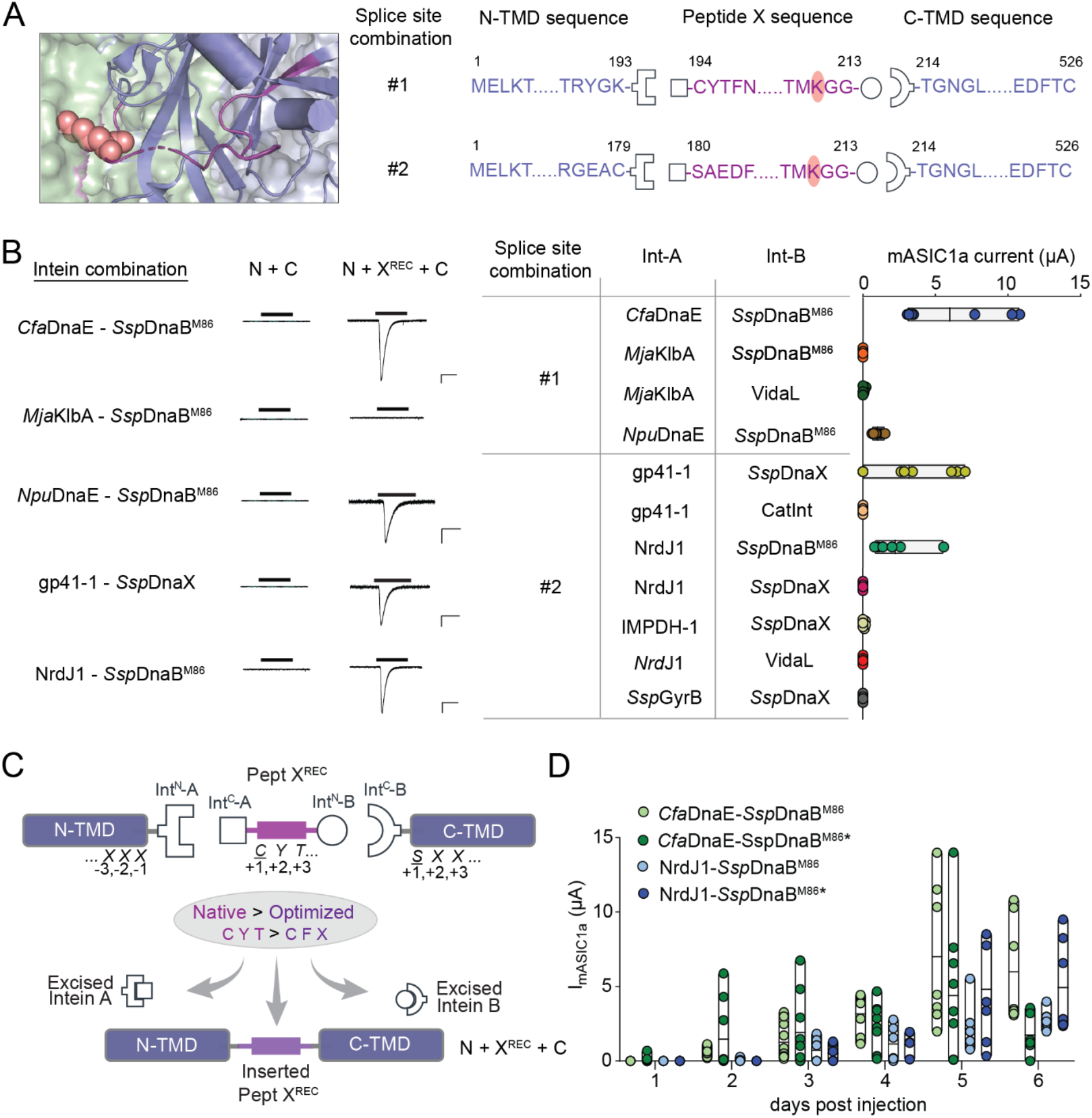
Evaluation of different split intein combinations and sequence context to optimize for highest current amplitudes. (A) Left panel: Structure of cASIC1 (same color-coding as in Figure 1A), with the part of the ECD to be inserted as Pept X highlighted in purple and Lys211 in salmon spheres. Right panel: Two splice site combinations were used in this study. The sequence margins of the N-and C-TMD fragments (violet) and the Pept X^REC^ (purple) shown along with the intein A (square symbols) and B (round symbols) parts linked to them. (B) Left panel: Representative traces of mASIC1a currents from five different split intein combinations. Note that the extein sequence has not been altered beyond the Thr214Ser/Cys substitution for the C-TMD constructs. Black bars indicate application of pH 6.0 to oocytes expressing only N- and C-TMD fragments (N + C, left) or in the presence of a recombinantly expressed Pept X^REC^ using the indicated split intein combination (N + X + C, right). Scale bars: x, 10 s; y, 1 μA. Right panel: List of all split inteins combinations tested here categorized based on splice site combination, along with resulting maximal mASIC1a-mediated currents in *Xenopus laevis* oocytes elicited by application of pH 6.0 (n=6-10 for each combination). (C) Schematic of the trans-splicing reaction with the relevant extein sequence residues in N-TMD, C-TMD and peptide X denoted with -1, -2, -3, +1, +2, +3, which corresponded to positions that were altered to optimize splicing. In this example, the split intein sequences corresponding to *Cfa*DnaE have been shown, where sequence optimization was required only for peptide X. The native (mASIC1a) sequence is shown in purple and optimized sequence is depicted in violet (left panel). (D) Daily progression of reconstituted mASIC1a current levels in oocytes compared between native (mASIC1a sequence) and optimized (*) extein sequences for two different combinations of split inteins (Intein A: *Cfa*DnaE or NrdJ1; Intein B: *Ssp*DnaBM86 (both)) (right panel) (n=6-11 for both combinations). Floating bars depict the mean along with minimum and maximum current margins.

Next, we generated mASIC1a constructs with four different split intein pairs at splice site combination #1 (*Cfa*DnaE-*Ssp*DnaB^M86^, *Mja*KlbA-*Ssp*DnaB^M86^, *Npu*DnaE-*Ssp*DnaB^M86^, and *Mja*KlbA-VidaL) and seven different split intein pairs at splice site combination #2 (NrdJ1-*Ssp*DnaB^M86^, NrdJ1-*Ssp*DnaX, NrdJ1-VidaL, gp41-1-*Ssp*DnaX, gp41-1-CatInt, IMPDH-1-*Ssp*DnaX, and *Ssp*GyrB-*Ssp*DnaX). In all cases, we used the native mASIC1a sequence as the extein sequence context, except for a Thr to Ser/Cys mutation at position 214 position of the C-TMD fragments (see Resource Table T1 for sequence details). We then translated the corresponding N-TMD, Pept X^REC^ and C-TMD fragments into mRNA and injected them into *Xenopus laevis* oocytes for recombinant expression. This was followed by measurements of whole-cell mASIC1a currents elicited by application of pH 6.0 over a period of 6 days using two-electrode voltage clamp (Supplemental Figure S1). For these recordings we made the following assumptions (see also ^29^): the absence of current would indicate no or incomplete splicing, while the presence of pH-induced currents would originate from fully reconstituted mASIC1a constructs, where the current amplitude is roughly indicative of the splicing efficiency.

As expected, expression of just the N-TMD and C-TMD fragments did not yield currents for any of the constructs (Figure 2B). This confirms that the N-TMD and C-TMD fragments do not yield functional channels in absence of Pept X^REC^. But when co-expressed with Pept X^REC^ mRNA, we observed robust currents for two of the split intein pairs at splice site combination #1 (*Cfa*DnaE as split intein A, *Ssp*DnaB^M86^ as split intein B; *Npu*DnaE as split intein A, *Ssp*DnaB^M86^ as split intein B, respectively) and two of the split intein pairs at splice site combination #2 (NrdJ1 as split intein A, *Ssp*DnaB^M86^ as split intein B and gp41-1 as split intein A, *Ssp*DnaX as split intein B, respectively) (Figure 2B). However, even for the most promising split intein combination (*Cfa*DnaE as split intein A, *Ssp*DnaB^M86^ as split intein B), currents rarely exceeded amplitudes of 10 μA, even after multiple days of incubation. This is significantly less than what can be achieved from expression of full-length WT mASIC1a, which typically yields >20 μA within about 24 hrs. We therefore sought to improve splicing yields and/or efficiency by optimizing the sequence context of the different split inteins tested here to more closely resemble their natural sequence context.

### Extein sequence optimization did not enhance current amplitudes

Several previous studies have demonstrated that the sequence contexts of the exteins, especially the -1, -2, -3, +1, +2, +3 positions around the split inteins can influence the efficiency and yields of splicing reactions of split inteins (see Figure 2C, left panel, ^21,29,33-36^). It is thus conceivable that the extein sequence context of the mASIC1a sequence around the splice sites impacts splicing efficiency. We aimed to address this potential issue by testing if mutating the (native) mASIC1a sequence to that of the split intein in its original context would improve apparent splicing yields.

To this end, we therefore introduced mutations in the N-TMD, Pept X^REC^ or C-TMD constructs for all 11 split intein combinations to reflect the natural extein sequence context of each tested split intein, which we refer to as optimized sequences. Resource Tables T2 and T3 outline the mutations incorporated for each split intein combination. Notably, extein sequence optimization failed to elicit pH-induced currents in all seven split intein combinations that had already yielded either negligible or no mASIC1a currents with their natural sequence context (*Mja*KlbA – *Ssp*DnaB^M86^, gp41-1 – CatInt, *Mja*KlbA – VidaL, NrdJ1 – *Ssp*DnaX, IMPDH1 – *Ssp*DnaX, NrdJ11 – VidaL and *Ssp*GyrB – *Ssp*DnaX; see Supplemental Figure S1). Further, extein sequence-optimized constructs with the *Npu*DnaE – *Ssp*DnaB^M86^ and gp41-1 – *Ssp*DnaX split intein combinations no longer elicited detectable pH-induced currents, suggesting a lack of functional channel reconstitution after sequence optimization. By contrast, sequence optimization of the *Cfa*DnaE - *Ssp*DnaB^M86^ split intein pair increased current amplitudes on day 2, although maximally observed current amplitudes (around day 5) were not affected (Figure 2D). Finally, the NrdJ1 - *Ssp*DnaB^M86^ split intein pair showed increased current amplitudes on day 6, yet not exceeding current amplitudes obtained with the *Cfa*DnaE - *Ssp*DnaB^M86^ split intein pair.

In conclusion, our sequence optimization attempts proved largely unsuccessful, even detrimental, to splicing, as assessed by current amplitudes. However, the robust currents observed with the native mASIC1a extein sequence context of the *Cfa*DnaE - *Ssp*DnaB^M86^ split intein pair (except for the Thr214Ser mutation), made this particular split intein combination the most promising candidate for additional optimization attempts.

### Covalently linking N-TMD and C-TMD did not increase current amplitudes

We next sought to test if a forced spatial proximity of the recombinantly expressed N-TMD and C-TMD fragments would increase splicing yields and thereby current amplitudes. To this end, we designed a construct in which N-TMD and C-TMD were linked by a 30-amino acid flexible linker, consisting of 6 repeats of a GlyGlyGlyGlySer sequence stretch (Figure 3A; note that this design eliminates the need for a *Faux* TMD). Injecting mRNA for this construct (NC linked) and the corresponding peptide X^REC^ fragment yielded pH-sensitive currents, but we did not observe currents in control experiments where only the mRNA for NC linked was injected (Figure 3B). However, maximal current amplitudes for NC linked + peptide X^REC^ remained consistently lower than for those observed for the separately expressed N-TMD + X^REC^ + C-TMD constructs (Figure 3C). We therefore chose the latter to proceed for a more detailed characterization, followed by insertion of synthetic peptides.

**Figure 3:**
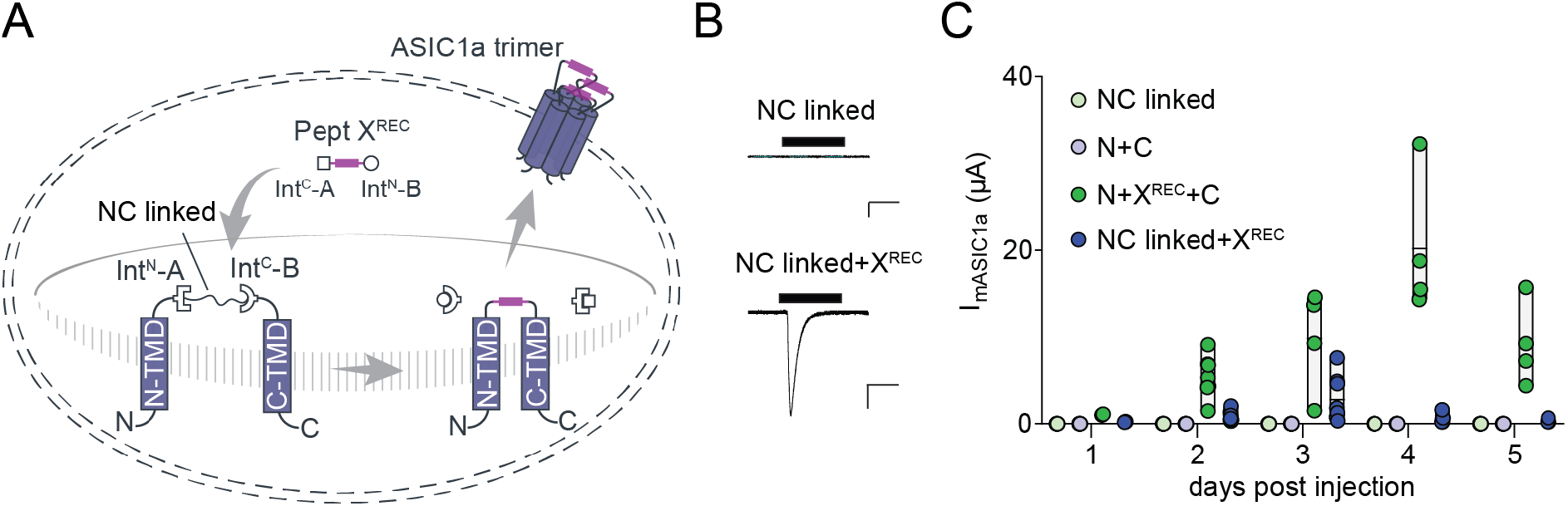
Forced proximity of N-TMD and C-TMD fragments does not appear to promote splicing. (A) Schematic overview of the strategy to reconstruct full-length mASIC1a from a single recombinantly expressed mASIC1a construct with N- and C-terminal fragments that were connected by a 30 amino acid (GGGGS)_6_ linker, termed NC linked. Inteins A (*Cfa*DnaE) and B (*Ssp*DnaB^M86^) are indicated by square and round symbols, respectively. (B) Representative traces of currents from *Xenopus laevis* oocytes expressing only the NC linked construct or NC linked in presence of the Pept X^REC^ fragment (NC linked + X^REC^) and subjected to application of pH 6.0. Scale bars: x, 10 s; y, 1 μA. (C) Daily progression of maximal pH 6.0-induced mASIC1a currents in oocytes separately expressing N + C (see Fig 2B) or the NC linked construct in the presence or absence of X^REC^ (n=4-8 for each combination). Floating bars depict the mean along with minimum and maximum current margins.

### Thr214Ser mutation leads to altered pH sensitivity of activation and SSD

Based on the above results, the *Cfa*DnaE - *Ssp*DnaB^M86^ split intein pair appeared to display the highest splicing yields. Because this split intein pair required the Thr214Ser mutation at the +1 position of the C-TMD fragment to enable *Ssp*DnaB^M86^-mediated splicing (no splicing was observed with the native Thr), we sought to investigate if this sequence alteration caused functional perturbations in mASIC1a function (Supplemental Figure S2). To this end, we first compared the pH sensitivity of activation and steady-state desensitization (SSD) of full-length wildtype mASIC1a channels to full-length mASIC1a channels containing the Thr214Ser mutation. This showed that the mutation itself caused a right-shift in both pH_50_ of activation and SSD (6.7 ± 0.1 vs 6.5 ± 0.1 (p=0.0003) and 7.2 ± 0.0 vs 6.9 ± 0.0 (p<0.0001), respectively), suggesting an overall lower sensitivity to protons (Supplemental Figure S2B and Supplemental Table T1). We expected that recombinantly expressed N-TMD+X^REC^+C-TMD (referred to as N+X^REC^+C) constructs (with the *Cfa*DnaE and *Ssp*DnaB^M86^ split intein pair and hence the Thr214Ser mutation) would yield mASIC1a channels with properties similar to full-length Thr214Ser channels. However, we found the spliced channels to display a minor right-shift in the pH_50_ of activation and a slight left-shift in the pH_50_ of SSD (6.5 ± 0.1 vs 6.4 ± 0.1 (p=0.006) and 6.9 ± 0.0 vs 7.1 ± 0.0 (p<0.0001) for N+X^REC^+C and full-length Thr214Ser, respectively; see Supplemental Figure S2B and Supplemental Table T1). The reason for these small but significant changes remains unclear. The near-identical properties, however, for both full-length and recombinantly reconstituted mASIC1a channels containing the Thr214Ser mutation and the robust expression of the latter led us to next attempt insertion of synthetic peptides in order to assess the role of Lys211 in mASIC1a function.

### ncAA lysine analogs at position 211 reveal importance of side chain length and charge to channel function

In order to assess the contributions of Lys211 side chain charge and length to mASIC1a function, we aimed to incorporate four different ncAA analogs of lysine: ornithine (Orn; protonated amino moiety like Lys, but one carbon shorter), homolysine (hLys; protonated amino moiety like Lys, but one carbon longer), hydroxyl nor-leucine (NleuOH; isosteric to Lys but uncharged) and thialysine (thiaLys; with a decrease in pKa of around one unit and a marginally altered bond geometry (bond angle around 90 degrees, instead of 109.5 degrees) compared to Lys).^37^ We used solid phase peptide synthesis (SPPS) to generate Pept X^SYN^ containing either Lys, Orn, hLys, NleuOH or thiaLys in position 211 (see Supplemental information for details). For functional characterization of channels containing the Lys analogs, we injected Pept X^SYN^ into *Xenopus laevis* oocytes that were pre-injected with the N-TMD and C-TMD fragment mRNAs ∼24 hrs in advance.^29^ Electrophysiological assessments of pH_50_ of activation and SSD were performed 24-48 hrs after peptide injection because cells injected with synthetic peptides typically showed increased mortality beyond 48 hrs after peptide injection. Reassuringly, all constructs generated robust currents (Figure 4A) and control experiments in which a Lys-bearing synthetic peptide was inserted (Lys211^SYN^) showed near-identical pH_50_ values for activation and SSD when compared to mASIC1a channels reconstituted from only recombinantly expressed components N+X^REC^+C (6.4 ± 0.1 and 7.1 ± 0.0, respectively; Figure 4A/B and Supplemental Table T2). This was similar for Orn-containing peptide X^SYN^ (Lys211Orn^SYN^), indicating that shortening of the side chain has little impact on channel function (activation pH_50_ = 6.2 ± 0.1; SSD pH_50_ = 7.1 ± 0.1). By contrast, increasing side chain length (Lys211hLys^SYN^), altering its pKa (Lys211thiaLys^SYN^) or removing the positive charge altogether (Lys211NleuOH^SYN^) produced a drastic right shift in the activation pH_50_ (5.2 ± 0.1 for hLys and NleuOH; 5.3 ± 0.1 for thiaLys), while showing no or relatively minor change in pH_50_ of SSD (7.0 ± 0.0 for hLys, 7.1 ± 0.1 for thiaLys and 7.2 ± 0.0 for NleuOH) (Figure 4B and Supplemental Table T2). Together, this indicates that a shorter side chain length at Lys211 has little functional impact but increasing side chain length or altering/removing its charge is detrimental to proton sensitivity of activation and, to a lesser extent, SSD.

**Figure 4:**
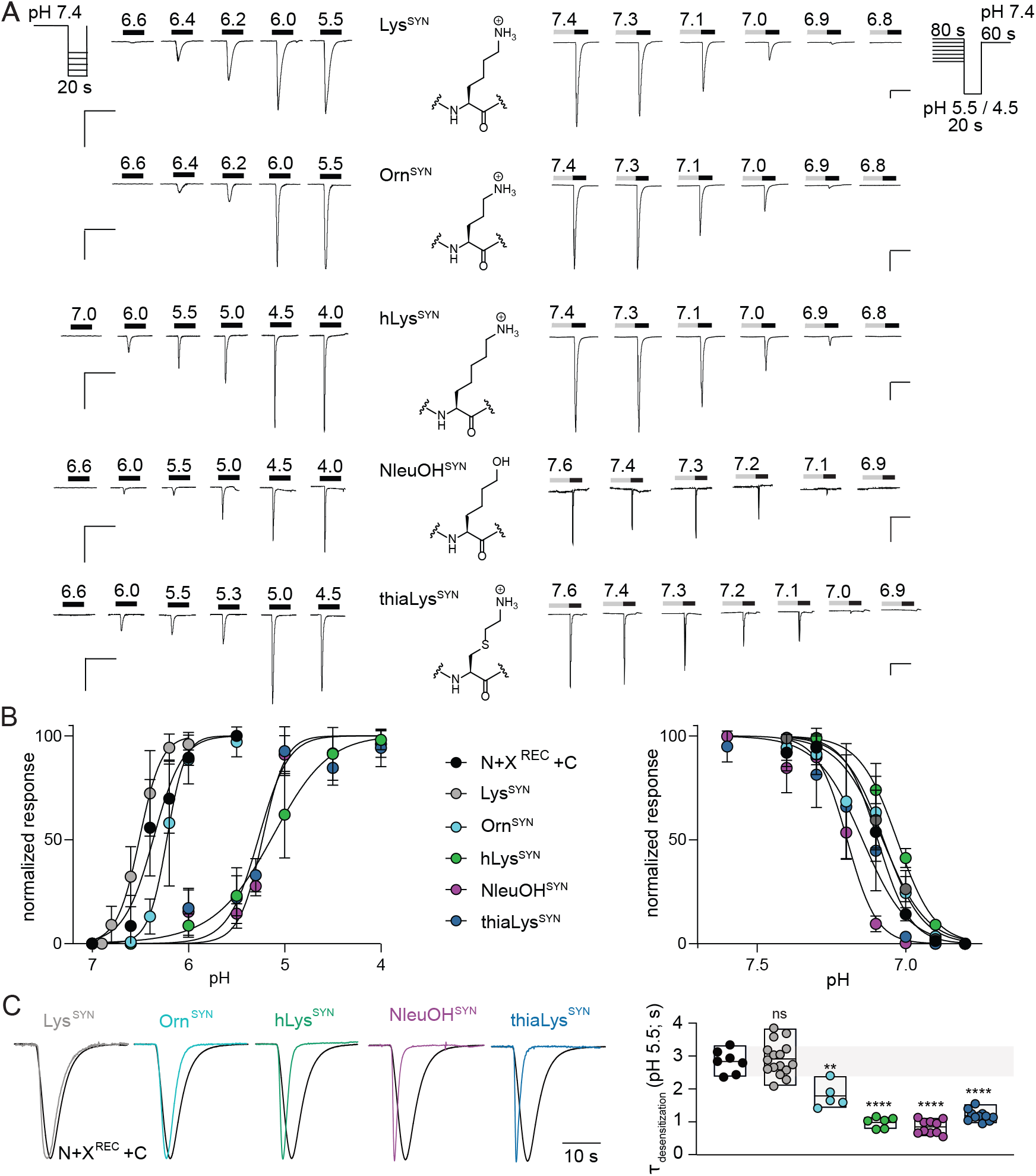
Functional characterization of mASIC1a channels trans-spliced with recombinant N and C terminal fragment and Pept X^SYN^. (A) Representative traces in the top panel depicts the pH activation profiles on the left and SSD profiles on the right with the chemical structure of the ncAA replacing Lys211 shown at the center. The buffer perfusion protocols are shown on the top left (activation) or top right (SSD) of the panels to depict the duration of each pH steps, and the 20 sec exposure to buffer with activating proton concentrations shown as thickened black bar on the current traces. Time period of exposure to desensitizing buffer conditions is shown in grey bars (last 30 sec of 80 sec). Scale bars: x, 60 sec; y, 0.5 μA. (B) The activation (left) and SSD (right) curves of pH response for all the spliced mASIC1a channels with four different ncAAs at the Lys211 position (n=6-10). (C) Comparison of kinetics of fast desensitization for all five synthetic peptide substitutions with the recombinant spliced construct of the mASIC1a channel, N+X^REC^+C^T214S^ variant (N+X^REC^+C). In left panel, N+X^REC^+C activation trace at pH_50_ is shown in black and all the traces (normalized to N+X^REC^+C trace) corresponding to other ncAAs has been shown in other colors. Quantitative evaluation of the desensitization time constants for each construct have been shown in the graph on the right panel. Scale bars: x, 10 sec. Each sphere represents individual data points (n=5-16 for each). The floating bars depict the mean along with minimum and maximum current margins, unpaired t-test to data from N+X^REC^+C, **p<0.005, ****p<0.0001.

We further evaluated fast desensitization kinetics (τ_desensitization_) at pH_50_ and pH 5.5 (Figure 4C, left panel and Supplemental Figure S3, respectively). Reassuringly, we found that the semisynthetic Lys-containing channels did not show significant differences in kinetics of fast desensitization compared with recombinant channels (N+X^REC^+C) (see Supplemental Table T2). By contrast, all ncAA analogs of Lys used here (Orn, hLys, thiaLys and NleuOH) led to a significant acceleration in τ_desensitization_ (Figure 4C, right panel, Supplemental Figure S3, right panel and Supplemental Table T2). Our data demonstrate that even the most subtle side chain manipulations at Lys211 drastically enhance the process of fast desensitization.

### Chloride binding affects channel dynamics

We turned to molecular modeling to determine how substituting position 211 with lysine analogs might impact the function of ASIC1a. In the trimeric crystal structure of chicken ASIC1 (cASIC1), the amine on the side chain equivalent to mASIC1a Lys211 (Lys212 by cASIC1 numbering) from the neighboring protomer coordinates the chloride ion across the interface (3.0Å). We created a simplified system based on PDB ID 5wku ^9^ that contained only a dimer and removed the TMD helices to reduce the number of atoms in the simulation. Then we used the Rosetta Relax Mover ^38^ using the Rosetta Scripts application ^39^ with restricted sampling only of loop 209-214 with either ASIC1a containing Lys, Orn, or hLys. The Rosetta total score is the sum of all residues in the simulation. Therefore, to probe the specific role of position 211, we sorted our results by looking at the Rosetta Energy Units (REU) for residue 211 (Figure 5A). Examining the lowest scoring structure from each simulation (Figure 5B) we find that the lowest scoring Lys and Orn both can still coordinate the chloride. However, hLys was no longer near the chloride ion, and instead interacted with the backbone of cysteine 361 in a neighboring loop (Figure 5B; green). Based on this result, we suspected that altering the lysine side chain length resulted in destabilizing chloride binding. Accordingly, we next investigated the REU for chloride (Figure 5C). In this analysis, we observed significant differences in the chloride REU for the different lysine analogs. Lys has the lowest average score and lowest overall score, (−2.2 and -3.4 respectively), followed by Orn (−2.1 and -3.1) and hLys was considerably worse (−1.5 and -2.2). These results suggest that neither Orn nor hLys can coordinate the chloride as well at Lys, and hLys is substantially worse than Orn. Examining the lowest scoring chloride models (Figure 5D) reveals that for Orn and Lys the sidechain rotamers are very similar to the lowest scoring 211 models (Figure 5B), but in the case of hLys the amine is now pointed towards the chloride. The score for position 211 in this model score is -2.1 REU and is about 3 REU less favorable than the alternative conformations of hLys (Figure 5B), which explains why the chloride coordinating structure was not frequently observed in the models (Figure 5B). Finally, we modeled hLys in the absence of chloride to determine if the side chain could enter the chloride pocket. However, surprisingly, an alternative hLys configuration where the side chain interacted with the backbone in a neighboring loop was observed (Figure 5E). Closer examination of these models reveals that the chloride pocket is more dynamic without the chloride present. We observed more dynamics in the loop near the chloride ion (Figure 5E; star) and we even observed models in which the disulfide bond that neighbors the Cl-ion adopts alternative rotamers. These dynamics were not observed in any of the structures where chloride was present. Collectively, these computational studies indicate that neither Orn nor hLys can coordinate Cl-as effectively as Lys, with hLys being significantly worse than Orn. Disruption of the Cl-ion binding site has been shown to accelerate kinetics of fast desensitization in functional experiments: both removing the charge at Lys211 through an Ala mutation and replacement of Cl-by larger anions accelerated mASIC1a fast desensitization kinetics.^12^ Given the role predicted for Lys211 in Cl-ion binding, we set out to assess the kinetics of fast desensitization in mASIC1a channels containing Lys or selected ncAA analogs of Lys in Pept X^SYN^ in buffers with different chloride concentrations. For the low-Cl-buffer, we substituted the majority of Cl-with the larger anion methanesulfonate, leaving a Cl-concentration of ∼18 mM, compared with ∼104 mM in regular ND96 (note that the Cl-concentration refers to the nominal concentration, i.e. before using HCl to adjust the pH of the buffer; actual concentrations are thus slightly higher). When activated at pH_50_, a reduction in Cl-led to a significantly accelerated τ_desensitization_ for Lys (Figure 5G, Supplemental Table T3). As expected, the desensitization kinetics of Orn and hLys were already faster than that of Lys in ND96 (Figure 4C). However, unlike in the case of Lys, a reduction in Cl-concentration did not accelerate the τ_desensitization_ further (Figure 5G, Supplemental Table T3). These findings are in line with the notion that the Lys analogs coordinate the Cl-ions less efficiently and are thus less affected by changes in Cl-concentrations.

**Figure 5:**
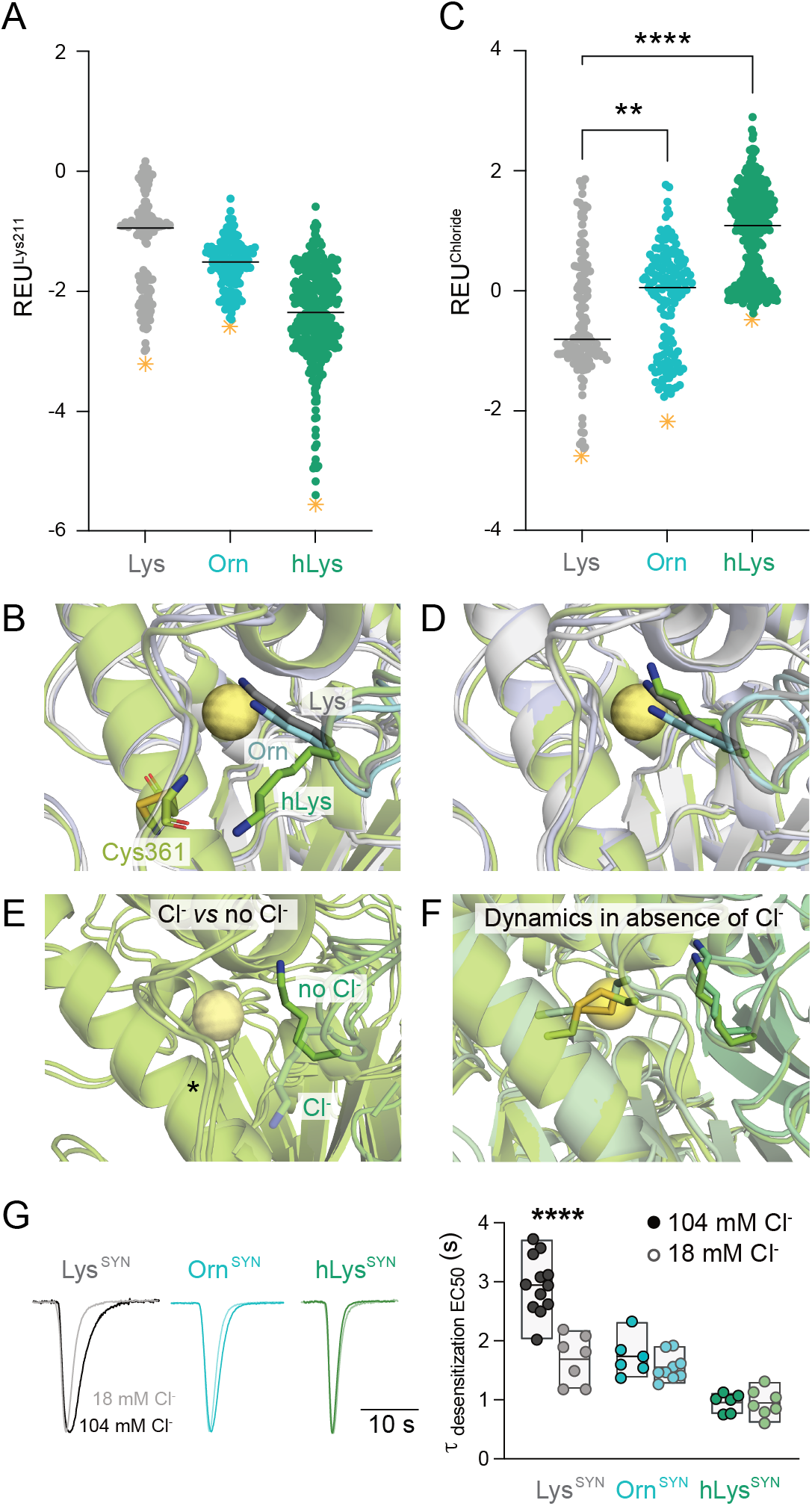
Effect of chloride binding on protein conformation and desensitization. (A) Rosetta energy for residue 211. (B) Representations of the lowest scoring models in A (Lys: grey, Orn; cyan hLys; green), indicated by the orange stars in panel A. (C) Rosetta energy for the chloride ion in the simulations. (D) Representations of the lowest scoring structures for Orn, Lys, and hLys indicated by the orange stars in panel C. (E) Model of the lowest scoring structure for hLys when Cl-is not present, the lowest scoring hLys structure with chloride present is shown in transparent mode. The star indicates the dynamic loop. (F) Overlay of two structures with different disulfide conformations to emphasize the dynamics when chloride is absent (note that original location of chloride is shown for reference). (G) Comparison of kinetics of fast desensitization for Lys^SYN^, Orn^SYN^ or hLys^SYN^ in functional experiments using either ND96 (∼104 mM Cl-) or low-chloride buffer (∼18 mM Cl-). In left panel, activation trace at pH_50_ in ND96 is shown in dark while activations in low-chloride buffer are shown in lighter colors. The panel on the right shows the quantitative comparison of the desensitization time constants. Each sphere represents an individual oocyte (n=6-12). The floating bars depict the mean along with minimum and maximum current margins, unpaired, ****p<0.0001. Scale bar: 10 sec.

## DISCUSSION

Structure-function studies have proven invaluable to advance our understanding of complex membrane proteins such as ion channels. However, progress regarding the contribution of evolutionarily highly conserved and/or functionally crucial side chains is frequently stymied by the limited chemical space afforded by conventional site-directed mutagenesis.^19^. Here we overcome this constraint by employing tPTS to incorporate a range of subtle Lys analogs into position Lys211 in mASIC1a. Our results underscore an important role of this side chain in both channel activation and fast desensitization.

### The role of Lys211 in mASIC1a activation and fast desensitization

Previous studies have suggested for Lys211 to be involved in ASIC activation via a H-bonding network with the backbones of Leu351 and Asp355 from the neighboring subunit. This was based on the observations that the distance between Lys211 and Leu351 is decreased in the open state compared with the resting state of cASIC1 ^7,9,11^ and that deleting Lys211 or covalently bridging the adjacent palm and thumb ECD subdomains by disulfide locking causes a right-shift in proton sensitivity.^11,18^ Here, we find that altering the pKa and altering side-chain geometry or removing the charge of Lys211 (thiaLys and NleuOH, respectively) is roughly as detrimental to mASIC1a activation, as is extending the side chain length by one carbon (hLys). By contrast, a shorter side chain is relatively well tolerated, as Orn incorporation in position 211 does not affect the proton sensitivity of activation.

On the other hand, we observed significantly accelerated desensitization kinetics with all Lys analogs incorporated at position 211. This suggests that the H-bonding network involved in activation is slightly more permissible towards minor disruptions than the process of fast desensitization. This notion is supported by the observation that the conventional Lys211Arg mutation displays near WT-like activation properties despite an increase in fast desensitization kinetics.^11^

Our data show that the process of fast desensitization is sensitive to even the most subtle of manipulations. The roughly 3-fold acceleration in τ_desensitization_ reported previously for the conventional Lys211Ala mutation ^12^ is similar to the change we observe for the longer Lys analog hLys or those with altered or removed charge (NleuOH and thiaLys). Our data suggests that the accelerated τ_desensitization_ are the result of disrupted coordination of Cl-ions that have been proposed to bind ASICs in a state-dependent manner, because they have been observed in open and desensitized structures of cASIC1 ^6,7^ but not in a resting state at low pH ^9,14^. In simulations, both removal of the Cl-ion or substitution of Lys by Orn or hLys destabilized the binding pocket. Functional experiments corroborate these findings and show that substitutions of Lys211 or a reduction of Cl-ions is sufficient but non-additive to accelerate-τ_desensitization_. The physiological relevance of Cl-ions in fast desensitization of various ASICs (incl heterologously expressed ASIC1a, 2a and 3 channels) is underscored by the finding that they affect ASIC currents in native dorsal root ganglia preparations and have been shown to possibly play a role in acid-induced neuronal death.^12,13^

Our finding that even minor alterations of side chain length and/or charge significantly affect fast desensitization kinetics via the disrupted coordination of Cl-ions is supported by the observation that proteinaceous ion binding sites tend to be highly sensitive to mutational disruptions, while H-bonding patterns or charge-charge interactions are often more resilient to small perturbations.^40,41^ Similar studies demonstrated that longer or shorter Lys analogs are detrimental to ATP recognition in P2X receptors, substrate recognition in Src homology 2 domains, and catalysis by histone lysine methyltransferases and ubiquitin chain-forming enzymes. ^29,42-44^

In summary, we conclude that Lys211 plays an important role in both ASIC activation and fast desensitization. This notion is supported by the finding that mutations at the adjacent G212 also affect both processes ^15^, thus underlining the broad functional relevance of this protein domain.

### Optimization attempts

Our initial screening of a range of split intein pairs and two different splice site combinations pointed towards the *Cfa*DnaE - *Ssp*DnaB^M86^ split intein pair as the most promising candidate. The large span of observed phenotypes, ranging from efficient channel reconstitution (*Cfa*DnaE - *Ssp*DnaB^M86^) to intermediate levels (e.g. gp41-1 – *Ssp*DnaX, *Npu*DnaE - *Ssp*DnaB^M86^ and NrdJ1 - *Ssp*DnaB^M86^) or no apparent reconstitution (e.g. *Mja*KlbA – *Ssp*DnaB^M86^, gp41-1 – CatInt and *Mja*KlbA – VidaL) clearly emphasizes the advantages of testing different split intein combinations for a given splice site.^21,29,33-36^ Notably, and despite its highly asymmetric split ^29^, *Ssp*DnaB^M86^ appears to be highly suitable as intein B: three out of four split intein pair combinations with *Ssp*DnaB^M86^ as intein B yielded measurable currents. By contrast, only one out of four designs with *Ssp*DnaX as intein B, and neither of the two designs with VidaL as intein B (Figure 2B) yielded functional mASIC1a constructs. This is not unexpected because the *Ssp*DnaB^M86^ split intein has undergone optimization from the wild-type *Ssp*DnaB intein to display greater tolerance toward extein sequences that deviate from its natural sequence context.^45,46^ Similarly, the *Cfa*DnaE split intein acquired its favorable splicing properties and stability from a consensus sequence-based design of related DnaE proteins.^47^

Somewhat surprisingly, our subsequent optimization attempts failed to improve apparent splicing yields and/or efficiency: neither physically linking the N-TMD/C-TMD fragments nor optimizing the extein sequence context led to a considerable increase in current amplitudes over those observed for the originally tested constructs based on the *Cfa*DnaE - *Ssp*DnaB^M86^ split intein pair. We conclude that physical proximity is not likely a major limiting factor and speculate that either sequence optimization was detrimental to channel function or the sequence context might be less important for non-cytosolic splicing events. The latter notion is supported by the fact that extein sequence optimization did not generally improve apparent splicing yields in any of the tested constructs. It is also possible that some of the mutations introduced through the optimization process may be detrimental to ASIC1a function. However, it should be noted that the current amplitudes achieved here with the *Cfa*DnaE - *Ssp*DnaB^M86^ split intein pair (1 to 6 μA) were significantly greater than those observed previously for tPTS in extracellular domains of structurally related P2X receptors (typically less than 500 nA, see ^29^). The resulting high signal-to-noise ratio therefore still enabled us to conduct a detailed structure-function analysis of the ncAA-containing mASIC1a variants.

### Limitations of our study

We observed minor differences in the pH sensitivity of activation and SSD of the full-length channel containing the Thr214Ser mutant compared with the N+X^REC^+C combination, which also contains Thr214Ser to enable splicing (Supplemental Figure S2). Both constructs should thus comprise the exact same amino acid sequence, the only difference being that the N+X^REC^+C combination must undergo tPTS to assemble into full-length channels. The observed functional difference could indicate the existence of a subpopulation of channels that contain not fully spliced protomers. We have carefully excluded the possibility of just the N-TMD and C-TMD fragments assembling into functional channels. It is conceivable, however, that in the presence of one or two fully spliced mASIC1a protomers, one or two non-or partially spliced N-TMD and/or C-TMD fragments could assemble to form part of a trimeric channel with altered functional properties. However, this remains speculative and further investigation of this possibility is hampered by the fact that the overall splicing efficiency is exceedingly difficult to quantify, but estimated to be less than 3% (see ^29^).

While *Xenopus laevis* oocytes are highly amenable to injection of the synthetic peptides required for tPTS^28-30^, they do not provide the intricate cellular environment afforded by neuronal tissue where ASICs are typically expressed. Additionally, their large physical size results in relatively slow solution exchange. This means we are limited to the study of steady-state measurements, as well as those of the relatively slow kinetics of fast desensitization and we cannot rule out that the τ_desensitization_ values reported for the ncAA analogs of Lys may be underestimated due to limited temporal resolution of TEVC. Lastly, we cannot accurately determine channel activation kinetics and are thus unable to assess the effect of ncAA introduction at Lys211 on the activation kinetics of mASIC1a.

## Conclusion

In this study we demonstrate how subtle analogs of a conserved Lys side chain in mASIC1a can be employed to decipher its contribution to both channel activation and fast desensitization with atomic precision. Our work highlights the potential of tPTS-based approaches for detailed structure-function studies in complex membrane proteins such as ligand-gated ion channels that can be studied with high signal-to-noise methods, making it ideally suited for use with electrophysiology.

## MATERIALS AND METHODS

A detailed description of materials and methods is included in the STAR methods section. In brief, mASIC1a variants were generated by site-directed mutagenesis on wildtype mASIC1a clone.^48^ The split intein-containing clones were obtained from a commercial source (Twist Biosciences). Recombinant, as well as semisynthetic mASIC1a constructs were expressed in *Xenopus laevis* oocytes, and proton-induced currents were measured using TEVC. First, a screen for recombinant reconstitution of mASIC1a channels using 11 different split intein combinations was performed. This was followed by different approaches to improve splicing yields (as measured by maximal current amplitudes). We finally conducted a detailed analysis of proton sensitivity of activation, steady-state desensitization properties of both recombinant and semisynthetic channels, including comparison of fast desensitization kinetics. We complemented our experimental work by molecular simulations using Rosetta applications to describe the conformational dynamics of the mASIC1a chloride binding pocket.

## Supporting information

STAR methods

## ACKNOWLEGEMENTS

We would like to thank Drs Valeria Kalienkova and Timothy Lynagh for suggestions regarding the experimental design and obtaining preliminary data, respectively. We would also like to thank Dr Mette H. Poulsen and Hendrik Harms for helpful comments on the manuscript. Our work was supported by the Lundbeck Foundation (R303-2018-2900 to IG) and the Independent Research Fund Denmark (9039-00335B to SAP).

## AUTHOR CONTRIBUTIONS

D.S., I.G., K.K.K. and S.A.P. designed the research. D.S., I.G., S.A.H., S.Y.O. and G.R.U. performed the experiments and analyzed the data. I.G. and G.J.V.D.H.V.N. designed the synthesis of the peptides. J.S.H. conducted the computational work. D.S., I.G., S.A.H., J.S.H. and S.A.P. wrote the manuscript with input from all authors.

## REFERENCES

1. Vullo, S., and Kellenberger, S. (2020). A molecular view of the function and pharmacology of acid-sensing ion channels. Pharmacol Res 154, 104166. 10.1016/j.phrs.2019.02.005.

2. Wemmie, J.A., Taugher, R.J., and Kreple, C.J. (2013). Acid-sensing ion channels in pain and disease. Nat Rev Neurosci 14, 461–471. 10.1038/nrn3529.

3. Boscardin, E., Alijevic, O., Hummler, E., Frateschi, S., and Kellenberger, S. (2016). The function and regulation of acid-sensing ion channels (ASICs) and the epithelial Na(+) channel (ENaC): IUPHAR Review 19. Br J Pharmacol 173, 2671–2701. 10.1111/bph.13533.

4. Kellenberger, S., and Schild, L. (2015). International Union of Basic and Clinical Pharmacology. XCI. structure, function, and pharmacology of acid-sensing ion channels and the epithelial Na+ channel. Pharmacol Rev 67, 1–35. 10.1124/pr.114.009225.

5. Heusser, S.A., and Pless, S.A. (2021). Acid-sensing ion channels as potential therapeutic targets. Trends Pharmacol Sci 42, 1035–1050. 10.1016/j.tips.2021.09.008.

6. Jasti, J., Furukawa, H., Gonzales, E.B., and Gouaux, E. (2007). Structure of acid-sensing ion channel 1 at 1.9 A resolution and low pH. Nature 449, 316–323. 10.1038/nature06163.

7. Baconguis, I., Bohlen, C.J., Goehring, A., Julius, D., and Gouaux, E. (2014). X-ray structure of acid-sensing ion channel 1-snake toxin complex reveals open state of a Na(+)-selective channel. Cell 156, 717–729. 10.1016/j.cell.2014.01.011.

8. Baconguis, I., and Gouaux, E. (2012). Structural plasticity and dynamic selectivity of acid-sensing ion channel-spider toxin complexes. Nature 489, 400-405. nature11375 [pii]10.1038/nature11375.

9. Yoder, N., Yoshioka, C., and Gouaux, E. (2018). Gating mechanisms of acid-sensing ion channels. Nature 555, 397–401. 10.1038/nature25782.

10. Grunder, S., and Pusch, M. (2015). Biophysical properties of acid-sensing ion channels (ASICs). Neuropharmacology 94, 9–18. 10.1016/j.neuropharm.2014.12.016.

11. Lynagh, T., Mikhaleva, Y., Colding, J.M., Glover, J.C., and Pless, S.A. (2018). Acid-sensing ion channels emerged over 600 Mya and are conserved throughout the deuterostomes. Proc Natl Acad Sci U S A 115, 8430–8435. 10.1073/pnas.1806614115.

12. Kusama, N., Harding, A.M., and Benson, C.J. (2010). Extracellular chloride modulates the desensitization kinetics of acid-sensing ion channel 1a (ASIC1a). J Biol Chem 285, 17425–17431. 10.1074/jbc.M109.091561.

13. Kusama, N., Gautam, M., Harding, A.M., Snyder, P.M., and Benson, C.J. (2013). Acid-sensing ion channels (ASICs) are differentially modulated by anions dependent on their subunit composition. Am J Physiol Cell Physiol 304, C89–101. 10.1152/ajpcell.00216.2012.

14. Yoder, N., and Gouaux, E. (2018). Divalent cation and chloride ion sites of chicken acid sensing ion channel 1a elucidated by x-ray crystallography. PLoS One 13, e0202134. 10.1371/journal.pone.0202134.

15. Bignucolo, O., Vullo, S., Ambrosio, N., Gautschi, I., and Kellenberger, S. (2020). Structural and Functional Analysis of Gly212 Mutants Reveals the Importance of Intersubunit Interactions in ASIC1a Channel Function. Front Mol Biosci 7, 58. 10.3389/fmolb.2020.00058.

16. Collier, D.M., and Snyder, P.M. (2009). Extracellular protons regulate human ENaC by modulating Na+ self-inhibition. J Biol Chem 284, 792–798. 10.1074/jbc.M806954200.

17. Collier, D.M., and Snyder, P.M. (2011). Identification of epithelial Na+ channel (ENaC) intersubunit Cl-inhibitory residues suggests a trimeric alpha gamma beta channel architecture. J Biol Chem 286, 6027–6032. 10.1074/jbc.M110.198127.

18. Gwiazda, K., Bonifacio, G., Vullo, S., and Kellenberger, S. (2015). Extracellular Subunit Interactions Control Transitions between Functional States of Acid-sensing Ion Channel 1a. J Biol Chem 290, 17956–17966. 10.1074/jbc.M115.641688.

19. Braun, N., Sheikh, Z.P., and Pless, S.A. (2020). The current chemical biology tool box for studying ion channels. J Physiol 598, 4455–4471. 10.1113/JP276695.

20. Plested, A.J.R., and Poulsen, M.H. (2021). Crosslinking glutamate receptor ion channels. Methods Enzymol 652, 161–192. 10.1016/bs.mie.2021.03.005.

21. Burton, A.J., Haugbro, M., Parisi, E., and Muir, T.W. (2020). Live-cell protein engineering with an ultra-short split intein. Proc Natl Acad Sci U S A 117, 12041–12049. 10.1073/pnas.2003613117.

22. Giriat, I., and Muir, T.W. (2003). Protein semi-synthesis in living cells. J Am Chem Soc 125, 7180–7181. 10.1021/ja034736i.

23. Ray, D.M., Flood, J.R., and David, Y. (2022). Harnessing Split-Inteins as a Tool for the Selective Modification of Surface Receptors in Live Cells. Chembiochem, e202200487. 10.1002/cbic.202200487.

24. Bhagawati, M., Hoffmann, S., Hoffgen, K.S., Piehler, J., Busch, K.B., and Mootz, H.D. (2020). In Cellulo Protein Semi-Synthesis from Endogenous and Exogenous Fragments Using the Ultra-Fast Split Gp41-1 Intein. Angew Chem Int Ed Engl. 10.1002/anie.202006822.

25. Thompson, R.E., and Muir, T.W. (2020). Chemoenzymatic Semisynthesis of Proteins. Chem Rev 120, 3051–3126. 10.1021/acs.chemrev.9b00450.

26. Lockless, S.W., and Muir, T.W. (2009). Traceless protein splicing utilizing evolved split inteins. Proc Natl Acad Sci U S A 106, 10999–11004. 10.1073/pnas.0902964106.

27. Shah, N.H., and Muir, T.W. (2014). Inteins: Nature’s Gift to Protein Chemists. Chem Sci 5, 446–461. 10.1039/C3SC52951G.

28. Sarkar, D., Harms, H., Galleano, I., Sheikh, Z.P., and Pless, S.A. (2021). Ion channel engineering using protein trans-splicing. Methods Enzymol 654, 19–48. 10.1016/bs.mie.2021.01.028.

29. Khoo, K.K., Galleano, I., Gasparri, F., Wieneke, R., Harms, H., Poulsen, M.H., Chua, H.C., Wulf, M., Tampe, R., and Pless, S.A. (2020). Chemical modification of proteins by insertion of synthetic peptides using tandem protein trans-splicing. Nat Commun 11, 2284. 10.1038/s41467-020-16208-6.

30. Galleano, I., Harms, H., Choudhury, K., Khoo, K., Delemotte, L., and Pless, S.A. (2021). Functional cross-talk between phosphorylation and disease-causing mutations in the cardiac sodium channel Na(v)1.5. Proc Natl Acad Sci U S A 118. 10.1073/pnas.2025320118.

31. Aranko, A.S., Oeemig, J.S., Zhou, D., Kajander, T., Wlodawer, A., and Iwai, H. (2014). Structure-based engineering and comparison of novel split inteins for protein ligation. Mol Biosyst 10, 1023–1034. 10.1039/c4mb00021h.

32. Aranko, A.S., Wlodawer, A., and Iwai, H. (2014). Nature’s recipe for splitting inteins. Protein Eng Des Sel 27, 263–271. 10.1093/protein/gzu028.

33. Pinto, F., Thornton, E.L., and Wang, B. (2020). An expanded library of orthogonal split inteins enables modular multi-peptide assemblies. Nat Commun 11, 1529. 10.1038/s41467-020-15272-2.

34. Zhang, X., Liu, X.Q., and Meng, Q. (2019). Engineered Ssp DnaX inteins for protein splicing with flanking proline residues. Saudi J Biol Sci 26, 854–859. 10.1016/j.sjbs.2017.07.010.

35. Iwai, H., Zuger, S., Jin, J., and Tam, P.H. (2006). Highly efficient protein trans-splicing by a naturally split DnaE intein from Nostoc punctiforme. FEBS Lett 580, 1853–1858. 10.1016/j.febslet.2006.02.045.

36. Stevens, A.J., Sekar, G., Gramespacher, J.A., Cowburn, D., and Muir, T.W. (2018). An Atypical Mechanism of Split Intein Molecular Recognition and Folding. J Am Chem Soc 140, 11791–11799. 10.1021/jacs.8b07334.

37. Gloss, L.M., and Kirsch, J.F. (1995). Decreasing the basicity of the active site base, Lys-258, of Escherichia coli aspartate aminotransferase by replacement with gamma-thialysine. Biochemistry 34, 3990–3998. 10.1021/bi00012a017.

38. Conway, P., Tyka, M.D., DiMaio, F., Konerding, D.E., and Baker, D. (2014). Relaxation of backbone bond geometry improves protein energy landscape modeling. Protein Sci 23, 47–55. 10.1002/pro.2389.

39. Fleishman, S.J., Leaver-Fay, A., Corn, J.E., Strauch, E.M., Khare, S.D., Koga, N., Ashworth, J., Murphy, P., Richter, F., Lemmon, G., et al. (2011). RosettaScripts: a scripting language interface to the Rosetta macromolecular modeling suite. PLoS One 6, e20161. 10.1371/journal.pone.0020161.

40. Hille, B. (2001). Ionic channels of excitable membranes., Third Edition (Sinauer Associates Inc.).

41. Zheng, J., and Trudeau, M.C. (2015). Handbook of Ion Channels, First Ed. Edition (CRC Press). 10.1201/b18027.

42. Temimi, A., Reddy, Y.V., White, P.B., Guo, H., Qian, P., and Mecinovic, J. (2017). Lysine Possesses the Optimal Chain Length for Histone Lysine Methyltransferase Catalysis. Sci Rep 7, 16148. 10.1038/s41598-017-16128-4.

43. Liwocha, J., Krist, D.T., van der Heden van Noort, G.J., Hansen, F.M., Truong, V.H., Karayel, O., Purser, N., Houston, D., Burton, N., Bostock, M.J., et al. (2021). Linkage-specific ubiquitin chain formation depends on a lysine hydrocarbon ruler. Nat Chem Biol 17, 272–279. 10.1038/s41589-020-00696-0.

44. Virdee, S., Macmillan, D., and Waksman, G. (2010). Semisynthetic Src SH2 domains demonstrate altered phosphopeptide specificity induced by incorporation of unnatural lysine derivatives. Chem Biol 17, 274–284. 10.1016/j.chembiol.2010.01.015.

45. Appleby, J.H., Zhou, K., Volkmann, G., and Liu, X.Q. (2009). Novel split intein for trans-splicing synthetic peptide onto C terminus of protein. J Biol Chem 284, 6194–6199. 10.1074/jbc.M805474200.

46. Appleby-Tagoe, J.H., Thiel, I.V., Wang, Y., Wang, Y., Mootz, H.D., and Liu, X.Q. (2011). Highly efficient and more general cis- and trans-splicing inteins through sequential directed evolution. J Biol Chem 286, 34440–34447. 10.1074/jbc.M111.277350.

47. Stevens, A.J., Brown, Z.Z., Shah, N.H., Sekar, G., Cowburn, D., and Muir, T.W. (2016). Design of a Split Intein with Exceptional Protein Splicing Activity. J Am Chem Soc 138, 2162–2165. 10.1021/jacs.5b13528.

48. Lynagh, T., Flood, E., Boiteux, C., Wulf, M., Komnatnyy, V.V., Colding, J.M., Allen, T.W., and Pless, S.A. (2017). A selectivity filter at the intracellular end of the acid-sensing ion channel pore. Elife 6. 10.7554/eLife.24630.

