## Supplemental Information for "Protein semisynthesis underscores role of a conserved lysine in activation and desensitization of acid-sensing ion channels"

[§] These authors contributed equally

* Co-corresponding author:

Stephan Pless

University of Copenhagen

**Table of contents**


| Section | Content | Page No. |
|---|---|---|
| 1 | Peptide Characterization | 03 |
| 2 | Supplementary Table T1: Data summary of functional analysis of recombinant mASIC1a constructs | 08 |
| 3 | Supplementary Table T2: Data summary of functional analysis of semisynthetic mASIC1a constructs | 08 |
| 4 | Supplementary Table T3: Data summary of functional analysis of desensitization analysis in low-chloride buffers | 08 |
| 5 | Supplementary Figure S1: Comparative splicing yields of 11 split intein pairs | 09 |
| 6 | Supplementary Figure S2: Functional properties of wildtype and Thr214Ser full-length channels and recombinant spliced ($N+X^{REC}+C$) channel. | 10 |
| 7 | Supplementary Figure S3: Desensitization kinetics at pH 5.5 | 11 |

### 1. Peptide Characterization

#### Int$^{N}$-B

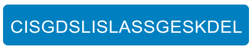


$[M+2H]^{2+}$ calc.: 1005.980 Da; $[M+3H]^{3+}$ calc.: 670.989 Da.

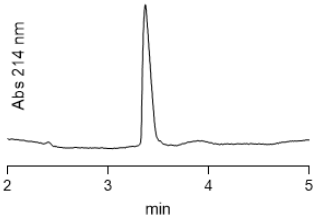


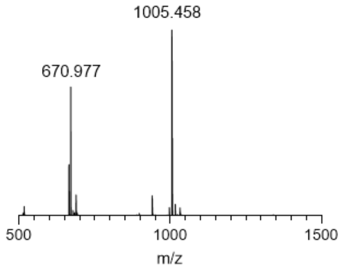


#### Int$^{C}$-A_thioester

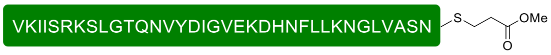


$[M+4H]^{4+}$ calc.: 994.041 Da; $[M+5H]^{5+}$ calc.: 795.434 Da; $[M+6H]^{6+}$ calc.: 663.030 Da.

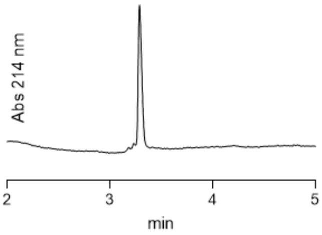


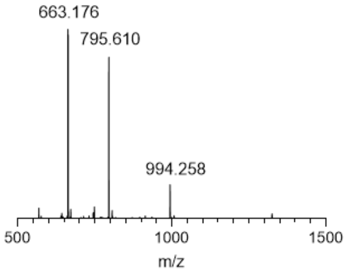


#### ASIC1a_Lys

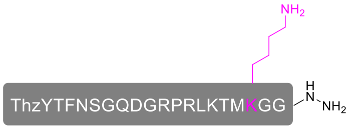


$[M+2H]^{2+}$ calc.: 1122.054 Da; $[M+3H]^{3+}$ calc.: 748.372 Da; $[M+4H]^{4+}$ calc.: 561.530 Da; $[M+5H]^{5+}$ calc.: 449.426 Da.

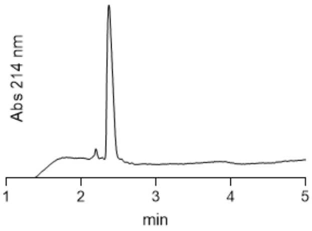


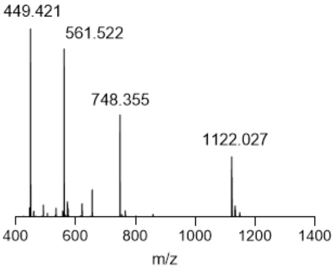

### ASIC1a_Orn

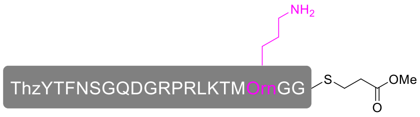


$[M+2H]^{2+}$ calc.: 1159.039 Da; $[M+3H]^{3+}$ calc.: 773.029 Da; $[M+4H]^{4+}$ calc.: 580.023 Da; $[M+5H]^{5+}$ calc.: 464.220 Da.

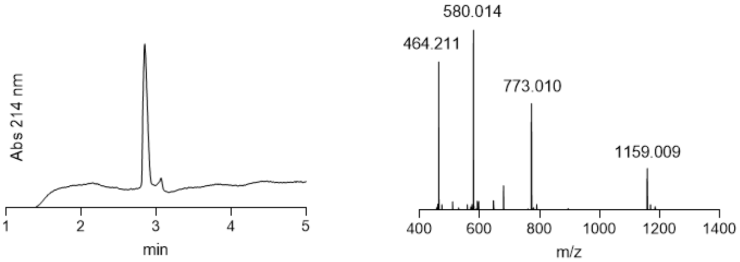


### ASIC1a_hLys

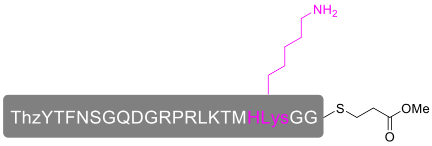


$[M+2H]^{2+}$ calc.: 1173.055 Da; $[M+3H]^{3+}$ calc.: 782.372 Da; $[M+4H]^{4+}$ calc.: 587.031 Da; $[M+5H]^{5+}$ calc.: 469.826 Da.

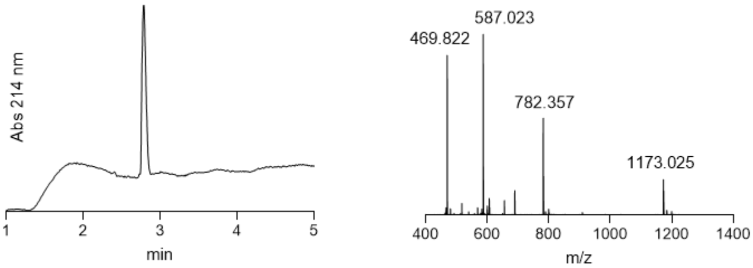


### ASIC1a_NleuOH

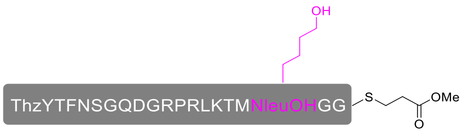


$[M+2H]^{2+}$ calc.: 1166.539 Da; $[M+3H]^{3+}$ calc.: 778.029 Da; $[M+4H]^{4+}$ calc.: 583.773 Da.

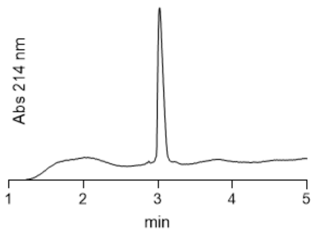

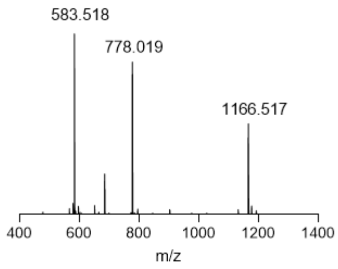


### ASIC1a_thiaLys

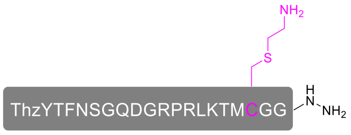


[M+2H]$^{2+}$ calc.: 1131.032 Da; [M+3H]$^{3+}$ calc.: 754.357 Da; [M+4H]$^{4+}$ calc.: 566.020 Da; [M+5H]$^{5+}$ calc.: 453.017 Da.

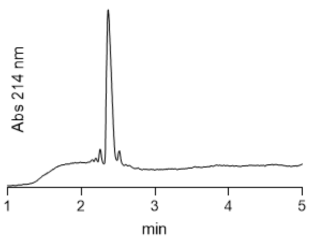

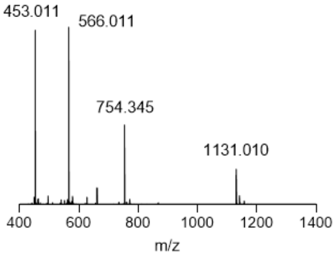


### PeptideX_Lys

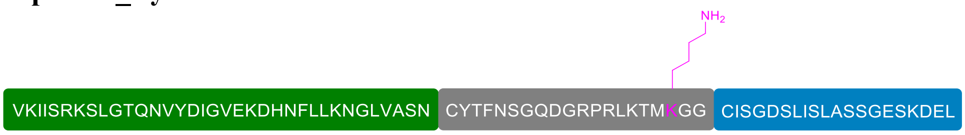


[M+6H]$^{6+}$ calc.: 1344.526 Da; [M+7H]$^{7+}$ calc.: 1152.595 Da; [M+8H]$^{8+}$ calc.: 1008.647 Da; [M+9H]$^{9+}$ calc.: 896.687 Da; [M+10H]$^{10+}$ calc.: 807.119 Da; [M+11H]$^{11+}$ calc.: 733.836 Da; [M+12H]$^{12+}$ calc.: 672.767 Da.
Deconvoluted mass [M+H]$^{+}$ calc.: 8064 Da.

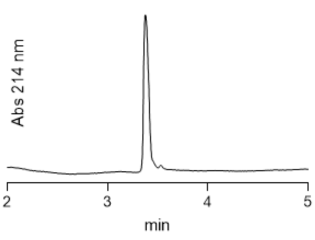

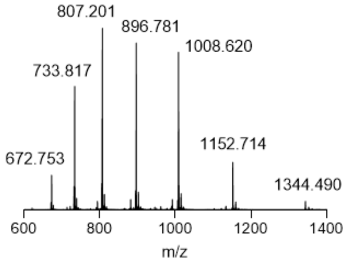

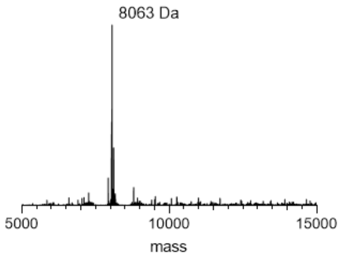


### PeptideX_Orn

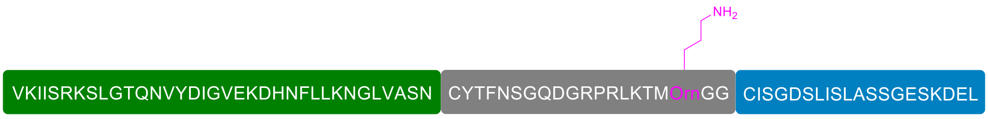


[M+6H]$^{6+}$ calc.: 1342.190 Da; [M+7H]$^{7+}$ calc.: 1150.593 Da; [M+8H]$^{8+}$ calc.: 1006.895 Da; [M+9H]$^{9+}$ calc.: 895.129 Da; [M+10H]$^{10+}$ calc.: 805.717 Da; [M+11H]$^{11+}$ calc.: 732.562 Da; [M+12H]$^{12+}$ calc.: 671.599 Da.
Deconvoluted mass [M+H]$^{+}$ calc.: 8050 Da.

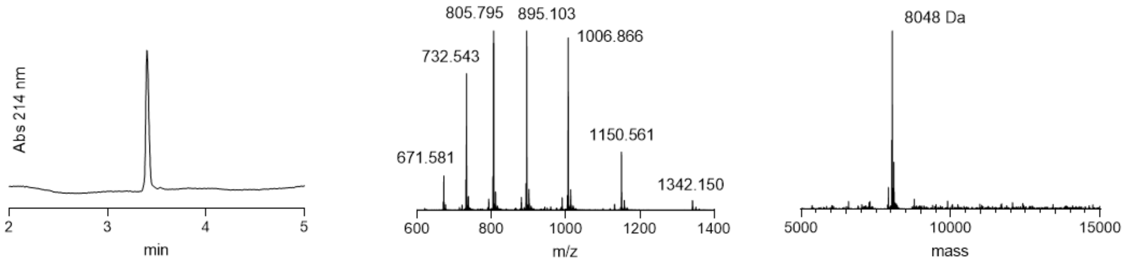


**PeptideX_hLys**

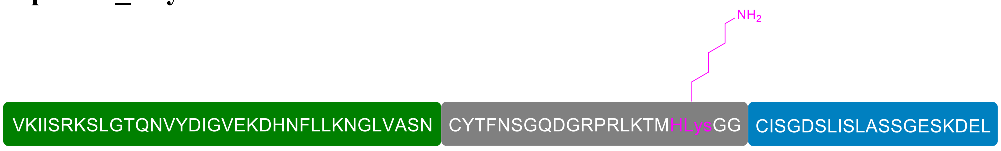


[M+6H]$^{6+}$ calc.: 1346.862 Da; [M+7H]$^{7+}$ calc.: 1154.597 Da; [M+8H]$^{8+}$ calc.: 1110.398 Da; [M+9H]$^{9+}$ calc.: 898.244 Da; [M+10H]$^{10+}$ calc.: 808.520 Da; [M+11H]$^{11+}$ calc.: 735.110 Da; [M+12H]$^{12+}$ calc.: 673.935 Da.
Deconvoluted mass [M+H]$^{+}$ calc.: 8078 Da.

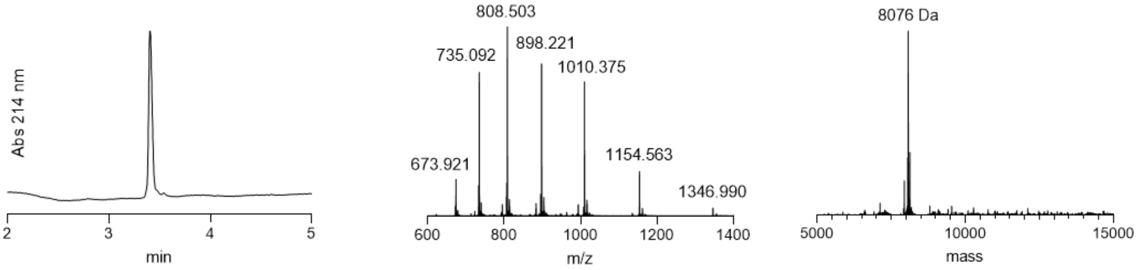


**PeptideX_NleuOH**

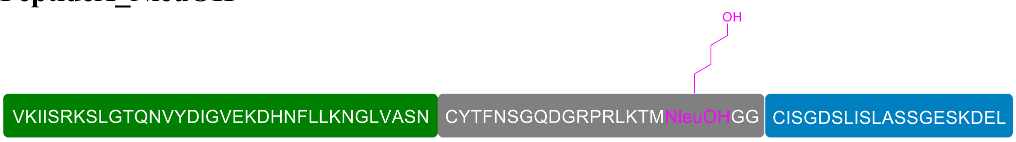


[M+6H]$^{6+}$ calc.: 1344.690 Da; [M+7H]$^{7+}$ calc.: 1152.736 Da; [M+8H]$^{8+}$ calc.: 1008.770 Da; [M+9H]$^{9+}$ calc.: 896.796 Da; [M+10H]$^{10+}$ calc.: 807.217 Da; [M+11H]$^{11+}$ calc.: 733.925 Da;.
Deconvoluted mass [M+H]$^{+}$ calc.: 8065 Da.

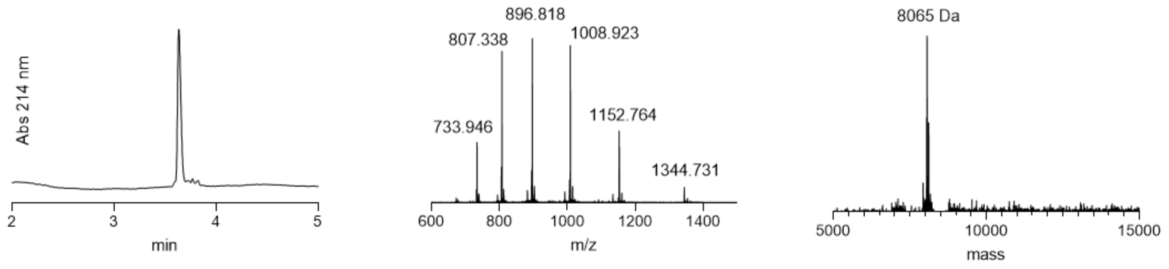


**PeptideX_thiaLys**

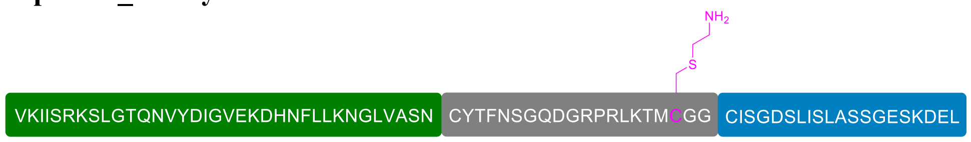


$[M+5H]^{5+}$ calc.: 1616.821 Da; $[M+6H]^{6+}$ calc.: 1347.519 Da; $[M+7H]^{7+}$ calc.: 1155.160 Da; $[M+8H]^{8+}$ calc.: 1010.891 Da; $[M+9H]^{9+}$ calc.: 898.682 Da; $[M+10H]^{10+}$ calc.: 808.914 Da; $[M+11H]^{11+}$ calc.: 735.468 Da; $[M+12H]^{12+}$ calc.: 674.263 Da;
Deconvoluted mass $[M+H]^{+}$ calc. : 8082 Da.

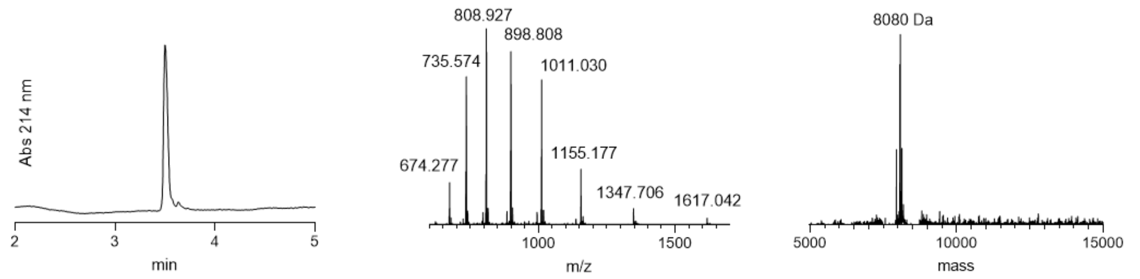

**2. Supplementary Table T1:** Data summary of functional analysis of recombinant mASIC1a constructs (mean ± SD, number of experimental repeats listed in Supplementary Figure S2B). Statistical analysis (unpaired t-test) of $pH_{50}$ values form mASIC1a Thr214Ser (full-length) and N+X$^{REC}$+C is performed with respect to mASIC1a WT (full-length) data. Significance levels: *** = $p < 0.001$; **** = $p < 0.0001$.

| mASIC1a variant | $pH_{50}$ (activation) | $pH_{50}$ (SSD) |
|---|---|---|
| WT (full-length) | 6.7 ± 0.1 | 7.2 ± 0.0 |
| Thr214Ser (full-length) | 6.5 ± 0.1 *** | 6.9 ± 0.0 **** |
| N+X$^{REC}$+C | 6.4 ± 0.1 **** | 7.1 ± 0.0 **** |

**3. Supplementary Table T2:** Data summary of functional analysis of semisynthetic mASIC1a constructs (mean ± SD, number of experimental repeats listed in Figure 4 in the main text). Statistical analysis (unpaired t-test) for semisynthetic channel data has been depicted in relation to N+X$^{REC}$+C channel data. Significance levels: * = $p < 0.05$; ** = $p < 0.01$; *** = $p < 0.001$; **** = $p < 0.0001$; ns = not significant.

| mASIC1a variant | $pH_{50}$ (activation) | $pH_{50}$ (SSD) | $\tau_{desensitization}$ (msec) at pH 5.5 | $\tau_{desensitization}$ (msec) at $pH_{50}$ |
|---|---|---|---|---|
| N+X$^{REC}$+C | 6.4 ± 0.1 | 7.1 ± 0.0 | 2910 ± 429 | 2744 ± 339 |
| K211Lys$^{SYN}$ | 6.4 ± 0.1$^{ns}$ | 7.1 ± 0.0$^{ns}$ | 2618 ± 533$^{ns}$ | 2818 ± 485$^{ns}$ |
| K211Orn$^{SYN}$ | 6.2 ± 0.1 * | 7.1 ± 0.1$^{ns}$ | 1726 ± 457 *** | 1735 ± 370 ** |
| K211hLys$^{SYN}$ | 5.2 ± 0.1**** | 7.0 ± 0.0 *** | 915 ± 270 **** | 954 ± 157 **** |
| K211NleuOH$^{SYN}$ | 5.2 ± 0.1**** | 7.2 ± 0.0 **** | 870 ± 300 **** | 822 ± 204 **** |
| K211thiaLys$^{SYN}$ | 5.3 ± 0.1**** | 7.1 ± 0.1$^{ns}$ | 1198 ± 270 **** | 1154 ± 166**** |

**4. Supplementary Table T3:** Data summary of desensitization data (mean ± SD, number of experimental repeats listed in Figure 5 in the main text). Statistical analysis (unpaired t-test) for desensitization kinetics at $pH_{50}$ in low-chloride buffer compared to $Ca^{2+}$ free ND96. Significance levels: **** = $p < 0.0001$; ns = not significant.

| mASIC1a variant | $\tau_{desensitization}$ (msec) at $pH_{50}$ | |
|---|---|---|
| | $Ca^{2+}$ free ND96 | Low-chloride buffer |
| K211Lys$^{SYN}$ | 2945± 491 | 1688 ± 413**** |
| K211Orn$^{SYN}$ | 1735 ± 332 | 1550 ± 225$^{ns}$ |
| K211hLys$^{SYN}$ | 952 ± 157 | 942 ± 233$^{ns}$ |

**5. Supplementary Figure S1:** Comparison of maximal currents yielded by the 11 split intein pairs tested here, recorded after 1-6 days post injection. N+C represents control experiments where only mRNAs for N-TMD and C-TMD fragments injected in *Xenopus laevis* oocytes. In case of the N+X$^{REC}$+C experiments, mRNA for the corresponding peptide X (X$^{REC}$) sequence was co-injected with the mRNAs for N-TMD and C-TMD fragments. Injection of native sequences is differentiated from the extein sequence optimized ones by (*) in these floating bar plots. Each floating bar represents the maximum and minimum values along with the average, collected over at least two batches of oocytes (n = 6-11, as indicted by numbers in brackets). Each data point is represented as a color-filled circle on the floating bars.

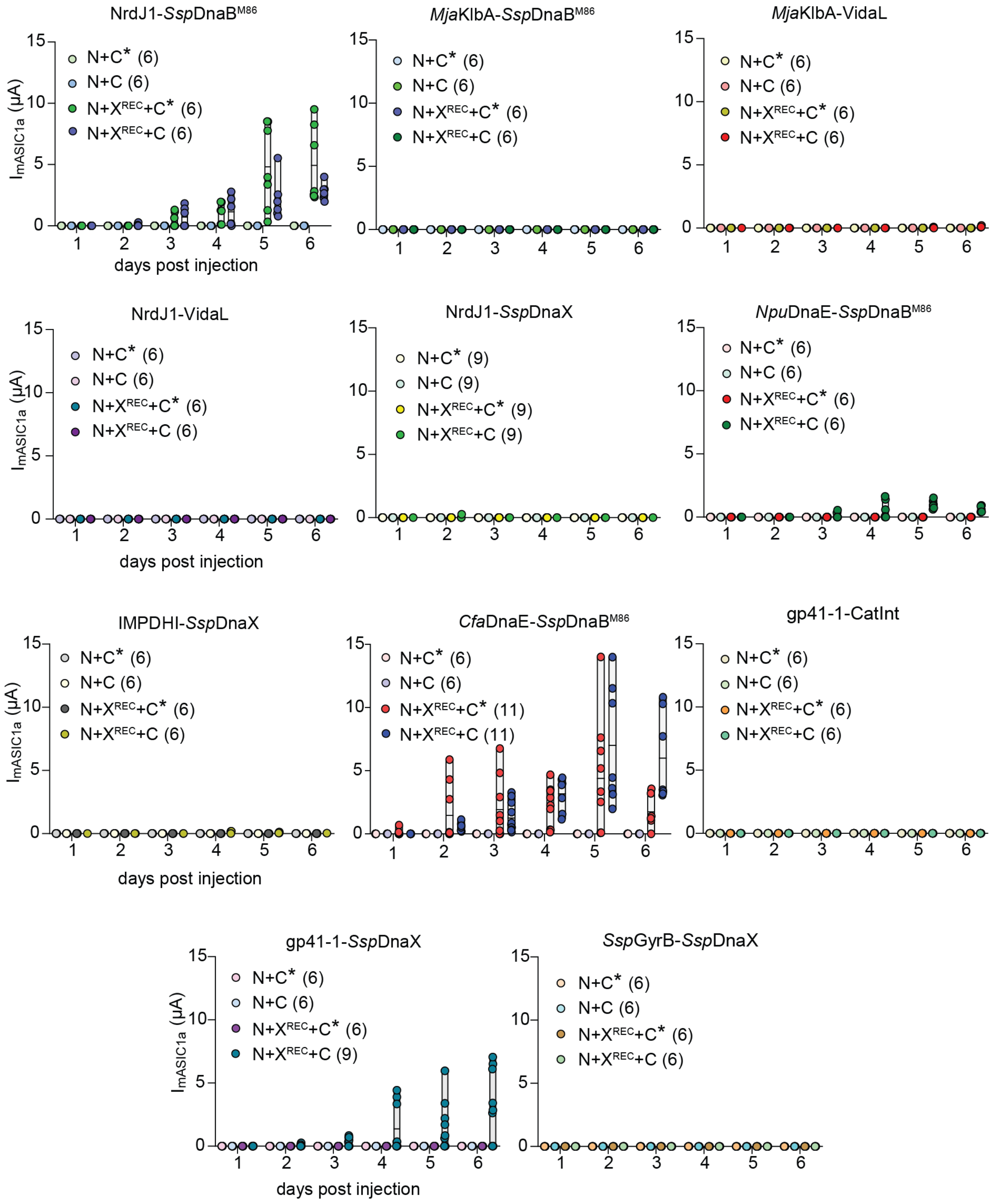

**6. Supplementary Figure S2:** Functional properties of full-length wildtype and full-length Thr214Ser mutant channels and recombinantly spliced (N+X$^{REC}$+C) channel. Scale bars: x, 10 s; y, 1 µA (unless otherwise mentioned). (A) Representative current traces of the full-length wildtype (mASIC1a$^{WT}$) and recombinantly spliced (N+X$^{REC}$+C) channels where the Thr at 214$^{th}$ position at the C-terminal fragment is kept unaltered (N+X$^{REC}$+C$^{T214}$) or replaced by Ser (N+X$^{REC}$+C$^{T214S}$). (B) Upper panel depicts representative traces for all the three constructs (left: activation, right: SSD, with protocols shown at the top). Bottom panel represents the pH sensitivity of activation and SSD. Each pH concentration-response plot is representative of the average of n = 5-11 individual data points averaged over at least two batches of oocytes. Error bars represent standard deviation.

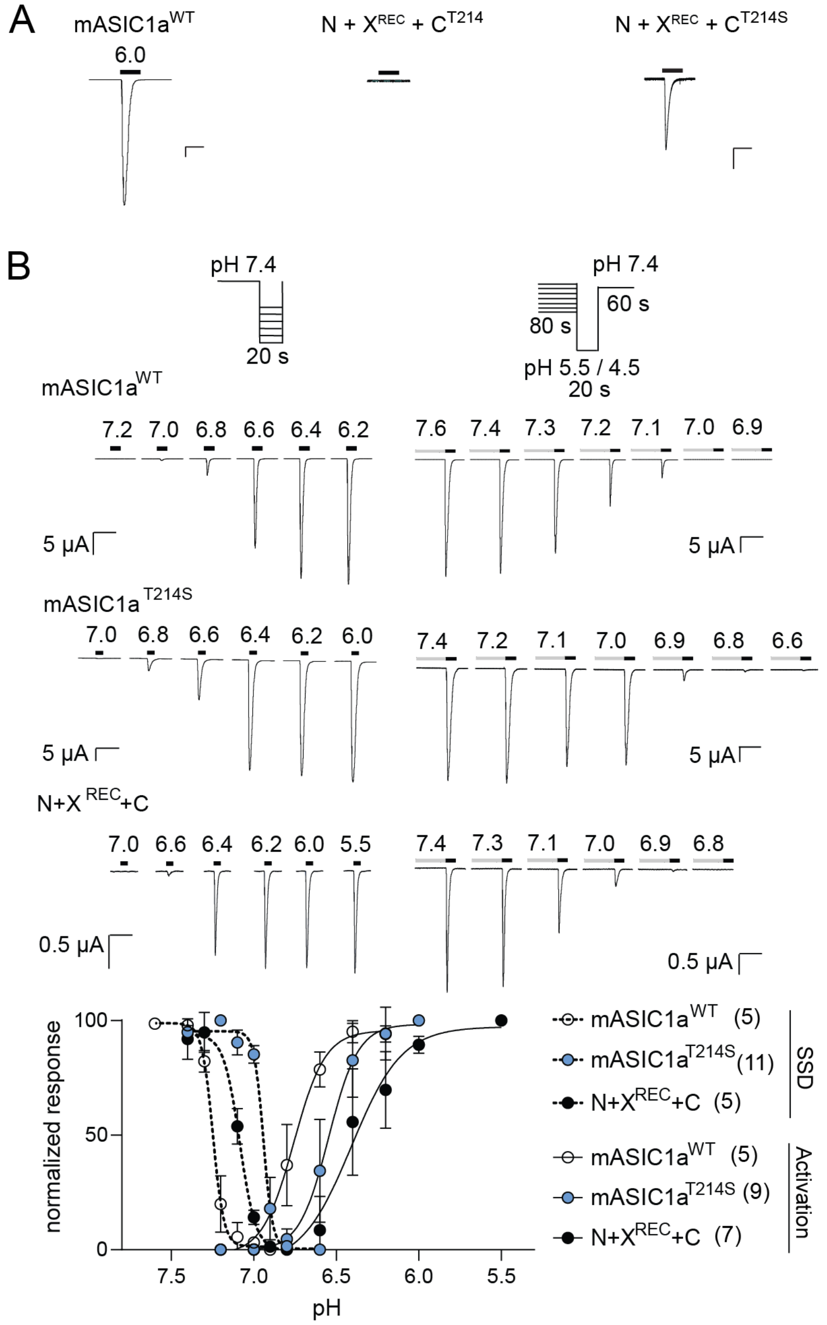

**7. Supplementary Figure S3:** Comparison of kinetics of fast desensitization for all five synthetic peptide substitutions with the recombinant spliced construct of the mASIC1a channel, N+X$^{REC}$+C T214S variant (N+X$^{REC}$+C). In left panel, N+X$^{REC}$+C activation trace at pH 5.5 is shown in black and all the traces (normalized to N+X$^{REC}$+C trace) corresponding to other ncAAs has been shown in other colors. Quantitative evaluation of the desensitization time constants for each construct have been shown in the graph on the right panel. Each sphere represents individual data points (n=6-8 for each). The floating bars depict the mean along with minimum and maximum current margins. Scale bars: x, 10 sec.

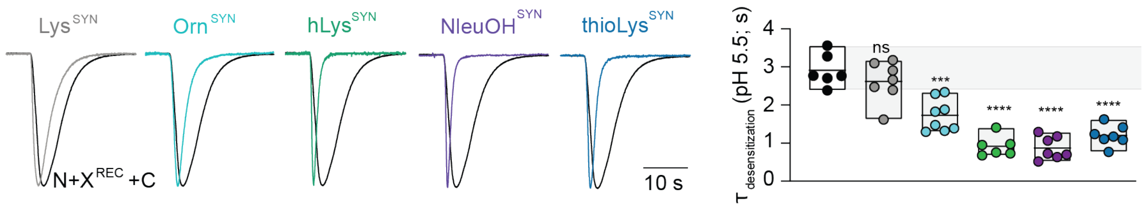
