## Supplementary material for "Protein semisynthesis underscores role of a conserved lysine in activation and desensitization of acid-sensing ion channels": STAR methods

### STAR Methods for

[§] These authors contributed equally

* Co-corresponding author:

Stephan Pless

University of Copenhagen

**Table of contents**

| Section | Content | Page No. |
|---|---|---|
| 1 | Resource Availability | 03 |
| 1A | Lead contact | 03 |
| 1B | Materials availability | 03 |
| 1C | Data and code availability | 03 |
| 2 | Experimental model and study participant details | 03 |
| 3 | Methods details | 03 |
| 3A | Peptide Synthesis - overview | 03 |
| 3B | Peptide Synthesis – details | 06 |
| 3C | Molecular biology | 08 |
| 3D | Preparation of *Xenopus laevis* oocytes and mRNA injections | 08 |
| 3E | Electrophysiological recordings | 09 |
| 3F | Molecular modeling | 09 |
| 4 | Quantification and statistical analysis | 09 |
| 5 | Key Resource Tables | 10 |
| 5A | Resource Table T1: Split intein linked mASIC1a constructs (Native sequence) | 11 |
| 5B | Resource Table T2: List of native and optimized extein sequences for split inteins used in this study. | 22 |
| 5C | Resource Table T3: Split intein linked mASIC1a constructs (Optimized sequence) | 23 |
| 6 | References | 30 |

### 1. RESOURCE AVAILABILITY

#### A. Lead contact

#### B. Materials availability

- Plasmids generated for this study are available upon request from the lead contact. Nucleotide and protein sequences of each construct generated in this study is detailed in Resource tables T1 and T3.
- There are restrictions to the availability of synthetic ASIC1a intein-peptides due to the amount that was originally produced for the project.

#### C. Data and code availability

- All raw data and analysis files related to the data reported in this paper will be shared by the lead contact upon request.
- This paper does not report original code. Details with respect to molecular modeling detailed in section 3F.
- Any additional information required to reanalyze the data reported in this paper is available from the lead contact upon request.

### 2. EXPERIMENTAL MODEL AND STUDY PARTICIPANT DETAILS

Female *Xenopus laevis* (the African clawed frog) were purchased from Xenopus 1, Corp. (Dexter, MI, USA). Stage 5 and 6 oocytes from adult female *Xenopus laevis* were surgically removed from frogs under anesthesia with 0.3% tricaine in accordance to License 2014-15-0201-00031, approved by the Danish Veterinary and Food Administration.

### 3. METHOD DETAILS

#### 3A. Peptide Synthesis - overview

SPPS was performed on a Syro II MultiSyntech Automated Peptide synthesizer under inert gas ($N_2$) application, using standard 9-fluorenylmethoxycarbonyl (Fmoc) based solid phase peptide chemistry using either pre-loaded Fmoc amino acid trityl, HMPA or Pyv resins (Rapp Polymere, Germany). All amino acids and dipeptide building blocks were double coupled in N-methyl-2-pyrrolidone (NMP) for 25 min using benzotriazol-1-yloxytripyrrolidinophosphonium hexafluorophosphate (PyBOP) (4 equiv) and *N,N*-diisopropylethylamine (DiPEA) (8 equiv) as coupling reagents. Fmoc removal was achieved with a solution of piperidine in NMP (20:80, v/v, 2x2 and 1x5 min).

Unless otherwise stated, the amino acids used for solid-phase peptide synthesis were Fmoc-Ala-OH, Fmoc-Cys(Trt)-OH, Fmoc-Phe-OH, Fmoc-Gly-OH, Fmoc-Ile-OH, Fmoc-Lys(Boc)-OH, Fmoc-Leu-OH, Fmoc-Pro-OH, Fmoc-His(Trt)- OH, Fmoc-Asn(Trt)-OH, Fmoc-Gln(Trt)-OH, Fmoc-Arg(Pbf)-OH, Fmoc-Ser(*t*Bu)-OH, Fmoc-Thr(*t*Bu)-OH, Fmoc-Tyr(*t*Bu)-OH, Fmoc-Asp(O*t*Bu)-OH, Fmoc-Glu(O*t*Bu)-OH, Fmoc-Met-OH, Fmoc-Val-OH, and Fmoc-Thz-OH.

**Cleavage from the resin and deprotection**

The polypeptide sequence was cleaved from the resin and deprotected by treatment with trifluoroacetic acid/triisopropylsilane/3,6-dioxa-1,8-octanedithiol (TFA/TIPS/DODT, 94:3:3, v/v/v) for 90 min followed by precipitation in cold diethylether ($Et_2O$). If the peptide sequence comprised methionine residues, the cleavage and deprotection was performed with an alternative mixture, TFA/TIPS/1,8-dithioloctane (94:3:3, v/v/v). The precipitated crude peptide was washed twice with $Et_2O$ and dried under vacuum. The solid crude peptide obtained was then subjected to preparative HPLC purification.

**C-terminal thioesterification**

Peptide C-terminal thioesterification was performed similarly to a previous report [1]. Briefly, the peptide was cleaved from the resin using a mixture of 1,1,1,3,3,3-hexafluoro-2-propanol (HFIP)–dichloromethane (20:80, v/v). The solvent mixture was then evaporated under reduced pressure and residues of HFIP were removed by multiple co-evaporations with dichloroethane. The solid residue was then left under high vacuum overnight. The protected peptide was dissolved in dimethylsulfoxide (DMSO) to reach an expected final concentration of 20 mM. 3-Thiopropanoate methyl ester (20 equiv) was added to the peptide solution, followed by DiPEA (5 equiv) and PyBOP (5 equiv). The reaction mixture was stirred at room temp for 3h to then be added dropwise to the TFA cleaving mixture described above. After 60 min, the crude peptide was precipitated by addition of $Et_2O$. The isolated peptide was again dissolved in neat TFA cleavage mixture and incubated for another 60 min. The crude peptide was precipitated in cold $Et_2O$, washed twice with $Et_2O$ and dried under vacuum. The solid crude peptide obtained was then subjected to preparative HPLC purification.

**Native Chemical Ligation_Method A**

One-pot ligation of three peptide fragments was performed similarly to a previous report [1]. Briefly, $Int^N$-B peptide was dissolved in NCL buffer (guanidinium hydrochloride 6 M, phosphate buffer 100 mM, tris(2-carboxyethyl)phosphine hydrochloride (TCEP) 20 mM, 4-mercaptophenylacetic acid (MPAA) 100 mM, pH 7) to reach 2 mM concentration, followed by addition of the ASIC1a peptide (1.0 equiv). The pH was adjusted to ~7 and the reaction mixture was incubated at room temperature. Conversion was monitored over time by LC-MS. Upon full conversion, TCEP and methoxylamine hydrochloride ($MeONH_2{\cdot}HCl$) were added to reach a final concentration of 40 and 200 mM, respectively, and the reaction mixture was incubated at 37 °C. Upon full unmasking of the N-terminal Cys residue, the pH was adjusted to ~7, MPAA was added to reach 200 mM and $Int^C$-A thioester was added (1.0 equiv). The pH was adjusted again to ~7 and the ligation was incubated at room temperature overnight. The reaction mixture was then directly subjected to semi-preparative HPLC purification.

**Native Chemical Ligation_Method B**

In-situ thioesterification of C-terminal hydrazide and following ligation of peptides were performed similarly to a previous report [2]. Briefly, the C-terminal hydrazide ASIC1a peptide was dissolved in guanidinium hydrochloride buffer (guanidinium hydrochloride 6 M, phosphate buffer 100 mM, pH ~ 7) to reach 3 mM concentration. Solid MPAA was added to reach 200 mM (the mixture is a suspension at this point; the mixture was sonicated for a few seconds in case big lumps of MPAA were present), followed by acetylacetone (3 equiv). The thioesterification was promoted by adjusting the pH to ~2.5. Approximately 5h were necessary to obtain quantitative conversion of starting material. $Int^N$-B peptide (1.0 equiv) was then dissolved in guanidinium hydrochloride buffer (guanidinium hydrochloride 6 M, phosphate buffer 100 mM, 50 mM TCEP, 200 mM MPAA, pH 7) to reach 3 mM concentration and the obtained solution was added to the thioesterification reaction mixture. The pH was adjusted to ~7 and the solution was incubated at room temperature. Upon ligation of the two peptide fragments, TCEP and $MeONH_2{\cdot}HCl$ were added to reach a final concentration of 40 and 200 mM, respectively, and the reaction mixture was incubated at 37 °C.

Upon full unmasking of N-terminal cysteine the pH was adjusted to ~7 and IntC-A thioester (1.0 equiv) was added to the reaction mixture. The pH was adjusted again to ~7 to initiate the second ligation. The reaction mixture was incubated at room temperature overnight and then subjected to semi-preparative HPLC purification.

**LC-MS analysis**
LC-MS analysis of crude reaction mixtures and pure products were performed on a Waters Acquity H-class UPLC equipped with Waters Acquity eLambda PhotoDiodeArray detector ($\lambda$ = 210-800 nm), Waters UPLC BEH C4 column (300 Å, 1.7 µm, 2.1 x 50 mm) and LCT$^{TM}$ ESI Mass Spectrometer (Column Temp = 40 °C). Samples were run using 3 mobile phases: A = $H_2O$, B = $CH_3CN$ and C = 48.75% $H_2O$ + 48.75% $CH_3CN$ + 2.5% FA at a flow rate of 0.5 mL/min. Gradient: 94% A + 2% B + 4% C to 36% A + 60% B + 4% C in 6.5 min.

MS analysis of purified products was performed on a Waters XEVO-G2 XS Q-TOF mass spectrometer equipped with a Waters UPLC BEH C4 column (300 Å, 1.7 µm, 2.1 x 50 mm) (Column Temp = 60 °C). Samples were run with a 1.6 minute gradient (run time 3 min) using two mobile phases: D = 99.9% $H_2O$ + 0.1% FA, E = 99.9% $CH_3CN$ + 0.1% FA (flow rate = 0.6 mL/min). Gradient: 94% A + 2% B + 4% C to 96% B + 4% C in 1.6 min. Data processing was performed using Waters MassLynx Mass Spectrometry Software 4.2 (deconvolution with Maxent1 function).

**Preparative HPLC purification**
Preparative RP-HPLC purifications were performed on a Waters HPLC equipped with a Waters 2489 UV/Vis detector, Waters fraction collector III and Waters XBridge BEH C18 OBD Prep Column (130 Å, 5 µm, 30 × 150 mm). Samples were run at a flowrate = 37.5 mL/min. Mobile phase: A = $H_2O$, B = $CH_3CN$ and C = 99% $H_2O$ + 1% TFA in $H_2O$. Fraction collection was triggered by UV intensity ($\lambda$ = 210 nm).

**Semi-preparative HPLC purification**
Semi-preparative RP-HPLC purifications were performed on a Shimadzu LC-20AT HPLC system equipped with a Shimadzu SPD-20A UV/Vis detector, a Shimadzu FRC-10A fraction collector and a Vydac 208TP C8 Column (300 Å, 10 µm, 10 × 250 mm) was used (Column Temp = 40 °C). Samples were run at a flowrate = 6.50 mL/min. Mobile phase: A = 99.95% $H_2O$ + 0.05% TFA and B = 99.95% $CH_3CN$ + 0.05% TFA. Gradient: 80% A + 20% B to 60% A + 40% solvent B in 25 min.

#### 3B. Peptide Synthesis – details

##### Int$^N$-B

SPPS and peptide cleavage were performed as outlined above in the general section on a 50 µmol scale. Purification was performed according to the general preparative HPLC purification method, with the following gradient: 80% A + 15% B + 5% D to 70% A + 25% B + 5% D in 15 min. A white fluffy material was obtained after lyophilization of pooled fractions (26.6 mg, 24% yield calculated as TFA salt).

##### Int$^C$-A_thioester

SPPS, thioesterification and peptide cleavage were performed as outlined above in the general section on a 50 µmol scale. Purification was performed according to the general preparative HPLC purification method, with the following gradient: 80% A + 15% B + 5% D to 60% A + 35% B + 5% D in 20 min. A white fluffy material was obtained after lyophilization of pooled fractions (46.0 mg, 21% yield calculated as TFA salt).

##### ASIC1a_Lys

SPPS and peptide cleavage were performed as outlined above in the general section on a 20 µmol scale; During the synthesis the Fmoc-L-Asp(O*t*Bu)-(Dmb)Gly-OH dipeptide building block was used. Purification was performed according to the general preparative HPLC purification method, with the following gradient: 93% A + 2% B + 5% D to 75% A + 20% B + 5% D in 25 min. A white fluffy material was obtained after lyophilization of pooled fractions (6.2 mg, 11% yield calculated as TFA salt).

##### ASIC1a_Orn

SPPS, thioesterification and peptide cleavage were performed as outlined above in the general section on a 20 µmol scale; During the synthesis the Fmoc-L-Asp(O*t*Bu)-(Dmb)Gly-OH dipeptide and Fmoc-Orn(Boc)-OH building blocks were used. Purification was performed according to the general preparative HPLC purification method, with the following gradient: 85% A + 10% B + 5% D to 65% A + 30% B + 5% D in 25 min. A white fluffy material was obtained after lyophilization of pooled fractions (3.7 mg, 6% yield calculated as TFA salt).

##### ASIC1a_hLys

SPPS, thioesterification and peptide cleavage were performed as outlined above in the general section on a 20 µmol scale; During the synthesis the Fmoc-L-Asp(O*t*Bu)-(Dmb)Gly-OH dipeptide and Fmoc-HLys(Boc)-OH building blocks were used. Purification was performed according general method A, with the following gradient: 90% A + 5% B + 5% D to 70% A + 25% B + 5% D in 25 min. A white fluffy material was obtained after lyophilization of pooled fractions (12.0 mg, 21% yield calculated as TFA salt).

##### ASIC1a_NleuOH

SPPS, thioesterification and peptide cleavage were performed as outlined above in the general section on a 20 µmol scale; During the synthesis the Fmoc-L-Asp(O*t*Bu)-(Dmb)Gly-OH dipeptide and Fmoc-L-Nleu(6-O*t*Bu)-OH building blocks were used. Purification was performed according to the general preparative HPLC purification method, with the following gradient: 90% A + 5% B + 5% D to 60% A + 35% B + 5% D in 25 min. A white fluffy material was obtained after lyophilization of pooled fractions (11.4 mg, 20% yield calculated as TFA salt).

**ASIC1a_thiaLys**

SPPS and peptide cleavage were performed as outlined above in the general section on a 20 µmol scale; During the synthesis the Fmoc-L-Asp(O*t*Bu)-(Dmb)Gly-OH dipeptide building block was used. Alkylation of the cysteine residue was performed on the crude peptide as previously described [3]. Briefly, ethylene imine was prepared by incubating a 2 M solution of 2-bromoethylamine hydrobromide in water with NaOH (2.2 equiv) at 55 °C for 5 min. Cysteine alkylation was performed in 300 mM Tris-Cl buffer pH 8.9 and 25 mM freshly prepared ethylene imine. After 5 h at room temperature, extra ethylene imine was added to reach 37.5 mM and the reaction mixture was further incubated for 1 h. Purification was performed according to the general preparative HPLC purification method, with the following gradient: 90% A + 5% B + 5% D to 60% A + 35% B + 5% D in 25 min. A white fluffy material was obtained after lyophilization of pooled fractions (2.5 mg, 4% yield calculated as TFA salt).

**PeptideX_Lys**

In situ thioesterification and ligations were performed according to general Native Chemical Ligation_Method B described above on a $1.7*10^{-7}$ mol scale. Purification was performed according to the general semi-preparative HPLC purification method. A white fluffy material was obtained after lyophilization of pooled fractions (0.6 mg, 37% yield calculated as TFA salt).

**PeptideX_Orn**

Ligations were performed according to general Native Chemical Ligation_Method A described above on a $6.9*10^{-7}$ mol scale. Purification was performed according to the general semi-preparative HPLC purification method. A white fluffy material was obtained after lyophilization of pooled fractions (2.2 mg, 34% yield calculated as TFA salt).

**PeptideX_hLys**

Ligations were performed according to general Native Chemical Ligation_Method A described above on a $6.9*10^{-7}$ mol scale. Purification was performed according to the general semi-preparative HPLC purification method. A white fluffy material was obtained after lyophilization of pooled fractions (2.7 mg, 42% yield calculated as TFA salt).

**PeptideX_NleuOH**

Ligations were performed according to general Native Chemical Ligation_Method A described above on a $5.4*10^{-7}$ mol scale. Purification was performed according to the general semi-preparative HPLC purification method. A white fluffy material was obtained after lyophilization of pooled fractions (1.5 mg, 30% yield calculated as TFA salt).

**PeptideX_thiaLys**

In situ thioesterification and ligations were performed according to general Native Chemical Ligation_Method B described above on a $3.4*10^{-7}$ mol scale. Purification was performed according to the general semi-preparative HPLC purification method. A white fluffy material was obtained after lyophilization of pooled fractions (0.9 mg, 28% yield calculated as TFA salt).

##### 3C. Molecular Biology

The wild-type mASIC1a gene used in this study was cloned in a pSP64 vector between the BamHI and SacI sites.[4] This construct was also used to generate the Thr214Ser mutant. The remaining split-intein sequence-containing constructs were ordered from Twist Bioscience as gene fragments cloned into the pUNIV vector between the NheI and XhoI sites.

Site-directed mutagenesis was performed by PCR using *Pfu*UltraII fusion polymerase (Agilent Technologies) and mutagenesis primers ordered from either Eurofins Genomics or Sigma Aldrich to introduce mutations (substitutions and deletions) in the mASIC1a channel sequence or optimize the native sequence of either the split inteins or the extein sequence of peptide X. Sanger sequencing of the full coding frame confirmed all mutations (Eurofins Genomics and Macrogen Europe).

cDNAs in pSP64 vector were linearized with EcoRI and those in pUNIV vector were linearized with NotI-HF (both from New England Biolabs) and capped mRNA was transcribed with the Ambion mMESSAGE mMACHINE SP6 and T7 kits (Thermo Fisher Scientific) for constructs in pSP64 and pUNIV vectors, respectively, as per recommendations of the manufacturer. mRNAs concentration was quantified using a NanoDrop One$^{C}$ instrument (Thermo Fisher Scientific) and mRNA was subsequently stored at -20 ºC.

##### 3D. Preparation of *Xenopus laevis* oocytes and mRNA injections

Oocytes were surgically extracted from female *Xenopus laevis* frogs under anesthesia with 0.3% tricaine. Following dissection of oocyte lobes and washing in $Ca^{2+}$-free OR2 (in mM; 82.5 NaCl, 2.5 KCl, 1 $MgCl_2$ and 5 HEPES; pH 7.4), oocyte digestion was performed with 1.5 mg/mL type 1 collagenase (Nordmark Biochemicals) dissolved in OR2 buffer and placed on a bio dancer shaker (New Brunswick Scientific). Following digestion and multiple washes in $Ca^{2+}$-free OR2 the oocytes were stored in OR2 at 18 ºC.

Injection needles made from glass capillaries (World Precision Instruments, item-nr. 504949) pulled by a Flaming/Brown Micropipette Puller Model P-1000 (Sutter Instrument), filled with either mRNA or peptide-containing solutions, were used for injecting at the equatorial region of defolliculated *Xenopus laevis* oocytes. Microinjections were executed using a Nanoliter 2010 microinjector (World Precision Instruments).

mASIC1a$^{WT}$ and mASIC1a$^{T214S}$ mRNA were injected into the defolliculated *Xenopus laevis* oocytes at 250 ng/µL (18.4 nL) and 500 ng/µL (23.8 nL), respectively. In order to check for the expression of spliced (reconstituted) mASIC1a from recombinant TMD, X$^{REC}$ and C-TMD fragments of the 11 split intein combinations examined in this study, a mRNA mix of 1000 ng/µL solutions of the recombinant N-TMD, X$^{REC}$ and C-TMD fragments (27.6 nL) were injected in oocytes. For control cells, volume of X$^{REC}$ mRNA was substituted with equal volume of nuclease-free water. While recording for mASIC1a channels reconstituted with synthetic peptides (X$^{SYN}$), 27.6 nL mRNA solution containing mRNAs for N-TMD and C-TMD fragments were injected on the day of oocyte defolliculation (day 1) and the peptide X$^{SYN}$ (750 µM solutions in nuclease-free water) was injected in the same oocytes the subsequent day (day 2) at 18.4 or 23.0 nL volume. Injection volumes $\geq$ 27.6 nL resulted in higher mortality of the oocytes. Injected oocytes were then transferred to a new petri dish containing ND96 or OR2 buffer with antibiotics (in mM: 96 NaCl, 2 KCl, 1 $MgCl_2$, 5 HEPES, 1.8 $CaCl_2$, 2.5 sodium pyruvate, 0.5 theophylline, 50 µg/mL gentamycin and tetracyclin; pH 7.4) and were stored at 18 ºC for up to six days post-injection.

##### 3E. Electrophysiological recordings

On the recording day, oocytes were subjected to two-electrode voltage clamp (TEVC) electrophysiology. Current measurements were performed from oocytes placed in a recording chamber [5] perfused continuously with $Ca^{2+}$-free ND96 solution (in mM: 96 NaCl, 2 KCl, 1.8 $BaCl_2$, 1 $MgCl_2$ and 5 HEPES, pH 7.4) through an automated 8-line, gravity-driven perfusion system operated by a ValveBank™ module (AutoMate Scientific) at a flow rate of ~ 1.5 mL/min. TEVC recordings were executed with an Oocyte Clamp OC-725C amplifier (Warner Instrument Corp.) at a holding potential of -60 mV. The measured currents were acquired at 2000 Hz via Axon™ Digidata® 1550 low-noise data acquisition system (Axon Instruments) using glass capillaries prepared from (Harvard Apparatus, item no. 30-0047, 1.2 OD x 0.94 x 100 L mm) backfilled with 3 M sterile filtered KCl (with microelectrode resistance between 0.1-1.0 MΩ). A 50-60 Hz Hum Bug noise eliminator (Quest Scientific) was used to minimize noise from the recordings. Current traces were visualized using the Clampex 10.7 software (Molecular Devices) and were digitally filtered at 10 Hz with an eight-pole Bessel filter for viewing and analysis. Solutions used for channel activation or SSD protocols were prepared from the $Ca^{2+}$-free ND96 solution or a MES-ND96 w/o $Ca^{2+}$ (for pH < 5.5) (in mM: 96 NaCl, 2 KCl, 1.8 $BaCl_2$, 2 MgCl and 5 MES), while low-chloride buffer consisted of (in mM): 10 NaCl, 85 sodium methanesulfonate, 2 KCl, 1.8 $BaCl_2$, 1 $MgCl_2$ and 5 HEPES. To adjust the pH of the solutions, 0.1-2 M solutions of either HCl or NaOH. Activation as well as SSD recordings from oocytes injected with either mASIC1a$^{WT}$ or mASIC1a$^{T214S}$ could be performed one-day post injection, whereas for the oocytes that were injected with N+C+X$^{REC}$ were recorded from 3-4 days post RNA injection. For the oocytes that were injected with N+C+X$^{SYN}$, were recorded 1-2 days post-injection with peptide X$^{SYN}$ solutions. The oocytes were exposed to different activation buffers with decreasing pH-values for 20 sec and were allowed to recover in the pH 7.4 buffer for 60 sec between each subsequent activation buffer exposure. While performing the SSD protocols, oocytes were perfused for 80 sec in the desensitization buffer before being switched to the activation buffer of either pH 5.5 [mASIC1a$^{WT}$, mASIC1a$^{T214S}$, N+C+X$^{REC}$, N+C+X(Lys211Lys$^{SYN}$), N+C+X(Lys211Orn$^{SYN}$)] or 4.5 [N+C+X(Lys211hLys$^{SYN}$), N+C+X(Lys211NleuOH$^{SYN}$), N+C+X(Lys211thioLys$^{SYN}$)], while allowing for recovery in pH 7.4 buffer for 60 sec after each activation step.

##### 3F. Molecular modeling

Structure PDB ID 5WKU was used as starting template, in which chain A and residues 42-73 (transmembrane region) and any other unnecessary ligands were removed. hLys was constructed as described previously[6] and the Rosetta Scripts Relax mover was used with the following options: repeats="3" min_type="lbfgs_armijo_nonmonotone" movemap_disables_packing_of_fixed_chi_positions="true" on a movemap that allowed backbone and sidechain minimization for residues 209-214. Jump minimization was disabled. Modeling was performed with Rosetta release 2021.50+master.4ff291ed825. All plotting was performed in Prism, whereas structural models were made using PyMol 2.5.2.

#### 4. QUANTIFICATION AND STATISTICAL ANALYSIS

Current traces were obtained using Clampfit 11.2 software (Molecular Devices). The recorded ASIC1a currents of the activation and SSD from all constructs were normalized and fitted with a Hill equation to obtain the values of the pH of half-maximal activation or SSD ($pH_{50}$). Currents amplitudes (µA) measured from mASIC1a spliced with different split intein combinations were plotted as a function of days post-injection of RNA and presented as mean ± SD. Desensitization kinetics were assessed using a single exponential function. Number of replicates (n) represents individual experimental oocytes. For experiments in Figure 5G, the same oocyte was activated 2-3 times and $\tau_{desensitization}$ was averaged to one replicate. Experimental data were obtained from at least two batches of oocytes. The activation, SSD and desensitization kinetics data from all the constructs were compared for statistical analyses using unpaired t-tests to their reference controls. Statistical details of experiments can be found in the figure legends. All data analyses were performed using Prism 9.2.0 (GraphPad Software). Figures were generated with Adobe Illustrator 2022 (Adobe Corp).

#### 5. KEY RESOURCES TABLE

| REAGENT or RESOURCE | SOURCE | IDENTIFIER |
|---|---|---|
| Chemicals, peptides, and recombinant proteins | | |
| PeptideX_Lys | This manuscript | Not available |
| PeptideX_Orn | This manuscript | Not available |
| PeptideX_hLys | This manuscript | Not available |
| PeptideX_NleuOH | This manuscript | Not available |
| PeptideX_thiaLys | This manuscript | Not available |
| Experimental models: Cell lines | | |
| Experimental models: Organisms/strains | | |
| Stage 5 or 6 oocytes from *Xenopus laevis* | Xenopus 1 | Not available |
| Oligonucleotides | | |
| As described in Resource tables T1 and T3 | This manuscript | Not available |
| Software and algorithms | | |
| Clampfit 11.2 | Molecular Devices | RRID:SCR_011323 |
| Prism 9.2.0 | GraphPad Software | RRID:SCR_002798 |
| Adobe Illustrator 2022 | Adobe Corp | RRID:SCR_010279 |
| MassLynx Mass Spectrometry Software 4.2 | Waters | RRID:SCR_014271 |
| ChemOffice 20.0 | Perkin Elmer | RRID:SCR_016768 |
| Rosetta release 2021.50+master.4ff291ed825 | Rosetta | RRID:SCR_015701 |
| PyMol (2.5.2) | PyMol | RRID:SCR_000305 |

**5A. Resource Table T1:** Split intein linked mASIC1a constructs (Native sequence)

| Split intein combination (Intein A - Intein B) | Name of construct | Sequence |
|---|---|---|
| *Cfa*DnaE-<br>*Ssp*DnaB$^{M86}$ | N-Fragment:<br>mASIC1a_(1-193)-<br>*Cfa*DnaE$^{N}$ | <u>Amino acid sequence:</u><br>MELKTEEEEVGGVQPVSIQAFASSSTLHGLAHIFSYERLSLKRALWALC<br>FLGSLAVLLCVCTERVQYYFCYHHVTKLDEVAASQLTFPAVTLCNLNE<br>FRFSQVSKNDLYHAGELLALLNNRYEIPDTQMADEKQLEILQDKANFR<br>SFKPKPFNMREFYDRAGHDIRDMLLSCHFRGEACSAEDFKVVFTRYGK<br>CLSYDTEILTVEYGFLPIGKIVEERIECTVYTVDKNGFVYTQPIAQWHNR<br>GEQEVFEYCLEDGSIIRATKDHKFMTTDGQMLPIDEIFERGLDLKQVDG<br>LP*<br><br><u>Nucleotide sequence:</u><br>ATGGAACTCAAGACCGAGGAAGAGGAAGTCGGCGGCGTTCAGCCA<br>GTGTCCATTCAGGCCTTTGCCAGCAGCAGCACACTGCACGGACTGG<br>CCCACATCTTCAGCTACGAGAGACTGAGCCTGAAGCGGGCCCTGTG<br>GGCTCTGTGTTTTCTGGGATCTCTGGCCGTGCTGCTGTGCGTGTGTA<br>CAGAGAGAGTGCAGTACTACTTCTGCTACCACCACGTGACCAAGCT<br>GGATGAGGTGGCCGCTTCTCAGCTGACATTCCCTGCCGTGACACTGT<br>GCAACCTGAACGAGTTCCGGTTCTCCCAGGTGTCCAAGAACGACCT<br>GTATCACGCCGGCGAACTGCTGGCCCTGCTGAACAACAGATACGAG<br>ATCCCCGACACACAGATGGCCGACGAGAAGCAGCTGGAAATTCTGC<br>AGGACAAGGCCAACTTCAGAAGCTTCAAGCCCAAGCCGTTCAACAT<br>GCGCGAGTTCTACGATAGAGCCGGCCACGACATCCGGGACATGCTG<br>CTGAGCTGTCACTTTAGAGGCGAGGCCTGTAGCGCCGAGGACTTCA<br>AGGTGGTGTTCACCAGATACGGCAAGTGCCTGAGCTACGACACCGA<br>GATCCTGACCGTGGAATACGGCTTCCTGCCTATCGGCAAGATCGTGG<br>AAGAACGGATCGAGTGCACCGTGTACACCGTGGACAAGAACGGCTT<br>CGTGTACACACAGCCTATCGCTCAGTGGCACAACCGGGGAGAGCAA<br>GAGGTGTTCGAGTACTGCCTGGAAGATGGCAGCATCATCCGGGCCA<br>CCAAGGACCACAAGTTCATGACCACCGACGGCCAGATGCTGCCCAT<br>CGACGAGATCTTTGAGAGAGGCCTGGACCTGAAACAGGTGGACGGA<br>CTGCCTTGA |
| | Peptide X:<br>*Cfa*DnaE$^{C}$-<br>mASIC1a_(194-213)-<br>*Ssp*DnaB$^{M86-N}$-linker-<br>ER retention sequence | <u>Amino acid sequence:</u><br>MVKIISRKSLGTQNVYDIGVEKDHNFLLKNGLVASNCYTFNSGQDGRP<br>RLKTMKGGCISGDSLISLASSGESKDEL*<br><br><u>Nucleotide sequence:</u><br>ATGGTCAAGATCATCAGCCGGAAGTCCCTGGGCACCCAGAACGTGT<br>ACGATATCGGCGTGGAAAAGGACCACAACTTTCTGCTGAAGAACGG<br>CCTGGTGGCCAGCAACTGCTACACCTTCAATAGCGGCCAGGACGGC<br>AGACCCCGGCTGAAAACAATGAAGGGCGGCTGCATCAGCGGCGAC<br>AGCCTGATTTCTCTGGCCAGCTCTGGCGAGAGCAAGGACGAACTGT<br>AA |
| | C-fragment:<br>IgK *faux* TMD-HA tag-linker-<br>*Ssp*DnaB$^{M86_C}$-<br>mASIC1a_(214-526) | <u>Amino acid sequence:</u><br>METDTLLLWVLLLWVPGSTGDYPYDVPDYAGSAGSAAGSGEFSTGKR<br>VPIKDLLGEKDFEIWAINEQTMKLESAKVSRVFCTGKKLVYTLKTRLGR<br>TIKATANHRFLTIDGWKRLDELSLKEHIALPRKLESSSLQLAPEIEKLPQS<br>DIYWDPIVSITETGVEEVFDLTVPGLRNFVANDIIVHNSGNGLEIMLDIQ<br>QDEYLPVWGETDETSFEAGIKVQIHSQDEPPFIDQLGFGVAPGFQTFVSC<br>QEQRLIYLPSPWGTCNAVTMDSDFFDSYSITACRIDCETRYLVENCNCR<br>MVHMPGDAPYCTPEQYKECADPALDFLVEKDQEYCVCEMPCNLTRYG<br>KELSMVKIPSKASAKYLAKKFNKSEQYIGENILVLDIFFEVLNYETIEQK<br>KAYEIAGLLGDIGGQMGLFIGASILTVLELFDYAYEVIKHRLCRRGKCQ<br>KEAKRNSADKGVALSLDDVKRHNPCESLRGHPAGMTYAANILPHHPA<br>RGTFEDFTC*<br><br><u>Nucleotide sequence:</u><br>ATGGAAACGGACACCCTGCTGCTGTGGGTGCTGTTGTTGTGGGTGCC<br>AGGCAGCACAGGCGACTACCCTTACGATGTGCCTGATTACGCCGGC<br>AGCGCTGGATCTGCTGCTGGAAGCGGAGAGTTTAGCACCGGCAAGA<br>GAGTGCCCATCAAGGACCTGCTGGGCGAGAAGGACTTTGAGATCTG<br>GGCCATCAACGAGCAGACCATGAAGCTGGAAAGCGCCAAGGTGTCC<br>CGGGTGTTCTGTACCGGCAAAAAGCTGGTGTACACACTGAAAACCC |

| | | |
|---|---|---|
| | | GGCTGGGCAGAACCATCAAGGCCACCGCCAACCACCGGTTTCTGAC<br>AATCGACGGCTGGAAGAGACTGGACGAGCTGAGCCTGAAAGAGCA<br>CATTGCCCTGCCTAGAAAGCTGGAATCCAGCAGCCTGCAGCTGGCC<br>CCTGAGATTGAGAAGCTGCCCCAGAGCGACATCTACTGGGACCCCA<br>TCGTGTCCATCACCGAGACAGGCGTGGAAGAGGTGTTCGACCTGAC<br>AGTGCCCGGCCTGAGAAACTTCGTGGCCAACGACATCATCGTGCAC<br>AACAGCGGCAACGGCCTGGAAATCATGCTGGACATTCAGCAGGACG<br>AGTACCTGCCTGTGTGGGGCGAGACAGACGAGACATCTTTTGAGGC<br>CGGCATCAAGGTGCAGATCCACAGCCAGGATGAGCCTCCATTCATC<br>GACCAGCTCGGCTTTGGAGTGGCCCCTGGCTTTCAGACCTTCGTGTC<br>CTGCCAAGAGCAGCGGCTGATCTACCTGCCTTCTCCTTGGGGCACCT<br>GTAACGCCGTGACCATGGACAGCGATTTCTTCGACAGCTACAGCAT<br>CACCGCCTGCCGGATCGACTGCGAGACAAGATACCTGGTGGAAAAC<br>TGCAACTGCCGGATGGTGCACATGCCTGGCGACGCCCCTTACTGTAC<br>ACCCGAGCAGTACAAAGAGTGCGCCGATCCTGCTCTGGACTTCCTG<br>GTTGAGAAGGACCAAGAGTACTGCGTGTGCGAGATGCCCTGCAACC<br>TGACCAGATACGGCAAAGAACTGAGCATGGTCAAGATCCCCAGCAA<br>GGCCTCTGCCAAGTACCTGGCCAAGAAGTTCAACAAGAGCGAGCAG<br>TATATCGGCGAGAACATCCTGGTGCTGGATATCTTCTTCGAGGTGCT<br>GAACTACGAGACAATCGAGCAGAAGAAGGCCTACGAGATCGCCGG<br>CCTGCTGGGAGATATTGGCGGACAGATGGGCCTGTTTATCGGCGCC<br>AGCATCCTGACCGTGCTGGAACTGTTCGACTACGCCTACGAAGTGAT<br>CAAGCACCGGCTGTGCAGACGGGGCAAGTGTCAGAAAGAGGCCAA<br>GAGAAACAGCGCCGACAAAGGCGTGGCCCTGAGCCTGGATGATGTG<br>AAGAGACACAACCCCTGCGAGAGCCTGAGAGGACATCCTGCCGGAA<br>TGACCTACGCCGCCAACATTCTGCCTCATCACCCTGCCAGAGGCACC<br>TTCGAGGACTTCACCTGTTGA |
| *Cfa*DnaE-*Ssp*DnaB$^{M86}$ | N-C linked construct:<br><br>mASIC1a_(1-193)-*Cfa*DnaE$^{N}$-(GGGGS)$_6$ linker-*Ssp*DnaB$^{M86_C}$-mASIC1a_(214-526) | Amino acid sequence:<br>MELKTEEEEVGGVQPVSIQAFASSSTLHGLAHIFSYERLSLKRALWALC<br>FLGSLAVLLCVCTERVQYYFCYHHVTKLDEVAASQLTFPAVTLCNLNE<br>FRFSQVSKNDLYHAGELLALLNNRYEIPDTQMADEKQLEILQDKANFR<br>SFKPKPFNMREFYDRAGHDIRDMLLSCHFRGEACSAEDFKVVFTRYGK<br>CLSYDTEILTVEYGFLPIGKIVEERIECTVYTVDKNGFVYTQPIAQWHNR<br>GEQEVFEYCLEDGSIIRATKDHKFMTTDGQMLPIDEIFERGLDLKQVDG<br>LPGGGGSGGGGSGGGGSGGGGSGGGGSGGGGSSTGKRVPIKDLLGEK<br>DFEIWAINEQTMKLESAKVSRVFCTGKKLVYTLKTRLGRTIKATANHR<br>FLTIDGWKRLDELSLKEHIALPRKLESSSLQLAPEIEKLPQSDIYWDPIVS<br>ITETGVEEVFDLTVPGLRNFVANDIIVHNSGNGLEIMLDIQQDEYLPVW<br>GETDETSFEAGIKVQIHSQDEPPFIDQLGFGVAPGFQTFVSCQEQRLIYLP<br>SPWGTCNAVTMDSDFFDSYSITACRIDCETRYLVENCNCRMVHMPGD<br>APYCTPEQYKECADPALDFLVEKDQEYCVCEMPCNLTRYGKELSMVKI<br>PSKASAKYLAKKFNKSEQYIGENILVLDIFFEVLNYETIEQKKAYEIAGL<br>LGDIGGQMGLFIGASILTVLELFDYAYEVIKHRLCRRGKCQKEAKRNSA<br>DKGVALSLDDVKRHNPCESLRGHPAGMTYAANILPHHPARGTFEDFTC<br>*<br><br>Nucleotide sequence:<br>ATGGAACTCAAGACCGAGGAAGAGGAAGTCGGCGGCGTTCAGCCA<br>GTGTCCATTCAGGCCTTTGCCAGCAGCAGCACACTGCACGGACTGG<br>CCCACATCTTCAGCTACGAGAGACTGAGCCTGAAGCGGGCCCTGTG<br>GGCTCTGTGTTTTCTGGGATCTCTGGCCGTGCTGCTGTGCGTGTGTA<br>CAGAGAGAGTGCAGTACTACTTCTGCTACCACCACGTGACCAAGCT<br>GGATGAGGTGGCCGCTTCTCAGCTGACATTCCCTGCCGTGACACTGT<br>GCAACCTGAACGAGTTCCGGTTCTCCCAGGTGTCCAAGAACGACCT<br>GTATCACGCCGGCGAACTGCTGGCCCTGCTGAACAACAGATACGAG<br>ATCCCCGACACACAGATGGCCGACGAGAAGCAGCTGGAAATTCTGC<br>AGGACAAGGCCAACTTCAGAAGCTTCAAGCCCAAGCCGTTCAACAT<br>GCGCGAGTTCTACGATAGAGCCGGCCACGACATCCGGGACATGCTG<br>CTGAGCTGTCACTTTAGAGGCGAGGCCTGTAGCGCCGAGGACTTCA<br>AGGTGGTGTTCACCAGATACGGCAAGTGCCTGAGCTACGACACCGA<br>GATCCTGACCGTGGAATACGGCTTCCTGCCTATCGGCAAGATCGTGG<br>AAGAACGGATCGAGTGCACCGTGTACACCGTGGACAAGAACGGCTT<br>CGTGTACACACAGCCTATCGCTCAGTGGCACAACCGGGGAGAGCAA<br>GAGGTGTTCGAGTACTGCCTGGAAGATGGCAGCATCATCCGGGCCA<br>CCAAGGACCACAAGTTCATGACCACCGACGGCCAGATGCTGCCCAT<br>CGACGAGATCTTTGAGAGAGGCCTGGACCTGAAACAGGTGGACGGA<br>CTGCCTGGTGGCGGTGGTTCTGGCGGTGGCGGTTCAGGCGGTGGCG |

| | | |
|---|---|---|
| | | GTTCTGGTGGCGGTGGCTCAGGCGGTGGCGGCTCAGGCGGTGGTGG<br>CTCTAGCACCGGCAAGAGAGTGCCCATCAAGGACCTGCTGGGCGAG<br>AAGGACTTTGAGATCTGGGCCATCAACGAGCAGACCATGAAGCTGG<br>AAAGCGCCAAGGTGTCCCGGGTGTTCTGTACCGGCAAAAAGCTGGT<br>GTACACACTGAAAACCCGGCTGGGCAGAACCATCAAGGCCACCGCC<br>AACCACCGGTTTCTGACAATCGACGGCTGGAAGAGACTGGACGAGC<br>TGAGCCTGAAAGAGCACATTGCCCTGCCTAGAAAGCTGGAATCCAG<br>CAGCCTGCAGCTGGCCCCTGAGATTGAGAAGCTGCCCCAGAGCGAC<br>ATCTACTGGGACCCCATCGTGTCCATCACCGAGACAGGCGTGGAAG<br>AGGTGTTCGACCTGACAGTGCCCGGCCTGAGAAACTTCGTGGCCAA<br>CGACATCATCGTGCACAACAGCGGCAACGGCCTGGAAATCATGCTG<br>GACATTCAGCAGGACGAGTACCTGCCTGTGTGGGGCGAGACAGACG<br>AGACATCTTTTGAGGCCGGCATCAAGGTGCAGATCCACAGCCAGGA<br>TGAGCCTCCATTCATCGACCAGCTCGGCTTTGGAGTGGCCCCTGGCT<br>TTCAGACCTTCGTGTCCTGCCAAGAGCAGCGGCTGATCTACCTGCCT<br>TCTCCTTGGGGCACCTGTAACGCCGTGACCATGGACAGCGATTTCTT<br>CGACAGCTACAGCATCACCGCCTGCCGGATCGACTGCGAGACAAGA<br>TACCTGGTGGAAAACTGCAACTGCCGGATGGTGCACATGCCTGGCG<br>ACGCCCCTTACTGTACACCCGAGCAGTACAAAGAGTGCGCCGATCC<br>TGCTCTGGACTTCCTGGTTGAGAAGGACCAAGAGTACTGCGTGTGC<br>GAGATGCCCTGCAACCTGACCAGATACGGCAAAGAACTGAGCATGG<br>TCAAGATCCCCAGCAAGGCCTCTGCCAAGTACCTGGCCAAGAAGTT<br>CAACAAGAGCGAGCAGTATATCGGCGAGAACATCCTGGTGCTGGAT<br>ATCTTCTTCGAGGTGCTGAACTACGAGACAATCGAGCAGAAGAAGG<br>CCTACGAGATCGCCGGCCTGCTGGGAGATATTGGCGGACAGATGGG<br>CCTGTTTATCGGCGCCAGCATCCTGACCGTGCTGGAACTGTTCGACT<br>ACGCCTACGAAGTGATCAAGCACCGGCTGTGCAGACGGGGCAAGTG<br>TCAGAAAGAGGCCAAGAGAAACAGCGCCGACAAAGGCGTGGCCCT<br>GAGCCTGGATGATGTGAAGAGACACAACCCCTGCGAGAGCCTGAGA<br>GGACATCCTGCCGGAATGACCTACGCCGCCAACATTCTGCCTCATCA<br>CCCTGCCAGAGGCACCTTCGAGGACTTCACCTGTTGA |
| NrdJ1- *Ssp*DnaB$^{M86}$ | N-Fragment:<br>mASIC1a_(1-179)-<br>Nrdj1$^{N}$ | Amino acid sequence:<br>MELKTEEEEVGGVQPVSIQAFASSSTLHGLAHIFSYERLSLKRALWALC<br>FLGSLAVLLCVCTERVQYYFCYHHVTKLDEVAASQLTFPAVTLCNLNE<br>FRFSQVSKNDLYHAGELLALLNNRYEIPDTQMADEKQLEILQDKANFR<br>SFKPKPFNMREFYDRAGHDIRDMLLSCHFRGEACCLVGSSEIITRNYGK<br>TTIKEVVEIFDNDKNIQVLAFNTHTDNIEWAPIKAAQLTRPNAELVELEI<br>DTLHGVKTIRCTPDHPVYTKNRGYVRADELTDDDELVVAI*<br><br>Nucleotide sequence:<br>ATGGAACTCAAGACCGAGGAAGAGGAAGTCGGCGGCGTTCAGCCA<br>GTGTCCATTCAGGCCTTTGCCAGCAGCAGCACACTGCACGGACTGG<br>CCCACATCTTCAGCTACGAGAGACTGAGCCTGAAGCGGGCCCTGTG<br>GGCTCTGTGTTTTCTGGGATCTCTGGCCGTGCTGCTGTGCGTGTGTA<br>CAGAGAGAGTGCAGTACTACTTCTGCTACCACCACGTGACCAAGCT<br>GGATGAGGTGGCCGCTTCTCAGCTGACATTCCCTGCCGTGACACTGT<br>GCAACCTGAACGAGTTCCGGTTCTCCCAGGTGTCCAAGAACGACCT<br>GTATCACGCCGGCGAACTGCTGGCCCTGCTGAACAACAGATACGAG<br>ATCCCCGACACACAGATGGCCGACGAGAAGCAGCTGGAAATTCTGC<br>AGGACAAGGCCAACTTCAGAAGCTTCAAGCCCAAGCCGTTCAACAT<br>GCGCGAGTTCTACGATAGAGCCGGCCACGACATCCGGGACATGCTG<br>CTGAGCTGTCACTTTAGAGGCGAGGCCTGTTGTCTGGTGGGATCTTC<br>TGAAATTATTACAAGAAATTATGGAAAAACAACAATTAAAGAAGTG<br>GTGGAAATTTTTGATAATGATAAAAATATTCAGGTGCTGGCTTTTAA<br>TACACACACAGATAATATTGAATGGGCTCCAATTAAAGCTGCTCAG<br>CTGACAAGACCAAATGCTGAACTGGTGGAACTGGAAATTGATACAC<br>TGCACGGAGTGAAAACAATTAGATGTACACCAGATCACCCAGTGTA<br>TACAAAAAATAGAGGATATGTGAGAGCTGATGAACTGACAGATGAT<br>GATGAACTGGTGGTGGCTATTTGA |
| | Peptide X:<br>NrdJ1$^{C}$-<br>mASIC1a_(180-213)-<br>*Ssp*DnaB$^{M86-N}$-linker-<br>ER retention sequence | Amino acid sequence:<br>MEAKTYIGKLKSRKIVSNEDTYDIQTSTHNFFANDILVHNSAEDFKVVF<br>TRYGKCYTFNSGQDGRPRLKTMKGGCISGDSLISLASSGESKDEL*<br><br>Nucleotide sequence:<br>ATGGAAGCTAAAACATATATTGGAAAACTGAAATCTAGAAAAATTG<br>TGTCTAATGAAGATACATATGATATTCAGACATCTACACACAATTTT<br>TTTGCTAATGATATTCTGGTGCACAATTCTGCCGAGGACTTCAAGGT |

| | | |
|---|---|---|
| | | GGTGTTCACCAGATACGGCAAGTGCTACACCTTCAATAGCGGCCAG<br>GACGGCAGACCCCGGCTGAAAACAATGAAGGGCGGCTGCATCAGC<br>GGCGACAGCCTGATTTCTCTGGCCAGCTCTGGCGAGAGCAAGGACG<br>AACTGTAA |
| | C-fragment:<br>IgK *faux* TMD-HA tag-linker *Ssp*DnaB$^{M86_C}$-mASIC1a_(214-526) | Same as shown for *Cfa*DnaE-*Ssp*DnaB$^{M86}$ split intein pair |
| *Mja*KlbA-*Ssp*DnaB$^{M86}$ | N-Fragment:<br>mASIC1a_(1-193)-*Mja*KlbA$^{N}$ | Amino acid sequence:<br>MELKTEEEEVGGVQPVSIQAFASSSTLHGLAHIFSYERLSLKRALWALC<br>FLGSLAVLLCVCTERVQYYFCYHHVTKLDEVAASQLTFPAVTLCNLNE<br>FRFSQVSKNDLYHAGELLALLNNRYEIPDTQMADEKQLEILQDKANFR<br>SFKPKPFNMREFYDRAGHDIRDMLLSCHFRGEACSAEDFKVVFTRYGK<br>ALAYDEPIYLSDGNIINIGEFVDKFFKKYKNSIKKEDNGFGWIDIGNENI<br>YIKSFNKLSLIIEDKRILRVWRKKYSGKLIKITTKNRREITLTHDHPVYIS<br>KTGEVLEINAEMVKVGDYIYIPKNNT*<br><br>Nucleotide sequence:<br>ATGGAACTCAAGACCGAGGAAGAGGAAGTCGGCGGCGTTCAGCCA<br>GTGTCCATTCAGGCCTTTGCCAGCAGCAGCACACTGCACGGACTGG<br>CCCACATCTTCAGCTACGAGAGACTGAGCCTGAAGCGGGCCCTGTG<br>GGCTCTGTGTTTTCTGGGATCTCTGGCCGTGCTGCTGTGCGTGTGTA<br>CAGAGAGAGTGCAGTACTACTTCTGCTACCACCACGTGACCAAGCT<br>GGATGAGGTGGCCGCTTCTCAGCTGACATTCCCTGCCGTGACACTGT<br>GCAACCTGAACGAGTTCCGGTTCTCCCAGGTGTCCAAGAACGACCT<br>GTATCACGCCGGCGAACTGCTGGCCCTGCTGAACAACAGATACGAG<br>ATCCCCGACACACAGATGGCCGACGAGAAGCAGCTGGAAATTCTGC<br>AGGACAAGGCCAACTTCAGAAGCTTCAAGCCCAAGCCGTTCAACAT<br>GCGCGAGTTCTACGATAGAGCCGGCCACGACATCCGGGACATGCTG<br>CTGAGCTGTCACTTTAGAGGCGAGGCCTGTAGCGCCGAGGACTTCA<br>AGGTGGTGTTCACCAGATACGGCAAGGCTCTGGCTTATGATGAACC<br>AATTTATCTGTCTGATGGAAATATTATTAATATTGGAGAGTTCGTAG<br>ACAAGTTCTTCAAGAAGTACAAGAACTCCATCAAGAAAGAGGATAA<br>TGGATTTGGATGGATTGATATTGGAAATGAAAATATTTATATTAAAT<br>CTTTTAATAAACTGTCTCTGATTATTGAAGATAAAAGAATTCTGAGA<br>GTGTGGAGAAAAAAATATTCTGGAAAACTGATTAAAATTACAACAA<br>AAAATAGAAGAGAAATTACACTGACACACGATCACCCAGTGTATAT<br>TTCTAAAACAGGAGAAGTGCTGGAAATTAATGCTGAAATGGTGAAA<br>GTGGGAGATTATATTTATATTCCAAAAAATAATACATGA |
| | Peptide X:<br>*Mja*KlbA$^{C}$-mASIC1a_(194-213)-*Ssp*DnaB$^{M86\text{-}N}$-linker-ER retention sequence | Amino acid sequence<br>MSSGSSINLDEVIKVETVDYNGHIYDLTVEDNHTYIAGKNEGFAVSNCY<br>TFNSGQDGRPRLKTMKGGCISGDSLISLASSGESKDEL*<br><br>Nucleotide sequence<br>ATGAGCTCTGGCTCTAGCATTAATCTGGATGAAGTGATTAAAGTGG<br>AAACAGTGGATTATAATGGACACATTTATGATCTGACAGTGGAAGA<br>TAATCACACATATATTGCTGGAAAAAATGAAGGATTTGCTGTGTCTA<br>ATTGCTATACATTCAATAGCGGCCAGGACGGCAGACCCCGGCTGAA<br>AACAATGAAGGGCGGCTGCATCAGCGGCGACAGCCTGATTTCTCTG<br>GCCAGCTCTGGCGAGAGCAAGGACGAACTGTAA |
| | C-fragment:<br>IgK *faux* TMD-HA tag-linker-*Ssp*DnaB$^{M86_C}$-mASIC1a_(214-526) | Same as shown for *Cfa*DnaE- *Ssp*DnaB$^{M86}$ split intein pair |
| *Npu*DnaE-*Ssp*DnaB$^{M86}$ | N-Fragment:<br>mASIC1a_(1-193)-*Npu*DnaE$^{N}$ | Amino acid sequence<br>MELKTEEEEVGGVQPVSIQAFASSSTLHGLAHIFSYERLSLKRALWALC<br>FLGSLAVLLCVCTERVQYYFCYHHVTKLDEVAASQLTFPAVTLCNLNE<br>FRFSQVSKNDLYHAGELLALLNNRYEIPDTQMADEKQLEILQDKANFR<br>SFKPKPFNMREFYDRAGHDIRDMLLSCHFRGEACSAEDFKVVFTRYGK<br>CLSYETEILTVEYGSLPIGKIVEKRIECTVYSVDNNGNIYTQPVAQWHDR<br>GEQEVFEYCLEDGSLIRATKDHKFMTVDGQMLPIDEIFERELDLMRVD<br>NLPNIKIATRKYLGKQNVYDIGVER* |

| | | |
|---|---|---|
| | | Nucleotide sequence:<br>ATGGAACTCAAGACCGAGGAAGAGGAAGTCGGCGGCGTTCAGCCA<br>GTGTCCATTCAGGCCTTTGCCAGCAGCAGCACACTGCACGGACTGG<br>CCCACATCTTCAGCTACGAGAGACTGAGCCTGAAGCGGGCCCTGTG<br>GGCTCTGTGTTTTCTGGGATCTCTGGCCGTGCTGCTGTGCGTGTGTA<br>CAGAGAGAGTGCAGTACTACTTCTGCTACCACCACGTGACCAAGCT<br>GGATGAGGTGGCCGCTTCTCAGCTGACATTCCCTGCCGTGACACTGT<br>GCAACCTGAACGAGTTCCGGTTCTCCCAGGTGTCCAAGAACGACCT<br>GTATCACGCCGGCGAACTGCTGGCCCTGCTGAACAACAGATACGAG<br>ATCCCCGACACACAGATGGCCGACGAGAAGCAGCTGGAAATTCTGC<br>AGGACAAGGCCAACTTCAGAAGCTTCAAGCCCAAGCCGTTCAACAT<br>GCGCGAGTTCTACGATAGAGCCGGCCACGACATCCGGGACATGCTG<br>CTGAGCTGTCACTTTAGAGGCGAGGCCTGTAGCGCCGAGGACTTCA<br>AGGTGGTGTTCACCAGATACGGCAAGTGCCTGAGCTATGAAACCGA<br>AATTCTGACCGTGGAATATGGCAGCCTGCCGATTGGCAAAATTGTG<br>GAAAAACGCATTGAATGCACCGTGTATAGCGTGGATAACAACGGCA<br>ACATTTATACCCAGCCGGTGGCGCAGTGGCATGATCGCGGCGAACA<br>GGAAGTGTTTGAATATTGCCTGGAAGATGGCAGCCTGATTCGCGCG<br>ACCAAAGATCATAAATTTATGACCGTGGATGGCCAGATGCTGCCGA<br>TTGATGAAATTTTTGAACGCGAACTGGATCTGATGCGCGTGGATAAC<br>CTGCCGAACATTAAAATTGCGACCCGCAAATATCTGGGCAAACAGA<br>ACGTGTATGATATTGGCGTGGAACGCTGA |
| | Peptide X:<br>*Npu*DnaE $^{C}$-<br>mASIC1a_(194-213)-<br>*Ssp*DnaB $^{M86\text{-}N}$-linker-<br>ER retention sequence | Amino acid sequence<br>MDHNFALKNGFIASNCYTFNSGQDGRPRLKTMKGGCISGDSLISLASSG<br>ESKDEL*<br><br>Nucleotide sequence<br>ATGGATCATAACTTTGCGCTGAAAAACGGCTTTATTGCGAGCAACTG<br>CTACACCTTCAATAGCGGCCAGGACGGCAGACCCCGGCTGAAAACA<br>ATGAAGGGCGGCTGCATCAGCGGCGACAGCCTGATTTCTCTGGCCA<br>GCTCTGGCGAGAGCAAGGACGAACTGTAA |
| | C-fragment:<br>IgK *faux* TMD-HA tag-linker-<br>*Ssp*DnaB $^{M86\_C}$-<br>mASIC1a_(214-526) | Same as shown for *Cfa*DnaE- *Ssp*DnaB $^{M86}$ split intein pair |
| gp41-1-*Ssp*DnaX | N-Fragment:<br>mASIC1a_(1-179)-<br>gp41-1 $^{N}$ | Amino acid sequence:<br>MELKTEEEEVGGVQPVSIQAFASSSTLHGLAHIFSYERLSLKRALWALC<br>FLGSLAVLLCVCTERVQYYFCYHHVTKLDEVAASQLTFPAVTLCNLNE<br>FRFSQVSKNDLYHAGELLALLNNRYEIPDTQMADEKQLEILQDKANFR<br>SFKPKPFNMREFYDRAGHDIRDMLLSCHFRGEACCLDLKTQVQTPQGM<br>KEISNIQVGDLVLSNTGYNEVLNVFPKSKKKSYKITLEDGKEIICSEEHL<br>FPTQTGEMNISGGLKEGMCLYVKE*<br><br>Nucleotide sequence:<br>ATGGAACTCAAGACCGAGGAAGAGGAAGTCGGCGGCGTTCAGCCA<br>GTGTCCATTCAGGCCTTTGCCAGCAGCAGCACACTGCACGGACTGG<br>CCCACATCTTCAGCTACGAGAGACTGAGCCTGAAGCGGGCCCTGTG<br>GGCTCTGTGTTTTCTGGGATCTCTGGCCGTGCTGCTGTGCGTGTGTA<br>CAGAGAGAGTGCAGTACTACTTCTGCTACCACCACGTGACCAAGCT<br>GGATGAGGTGGCCGCTTCTCAGCTGACATTCCCTGCCGTGACACTGT<br>GCAACCTGAACGAGTTCCGGTTCTCCCAGGTGTCCAAGAACGACCT<br>GTATCACGCCGGCGAACTGCTGGCCCTGCTGAACAACAGATACGAG<br>ATCCCCGACACACAGATGGCCGACGAGAAGCAGCTGGAAATTCTGC<br>AGGACAAGGCCAACTTCAGAAGCTTCAAGCCCAAGCCGTTCAACAT<br>GCGCGAGTTCTACGATAGAGCCGGCCACGACATCCGGGACATGCTG<br>CTGAGCTGTCACTTTAGAGGCGAGGCCTGTTGCCTGGATCTGAAAAC<br>CCAGGTGCAGACCCCGCAGGGCATGAAAGAAATTAGCAACATTCAG<br>GTGGGCGATCTGGTGCTGAGCAACACCGGCTATAACGAAGTGCTGA<br>ACGTGTTTCCGAAAAGCAAGAAAAAGAGCTATAAAATTACCCTGGA<br>AGATGGCAAAGAAATTATTTGCAGCGAAGAACATCTGTTTCCGACC<br>CAGACCGGCGAAATGAACATTAGCGGCGGCCTGAAAGAAGGCATGT |

| | | |
|---|---|---|
| | | GCCTGTATGTGAAAGAATGA |
| | Peptide X:<br>gp41-1$^{C}$-mASIC1a_(180-213)-*Ssp*DnaX$^{N}$-linker-ER retention sequence | Amino acid sequence:<br>MLKKILKIEELDERELIDIEVSGNHLFYANDILTHNSAEDFKVVFTRYGK<br>CYTFNSGQDGRPRLKTMKGGCLTGDSQVLTRSSGESKDEL*<br><br>Nucleotide sequence:<br>ATGCTGAAAAAAATTCTGAAAATTGAAGAACTGGATGAACGCGAAC<br>TGATTGATATTGAAGTGAGCGGCAACCATCTGTTTTATGCGAACGAT<br>ATTCTGACCCATAACAGCGCCGAGGACTTCAAGGTGGTGTTCACCA<br>GATACGGCAAGTGCTACACCTTCAATAGCGGCCAGGACGGCAGACC<br>CCGGCTGAAAACAATGAAGGGCGGCTGCCTGACCGGCGATAGCCAG<br>GTGCTGACCCGCAGCTCTGGCGAGAGCAAGGACGAACTGTAA |
| | C-fragment:<br>IgK *faux* TMD-HA tag-linker- *Ssp*DnaX$^{C}$-mASIC1a_(214-526) | Amino acid sequence:<br>METDTLLLWVLLLWVPGSTGDYPYDVPDYAGSAGSAAGSGEFNGLMS<br>IDNPQIKGREVLSYNETLQQWEYKKVLRWLDRGEKQTLSIKTKNSTVR<br>CTANHLIRTEQGWTRAENITPGMKILSPAPQWHTNFEEVESVTKGQVE<br>KVYDLEVEDNHNFVANGLLVHNCGNGLEIMLDIQQDEYLPVWGETDE<br>TSFEAGIKVQIHSQDEPPFIDQLGFGVAPGFQTFVSCQEQRLIYLPSPWG<br>TCNAVTMDSDFFDSYSITACRIDCETRYLVENCNCRMVHMPGDAPYCT<br>PEQYKECADPALDFLVEKDQEYCVCEMPCNLTRYGKELSMVKIPSKAS<br>AKYLAKKFNKSEQYIGENILVLDIFFEVLNYETIEQKKAYEIAGLLGDIG<br>GQMGLFIGASILTVLELFDYAYEVIKHRLCRRGKCQKEAKRNSADKGV<br>ALSLDDVKRHNPCESLRGHPAGMTYAANILPHHPARGTFEDFTC*<br><br>Nucleotide sequence:<br>ATGGAAACGGACACCCTGCTGCTGTGGGTGCTGTTGTTGTGGGTGCC<br>AGGCAGCACAGGCGACTACCCTTACGATGTGCCTGATTACGCCGGC<br>AGCGCTGGATCTGCTGCTGGAAGCGGAGAGTTTAACGGCCTGATGA<br>GCATTGATAACCCGCAGATTAAAGGCCGCGAAGTGCTGAGCTATAA<br>CGAAACCCTGCAGCAGTGGGAATATAAAAAAGTGCTGCGCTGGCTG<br>GATCGCGGCGAAAAACAGACCCTGAGCATTAAAACCAAAAACAGC<br>ACCGTGCGCTGCACCGCGAACCATCTGATTCGCACCGAACAGGGCT<br>GGACCCGCGCGGAAAACATTACCCCGGGCATGAAAATTCTGAGCCC<br>GGCGCCGCAGTGGCATACCAACTTTGAAGAAGTGGAAAGCGTGACC<br>AAAGGCCAGGTGGAAAAAGTGTATGATCTGGAAGTGGAAGATAAC<br>CATAACTTTGTGGCGAACGGCCTGCTGGTGCATAACTGCGGCAACG<br>GCCTGGAAATCATGCTGGACATTCAGCAGGACGAGTACCTGCCTGT<br>GTGGGGCGAGACAGACGAGACATCTTTTGAGGCCGGCATCAAGGTG<br>CAGATCCACAGCCAGGATGAGCCTCCATTCATCGACCAGCTCGGCTT<br>TGGAGTGGCCCCTGGCTTTCAGACCTTCGTGTCCTGCCAAGAGCAGC<br>GGCTGATCTACCTGCCTTCTCCTTGGGGCACCTGTAACGCCGTGACC<br>ATGGACAGCGATTTCTTCGACAGCTACAGCATCACCGCCTGCCGGAT<br>CGACTGCGAGACAAGATACCTGGTGGAAAACTGCAACTGCCGGATG<br>GTGCACATGCCTGGCGACGCCCCTTACTGTACACCCGAGCAGTACA<br>AAGAGTGCGCCGATCCTGCTCTGGACTTCCTGGTTGAGAAGGACCA<br>AGAGTACTGCGTGTGCGAGATGCCCTGCAACCTGACCAGATACGGC<br>AAAGAACTGAGCATGGTCAAGATCCCCAGCAAGGCCTCTGCCAAGT<br>ACCTGGCCAAGAAGTTCAACAAGAGCGAGCAGTATATCGGCGAGAA<br>CATCCTGGTGCTGGATATCTTCTTCGAGGTGCTGAACTACGAGACAA<br>TCGAGCAGAAGAAGGCCTACGAGATCGCCGGCCTGCTGGGAGATAT<br>TGGCGGACAGATGGGCCTGTTTATCGGCGCCAGCATCCTGACCGTG<br>CTGGAACTGTTCGACTACGCCTACGAAGTGATCAAGCACCGGCTGT<br>GCAGACGGGGCAAGTGTCAGAAAGAGGCCAAGAGAAACAGCGCCG<br>ACAAAGGCGTGGCCCTGAGCCTGGATGATGTGAAGAGACACAACCC<br>CTGCGAGAGCCTGAGAGGACATCCTGCCGGAATGACCTACGCCGCC<br>AACATTCTGCCTCATCACCCTGCCAGAGGCACCTTCGAGGACTTCAC<br>CTGTTGA |

| | | |
|---|---|---|
| gp41-1-CatInt | N-Fragment:<br>mASIC1a_(1-179)-<br>gp41-1$^{N}$ | Same as shown for gp41-1-*Ssp*DnaX split intein pair |
| | Peptide X:<br>gp41-1$^{C}$-<br>mASIC1a_(180-213)-<br>CatInt$^{N}$-linker-ER<br>retention sequence | <u>Amino acid sequence:</u><br>MLKKILKIEELDERELIDIEVSGNHLFYANDILTHNSAEDFKVVFTRYGK<br>CYTFNSGQDGRPRLKTMKGGCLSGDTMIEILDDDGIIQKISMEDLYQRL<br>ASSGESKDEL*<br><br><u>Nucleotide sequence:</u><br>ATGCTGAAAAAAATTCTGAAAATTGAAGAACTGGATGAACGCGAAC<br>TGATTGATATTGAAGTGAGCGGCAACCATCTGTTTTATGCGAACGAT<br>ATTCTGACCCATAACAGCGCCGAGGACTTCAAGGTGGTGTTCACCA<br>GATACGGCAAGTGCTACACCTTCAATAGCGGCCAGGACGGCAGACC<br>CCGGCTGAAAACAATGAAGGGCGGCTGCCTGAGCGGCGATACCATG<br>ATTGAAATTCTGGATGATGATGGCATTATTCAGAAAATTAGCATGG<br>AAGATCTGTATCAGCGCCTGGCGAGCTCTGGCGAGAGCAAGGACGA<br>ACTGTAA |
| | C-fragment:<br>IgK *faux* TMD-HA<br>tag-linker-CatInt$^{C}$-<br>mASIC1a_(214-526) | <u>Amino acid sequence:</u><br>METDTLLLWVLLLWVPGSTGDYPYDVPDYAGSAGSAAGSGEFMFKLN<br>TKNIKVLTPSGFKSFSGIQKVYKPFYHHIIFDDGSEIKCSDNHSFGKDKIK<br>ASTIKVGDYLQGKKVLYNEIVEEGIYLYDLLNVGEDNLYYTNGIVSHN<br>CGNGLEIMLDIQQDEYLPVWGETDETSFEAGIKVQIHSQDEPPFIDQLGF<br>GVAPGFQTFVSCQEQRLIYLPSPWGTCNAVTMDSDFFDSYSITACRIDC<br>ETRYLVENCNCRMVHMPGDAPYCTPEQYKECADPALDFLVEKDQEYC<br>VCEMPCNLTRYGKELSMVKIPSKASAKYLAKKFNKSEQYIGENILVLDI<br>FFEVLNYETIEQKKAYEIAGLLGDIGGQMGLFIGASILTVLELFDYAYEV<br>IKHRLCRRGKCQKEAKRNSADKGVALSLDDVKRHNPCESLRGHPAGM<br>TYAANILPHHPARGTFEDFTC*<br><br><u>Nucleotide sequence:</u><br>ATGGAAACGGACACCCTGCTGCTGTGGGTGCTGTTGTTGTGGGTGCC<br>AGGCAGCACAGGCGACTACCCTTACGATGTGCCTGATTACGCCGGC<br>AGCGCTGGATCTGCTGCTGGAAGCGGAGAGTTTATGTTTAAACTGA<br>ACACCAAAAACATTAAAGTGCTGACCCCGAGCGGCTTTAAAAGCTT<br>TAGCGGCATTCAGAAAGTGTATAAACCGTTTTATCATCATATTATTT<br>TTGATGATGGCAGCGAAATTAAATGCAGCGATAACCATAGCTTTGG<br>CAAAGATAAAATTAAAGCGAGCACCATTAAAGTGGGCGATTATCTG<br>CAGGGCAAAAAAGTGCTGTATAACGAAATTGTGGAAGAAGGCATTT<br>ATCTGTATGATCTGCTGAACGTGGGCGAAGATAACCTGTATTATACC<br>AACGGCATTGTGAGCCATAACTGCGGCAACGGCCTGGAAATCATGC<br>TGGACATTCAGCAGGACGAGTACCTGCCTGTGTGGGGCGAGACAGA<br>CGAGACATCTTTTGAGGCCGGCATCAAGGTGCAGATCCACAGCCAG<br>GATGAGCCTCCATTCATCGACCAGCTCGGCTTTGGAGTGGCCCCTGG<br>CTTTCAGACCTTCGTGTCCTGCCAAGAGCAGCGGCTGATCTACCTGC<br>CTTCTCCTTGGGGCACCTGTAACGCCGTGACCATGGACAGCGATTTC<br>TTCGACAGCTACAGCATCACCGCCTGCCGGATCGACTGCGAGACAA<br>GATACCTGGTGGAAAACTGCAACTGCCGGATGGTGCACATGCCTGG<br>CGACGCCCCTTACTGTACACCCGAGCAGTACAAAGAGTGCGCCGAT<br>CCTGCTCTGGACTTCCTGGTTGAGAAGGACCAAGAGTACTGCGTGTG<br>CGAGATGCCCTGCAACCTGACCAGATACGGCAAAGAACTGAGCATG<br>GTCAAGATCCCCAGCAAGGCCTCTGCCAAGTACCTGGCCAAGAAGT<br>TCAACAAGAGCGAGCAGTATATCGGCGAGAACATCCTGGTGCTGGA<br>TATCTTCTTCGAGGTGCTGAACTACGAGACAATCGAGCAGAAGAAG<br>GCCTACGAGATCGCCGGCCTGCTGGGAGATATTGGCGGACAGATGG<br>GCCTGTTTATCGGCGCCAGCATCCTGACCGTGCTGGAACTGTTCGAC<br>TACGCCTACGAAGTGATCAAGCACCGGCTGTGCAGACGGGGCAAGT<br>GTCAGAAAGAGGCCAAGAGAAACAGCGCCGACAAAGGCGTGGCCC<br>TGAGCCTGGATGATGTGAAGAGACACAACCCCTGCGAGAGCCTGAG<br>AGGACATCCTGCCGGAATGACCTACGCCGCCAACATTCTGCCTCATC<br>ACCCTGCCAGAGGCACCTTCGAGGACTTCACCTGTTGA |
| *Mja*KlbA-VidaL | N-Fragment:<br>mASIC1a_(1-193)-<br>*Mja*KlbA$^{N}$ | Same as *Mja*KlbA- *Ssp*DnaB$^{M86}$ split intein pair |

| | | |
|---|---|---|
| | Peptide X:<br>*Mja*KlbA$^{C}$-mASIC1a_(194-213)-VidaL$^{N}$-linker-ER retention sequence | Amino acid sequence:<br>MSSGSSINLDEVIKVETVDYNGHIYDLTVEDNHTYIAGKNEGFAVSNCY<br>TFNSGQDGRPRLKTMKGGESGALPKEAVVQIRLTKKGSSGESKDEL*<br><br>Nucleotide sequence:<br>ATGAGCTCTGGCTCTAGCATTAATCTGGATGAAGTGATTAAAGTGG<br>AAACAGTGGATTATAATGGACACATTTATGATCTGACAGTGGAAGA<br>TAATCACACATATATTGCTGGAAAAAATGAAGGATTTGCTGTGTCTA<br>ATTGCTACACCTTCAATAGCGGCCAGGACGGCAGACCCCGGCTGAA<br>AACAATGAAGGGCGGCGAATCTGGAGCTCTGCCAAAAGAAGCTGTG<br>GTGCAGATTAGACTGACAAAAAAAGGAAGCTCTGGCGAGAGCAAG<br>GACGAACTGTAA |
| | C-fragment:<br>IgK *faux* TMD-HA tag-linker-VidaL$^{C}$-mASIC1a_(214-526) | Amino acid sequence:<br>METDTLLLWVLLLWVPGSTGDYPYDVPDYAGSAGSAAGSGEFMIEEK<br>KVTVQELRELYLSGEYTIEIDTPDGYQTIGKWFDKGVLSMVRVATATY<br>ETVCAFNHMIQLADNTWVQACELDVGVDIQTAAGIQPVMLVEDTSDA<br>ECYDFEVMHPNHRYYGDGIVSHNSGNGLEIMLDIQQDEYLPVWGETDE<br>TSFEAGIKVQIHSQDEPPFIDQLGFGVAPGFQTFVSCQEQRLIYLPSPWG<br>TCNAVTMDSDFFDSYSITACRIDCETRYLVENCNCRMVHMPGDAPYCT<br>PEQYKECADPALDFLVEKDQEYCVCEMPCNLTRYGKELSMVKIPSKAS<br>AKYLAKKFNKSEQYIGENILVLDIFFEVLNYETIEQKKAYEIAGLLGDIG<br>GQMGLFIGASILTVLELFDYAYEVIKHRLCRRGKCQKEAKRNSADKGV<br>ALSLDDVKRHNPCESLRGHPAGMTYAANILPHHPARGTFEDFTC*<br><br>Nucleotide sequence:<br>ATGGAAACGGACACCCTGCTGCTGTGGGTGCTGTTGTTGTGGGTGCC<br>AGGCAGCACAGGCGACTACCCTTACGATGTGCCTGATTACGCCGGC<br>AGCGCTGGATCTGCTGCTGGAAGCGGAGAGTTTATGATTGAAGAAA<br>AAAAAGTGACAGTGCAGGAACTGAGAGAACTGTATCTGTCTGGAGA<br>ATATACAATTGAAATTGATACACCAGATGGATATCAGACAATTGGA<br>AAATGGTTTGATAAAGGAGTGCTGTCTATGGTGAGAGTGGCTACAG<br>CTACATATGAAACAGTGTGTGCTTTTAATCACATGATTCAGCTGGCT<br>GATAATACATGGGTGCAGGCTTGTGAACTGGATGTGGGAGTGGATA<br>TTCAGACAGCTGCTGGAATTCAGCCAGTGATGCTGGTGGAAGATAC<br>ATCTGATGCTGAATGTTATGATTTTGAAGTGATGCACCCAAATCACA<br>GATATTATGGAGATGGAATTGTGTCTCACAATTCTGGAAAAGGCCT<br>GGAAATCATGCTGGACATTCAGCAGGACGAGTACCTGCCTGTGTGG<br>GGCGAGACAGACGAGACATCTTTTGAGGCCGGCATCAAGGTGCAGA<br>TCCACAGCCAGGATGAGCCTCCATTCATCGACCAGCTCGGCTTTGGA<br>GTGGCCCCTGGCTTTCAGACCTTCGTGTCCTGCCAAGAGCAGCGGCT<br>GATCTACCTGCCTTCTCCTTGGGGCACCTGTAACGCCGTGACCATGG<br>ACAGCGATTTCTTCGACAGCTACAGCATCACCGCCTGCCGGATCGAC<br>TGCGAGACAAGATACCTGGTGGAAAACTGCAACTGCCGGATGGTGC<br>ACATGCCTGGCGACGCCCCTTACTGTACACCCGAGCAGTACAAAGA<br>GTGCGCCGATCCTGCTCTGGACTTCCTGGTTGAGAAGGACCAAGAG<br>TACTGCGTGTGCGAGATGCCCTGCAACCTGACCAGATACGGCAAAG<br>AACTGAGCATGGTCAAGATCCCCAGCAAGGCCTCTGCCAAGTACCT<br>GGCCAAGAAGTTCAACAAGAGCGAGCAGTATATCGGCGAGAACATC<br>CTGGTGCTGGATATCTTCTTCGAGGTGCTGAACTACGAGACAATCGA<br>GCAGAAGAAGGCCTACGAGATCGCCGGCCTGCTGGGAGATATTGGC<br>GGACAGATGGGCCTGTTTATCGGCGCCAGCATCCTGACCGTGCTGG<br>AACTGTTCGACTACGCCTACGAAGTGATCAAGCACCGGCTGTGCAG<br>ACGGGGCAAGTGTCAGAAAGAGGCCAAGAGAAACAGCGCCGACAA<br>AGGCGTGGCCCTGAGCCTGGATGATGTGAAGAGACACAACCCCTGC<br>GAGAGCCTGAGAGGACATCCTGCCGGAATGACCTACGCCGCCAACA<br>TTCTGCCTCATCACCCTGCCAGAGGCACCTTCGAGGACTTCACCTGT<br>TGA |
| NrdJ1-*Ssp*DnaX | N-Fragment:<br>mASIC1a_(1-179)-NrdJ1$^{N}$ | Same as shown for NrdJ1-*Ssp*DnaB$^{M86}$ split intein pair |

| | | |
|---|---|---|
| | Peptide X:<br>NrdJ1$^{C}$-mASIC1a_(180-213)-*Ssp*DnaX$^{N}$-linker-ER retention sequence | Amino acid sequence:<br>MEAKTYIGKLKSRKIVSNEDTYDIQTSTHNFFANDILVHNSAEDFKVVF<br>TRYGKCYTFNSGQDGRPRLKTMKGGCLTGDSQVLTRSSGESKDEL<br><br>Nucleotide sequence:<br>ATGGAAGCTAAAACATATATTGGAAAACTGAAATCTAGAAAAATTG<br>TGTCTAATGAAGATACATATGATATTCAGACATCTACACACAATTTT<br>TTTGCTAATGATATTCTGGTGCACAATTCTGCCGAGGACTTCAAGGT<br>GGTGTTCACCAGATACGGCAAGTGCTACACCTTCAATAGCGGCCAG<br>GACGGCAGACCCCGGCTGAAAACAATGAAGGGCGGCTGCCTGACCG<br>GCGATAGCCAGGTGCTGACCCGCAGCTCTGGCGAGAGCAAGGACGA<br>ACTGTAA |
| | C-fragment:<br>IgK *faux* TMD-HA tag-linker-*Ssp*DnaX$^{C}$-mASIC1a_(214-526) | Same as shown for gp41-1-*Ssp*DnaX split intein pair |
| IMPDH-1-*Ssp*DnaX | N-Fragment:<br>mASIC1a_(1-179)-IMPDH-1$^{N}$ | Amino acid sequence:<br>MELKTEEEEVGGVQPVSIQAFASSSTLHGLAHIFSYERLSLKRALWALC<br>FLGSLAVLLCVCTERVQYYFCYHHVTKLDEVAASQLTFPAVTLCNLNE<br>FRFSQVSKNDLYHAGELLALLNNRYEIPDTQMADEKQLEILQDKANFR<br>SFKPKPFNMREFYDRAGHDIRDMLLSCHFRGEACCFVPGTLVNTENGL<br>KKIEEIKVGDKVFSHTGKLQEVVDTLIFDRDEEIISINGIDCTKNHEFYVI<br>DKENANRVNEDNIHLFARWVHAEELDMKKHLLIELE*<br><br>Nucleotide sequence:<br>ATGGAACTCAAGACCGAGGAAGAGGAAGTCGGCGGCGTTCAGCCA<br>GTGTCCATTCAGGCCTTTGCCAGCAGCAGCACACTGCACGGACTGG<br>CCCACATCTTCAGCTACGAGAGACTGAGCCTGAAGCGGGCCCTGTG<br>GGCTCTGTGTTTTCTGGGATCTCTGGCCGTGCTGCTGTGCGTGTGTA<br>CAGAGAGAGTGCAGTACTACTTCTGCTACCACCACGTGACCAAGCT<br>GGATGAGGTGGCCGCTTCTCAGCTGACATTCCCTGCCGTGACACTGT<br>GCAACCTGAACGAGTTCCGGTTCTCCCAGGTGTCCAAGAACGACCT<br>GTATCACGCCGGCGAACTGCTGGCCCTGCTGAACAACAGATACGAG<br>ATCCCCGACACACAGATGGCCGACGAGAAGCAGCTGGAAATTCTGC<br>AGGACAAGGCCAACTTCAGAAGCTTCAAGCCCAAGCCGTTCAACAT<br>GCGCGAGTTCTACGATAGAGCCGGCCACGACATCCGGGACATGCTG<br>CTGAGCTGTCACTTTAGAGGCGAGGCCTGTTGCTTTGTGCCGGGCAC<br>CCTGGTGAACACCGAAAACGGCCTGAAAAAAATTGAAGAAATTAAA<br>GTGGGCGATAAAGTGTTTAGCCATACCGGCAAACTGCAGGAAGTGG<br>TGGATACCCTGATTTTTGATCGCGATGAAGAAATTATTAGCATTAAC<br>GGCATTGATTGCACCAAAAACCATGAATTTTATGTGATTGATAAAG<br>AAAACGCGAACCGCGTGAACGAAGATAACATTCATCTGTTTGCGCG<br>CTGGGTGCATGCGGAAGAACTGGATATGAAAAAACATCTGCTGATT<br>GAACTGGAATGA |
| | Peptide X:<br>IMPDH-1$^{C}$-mASIC1a_(180-213)-*Ssp*DnaX$^{N}$-linker-ER retention sequence | Amino acid sequence:<br>MKFKLKEITSIETKHYKGKVHDLTVNQDHSYNVRGTVVHNSAEDFKV<br>VFTRYGKCYTFNSGQDGRPRLKTMKGGCLTGDSQVLTRSSGESKDEL*<br><br>Nucleotide sequence:<br>ATGAAGTTCAAGCTTAAAGAGATCACTTCAATCGAGACAAAGCACT<br>ACAAGGGGAAGGTCCACGACCTTACGGTAAATCAAGACCACTCTTA<br>CAATGTTCGGGGTACGGTCGTCCACAATTCTGCTGAAGATTTTAAAG<br>TTGTTTTCACTCGATATGGTAAATGTTATACATTTAACTCTGGTCAA<br>GATGGACGCCCCAGACTAAAGACCATGAAAGGTGGATGTCTAACAG<br>GAGACTCTCAAGTTCTTACTCGGTCAAGTGGTGAATCAAAAGATGA<br>GCTCTAG |
| | C-fragment:<br>IgK *faux* TMD-HA tag-linker- *Ssp*DnaX$^{C}$-mASIC1a_(214-526) | Same as shown for gp41-1-*Ssp*DnaX split intein pair |

| | | |
|---|---|---|
| NrdJ1-VidaL | N-Fragment:<br>mASIC1a_(1-179)-<br>NrdJ1$^{N}$ | Same as shown for NrdJ1-*Ssp*DnaB$^{M86}$ split intein pair |
| | Peptide X:<br>NrdJ1$^{C}$-<br>mASIC1a_(180-213)-<br>VidaL$^{N}$-linker-ER<br>retention sequence | Amino acid sequence:<br>MEAKTYIGKLKSRKIVSNEDTYDIQTSTHNFFANDILVHNSAEDFKVVF<br>TRYGKCYTFNSGQDGRPRLKTMKGGESGALPKEAVVQIRLTKKGSSGE<br>SKDEL*<br><br>Nucleotide sequence:<br>ATGGAAGCTAAAACATATATTGGAAAACTGAAATCTAGAAAAATTG<br>TGTCTAATGAAGATACATATGATATTCAGACATCTACACACAATTTT<br>TTTGCTAATGATATTCTGGTGCACAATTCTGCCGAGGACTTCAAGGT<br>GGTGTTCACCAGATACGGCAAGTGCTACACCTTCAATAGCGGCCAG<br>GACGGCAGACCCCGGCTGAAAACAATGAAGGGCGGCGAATCTGGA<br>GCTCTGCCAAAAGAAGCTGTGGTGCAGATTAGACTGACAAAAAAAG<br>GAAGCTCTGGCGAGAGCAAGGACGAACTGTAA |
| | C-fragment:<br>IgK *faux* TMD-HA<br>tag-linker-VidaL$^{C}$-<br>mASIC1a_(214-526) | Same as shown for *Mja*KlbA-VidaL split intein pair |
| *Ssp*GyrB-*Ssp*DnaX | N-Fragment:<br>mASIC1a_(1-179)-<br>*Ssp*GyrB$^{N}$ | Amino acid sequence:<br>MELKTEEEEVGGVQPVSIQAFASSSTLHGLAHIFSYERLSLKRALWALC<br>FLGSLAVLLCVCTERVQYYFCYHHVTKLDEVAASQLTFPAVTLCNLNE<br>FRFSQVSKNDLYHAGELLALLNNRYEIPDTQMADEKQLEILQDKANFR<br>SFKPKPFNMREFYDRAGHDIRDMLLSCHFRGEACCFSGDTLVALTDGR<br>SVSFEQLVEEEKQGKQNFCYTIRHDGSIGVEKIINARKTKTNAKVIKVTL<br>DNGESIICTPDHKFMLRDGSYKCAMDLTLDDSLMPLHRKISTTEDSGHA<br>MEAVLNYNHRIVNIEAVSETIDVYDIEVPHTHNFALAS*<br><br>Nucleotide sequence:<br>ATGGAACTCAAGACCGAGGAAGAGGAAGTCGGCGGCGTTCAGCCA<br>GTGTCCATTCAGGCCTTTGCCAGCAGCAGCACACTGCACGGACTGG<br>CCCACATCTTCAGCTACGAGAGACTGAGCCTGAAGCGGGCCCTGTG<br>GGCTCTGTGTTTTCTGGGATCTCTGGCCGTGCTGCTGTGCGTGTGTA<br>CAGAGAGAGTGCAGTACTACTTCTGCTACCACCACGTGACCAAGCT<br>GGATGAGGTGGCCGCTTCTCAGCTGACATTCCCTGCCGTGACACTGT<br>GCAACCTGAACGAGTTCCGGTTCTCCCAGGTGTCCAAGAACGACCT<br>GTATCACGCCGGCGAACTGCTGGCCCTGCTGAACAACAGATACGAG<br>ATCCCCGACACACAGATGGCCGACGAGAAGCAGCTGGAAATTCTGC<br>AGGACAAGGCCAACTTCAGAAGCTTCAAGCCCAAGCCGTTCAACAT<br>GCGCGAGTTCTACGATAGAGCCGGCCACGACATCCGGGACATGCTG<br>CTGAGCTGTCACTTTAGAGGCGAGGCCTGTTGTTTTTCTGGAGATAC<br>ATTAGTCGCTTTAACTGATGGTCGTAGCGTTAGCTTTGAGCAATTGG<br>TTGAAGAAGAAAAACAAGGAAAACAAAACTTTTGTTATACCATCCG<br>CCATGATGGTTCTATAGGGGTTGAAAAAATCATCAATGCCCGCAAA<br>ACAAAAACTAATGCGAAGGTAATCAAGGTTACGTTGGACAATGGTG<br>AGTCTATTATTTGCACCCCGGATCATAAATTCATGTTGCGGGATGGG<br>AGCTACAAATGTGCGATGGATTTAACTCTCGATGATTCGTTAATGCC<br>GTTACACCGAAAAATTTCGACTACGGAAGATTCTGGTCATGCGATG<br>GAAGCAGTATTAAATTACAATCACAGAATTGTAAATATTGAAGCTG<br>TGTCAGAAACAATCGATGTTTATGATATTGAGGTTCCCCACACCCAC<br>AATTTTGCTTTGGCAAGCTGA |
| | Peptide X:<br>*Ssp*GyrB$^{C}$-<br>mASIC1a_(180-213)-<br>*Ssp*DnaX$^{N}$-linker-ER<br>retention sequence | Amino acid sequence:<br>MGVFVHNSAEDFKVVFTRYGKCYTFNSGQDGRPRLKTMKGGCLTGDS<br>QVLTRSSGESKDEL*<br><br>Nucleotide sequence:<br>ATGGGAGTGTTTGTCCATAACAGCGCCGAGGACTTCAAGGTGGTGT<br>TCACCAGATACGGCAAGTGCTACACCTTCAATAGCGGCCAGGACGG<br>CAGACCCCGGCTGAAAACAATGAAGGGCGGCTGCCTGACCGGCGAT<br>AGCCAGGTGCTGACCCGCAGCTCTGGCGAGAGCAAGGACGAACTGT |

| | | |
|---|---|---|
| | | AA |
| | C-fragment: IgK *faux* TMD-HA tag-linker- *Ssp*DnaX$^{C}$-mASIC1a_(214-526) | Same as shown for gp41-1-*Ssp*DnaX split intein pair |

**5B. Resource Table T2:** List of native and optimized extein sequences for split inteins used in this study.

| Split intein used | Optimized +1, +2, +3 extein residues | Native +1, +2, +3 residues | Optimized +1, +2, +3 residues | Additional optimization at -3, -2, -1 positions |
|---|---|---|---|---|
| *Cfa*DnaE | C, F, N[1] | C, Y, T | C, F, N | - |
| *Ssp*DnaB-M86 | S, X, X[1] | S, G, N | S, G, N | - |
| gp41-1 | S, S, S[7] | S, A, E | S, S, S | - |
| IMPDH-1 | S, I, C[7] | S, A, E | S, I, C | - |
| *Ssp*DnaX | C, P, X[8] | C, G, N | C, P, T | - |
| *Npu*DnaE | C, F, N[9] | C, Y, T | C, F, N | - |
| *Ssp*GyrB | S, A, K[7] | S, A, E | S, A, K | - |
| *Nrd*J1 | S, E, I[7] | S, A, E | S, E, I | N, P, C |
| CatInt | C, E, F[10] | C, G, N | C, E, F | X, F, E |
| *Mja*KlbA | C, S, G[7] | C, Y, T | C, S, G | H, D, G |
| VidaL | S, G/S, E/K[11] | S, G, N | S, G, K | - |

**5C. Resource Table T3** Split intein linked mASIC1a constructs (Optimized sequence); changes in protein and nucleic acid sequences have been underlined.

| Split intein combination (Intein A - Intein B) | Name of construct | Sequence |
|---|---|---|
| *Cfa*DnaE-<br>*Ssp*DnaB$^{M86}$ | N-Fragment:<br>mASIC1a_(1-193)-<br>*Cfa*DnaE$^{N}$ | No sequence optimization required |
| | Peptide X:<br>*Cfa*DnaE$^{C}$-<br>mASIC1a_(194-213)-*Ssp*DnaB$^{M86\text{-}N}$-<br>linker-ER retention sequence<br><br>+1+2+3<br>C Y T > C F N<br>(for *Cfa*DnaE$^{C}$) | Amino acid sequence:<br>MVKIISRKSLGTQNVYDIGVEKDHNFLLKNGLVASNCFNFNSGQDGRP<br>RLKTMKGGCISGDSLISLASSGESKDEL*<br><br>Nucleotide sequence:<br>ATGGTCAAGATCATCAGCCGGAAGTCCCTGGGCACCCAGAACGTG<br>TACGATATCGGCGTGGAAAAGGACCACAACTTTCTGCTGAAGAAC<br>GGCCTGGTGGCCAGCAACTGCTTTAACTTCAATAGCGGCCAGGACG<br>GCAGACCCCGGCTGAAAACAATGAAGGGCGGCTGCATCAGCGGCG<br>ACAGCCTGATTTCTCTGGCCAGCTCTGGCGAGAGCAAGGACGAACT<br>GTAA |
| | C-fragment:<br>*Ssp*DnaB$^{M86_C}$-<br>mASIC1a_(214-526) | No sequence optimization required |
| NrdJ1-<br>*Ssp*DnaB$^{M86}$ | N-Fragment:<br>mASIC1a_(1-179)-<br>NrdJ1$^{N}$<br><br>-3 -2 -1<br>E A C > N P C<br>(for NrdJ1$^{C}$) | Amino acid sequence:<br>MELKTEEEEVGGVQPVSIQAFASSSTLHGLAHIFSYERLSLKRALWAL<br>CFLGSLAVLLCVCTERVQYYFCYHHVTKLDEVAASQLTFPAVTLCNL<br>NEFRFSQVSKNDLYHAGELLALLNNRYEIPDTQMADEKQLEILQDKA<br>NFRSFKPKPFNMREFYDRAGHDIRDMLLSCHFRGNPCCLVGSSEIITRN<br>YGKTTIKEVVEIFDNDKNIQVLAFNTHTDNIEWAPIKAAQLTRPNAEL<br>VELEIDTLHGVKTIRCTPDHPVYTKNRGYVRADELTDDDELVVAI*<br><br>Nucleotide sequence:<br>ATGGAACTCAAGACCGAGGAAGAGGAAGTCGGCGGCGTTCAGCCA<br>GTGTCCATTCAGGCCTTTGCCAGCAGCAGCACACTGCACGGACTGG<br>CCCACATCTTCAGCTACGAGAGACTGAGCCTGAAGCGGGCCCTGTG<br>GGCTCTGTGTTTTCTGGGATCTCTGGCCGTGCTGCTGTGCGTGTGTA<br>CAGAGAGAGTGCAGTACTACTTCTGCTACCACCACGTGACCAAGCT<br>GGATGAGGTGGCCGCTTCTCAGCTGACATTCCCTGCCGTGACACTG<br>TGCAACCTGAACGAGTTCCGGTTCTCCCAGGTGTCCAAGAACGACC<br>TGTATCACGCCGGCGAACTGCTGGCCCTGCTGAACAACAGATACGA<br>GATCCCCGACACACAGATGGCCGACGAGAAGCAGCTGGAAATTCT<br>GCAGGACAAGGCCAACTTCAGAAGCTTCAAGCCCAAGCCGTTCAA<br>CATGCGCGAGTTCTACGATAGAGCCGGCCACGACATCCGGGACAT<br>GCTGCTGAGCTGTCACTTTAGAGGCAATCCATGTTGTCTGGTGGGA<br>TCTTCTGAAATTATTACAAGAAATTATGGAAAAACAACAATTAAAG<br>AAGTGGTGGAAATTTTTGATAATGATAAAAATATTCAGGTGCTGGC<br>TTTTAATACACACACAGATAATATTGAATGGGCTCCAATTAAAGCT<br>GCTCAGCTGACAAGACCAAATGCTGAACTGGTGGAACTGGAAATT<br>GATACACTGCACGGAGTGAAAACAATTAGATGTACACCAGATCAC<br>CCAGTGTATACAAAAAATAGAGGATATGTGAGAGCTGATGAACTG<br>ACAGATGATGATGAACTGGTGGTGGCTATTTGA |
| | Peptide X:<br>NrdJ1$^{C}$-<br>mASIC1a_(180-213)-*Ssp*DnaB$^{M86\text{-}N}$-<br>linker-ER retention sequence | Amino acid sequence:<br>MEAKTYIGKLKSRKIVSNEDTYDIQTSTHNFFANDILVHNSEIDFKVVF<br>TRYGKCYTFNSGQDGRPRLKTMKGGCISGDSLISLASSGESKDEL*<br><br>Nucleotide sequence:<br>ATGGAAGCTAAAACATATATTGGAAAACTGAAATCTAGAAAAATT<br>GTGTCTAATGAAGATACATATGATATTCAGACATCTACACACAATT |

| | | |
|---|---|---|
| | +1+2+3<br>S A E > S E I<br>(for NrdJ1$^{C}$) | TTTTTGCTAATGATATTCTGGTGCACAATTCTGAAATTGACTTCAAG<br>GTGGTGTTCACCAGATACGGCAAGTGCTACACCTTCAATAGCGGCC<br>AGGACGGCAGACCCCGGCTGAAAACAATGAAGGGCGGCTGCATCA<br>GCGGCGACAGCCTGATTTCTCTGGCCAGCTCTGGCGAGAGCAAGG<br>ACGAACTGTAA |
| | C-fragment:<br>*Ssp*DnaB$^{M86_C}$-mASIC1a_(214-526) | No sequence optimization required |
| *Mja*KlbA-*Ssp*DnaB$^{M86}$ | N-Fragment:<br>mASIC1a_(1-193)-*Mja*KlbA$^{N}$<br><br>-3 -2 -1<br>Y G K > H D G<br>(for *Mja*KlbA$^{N}$) | Amino acid sequence:<br>MELKTEEEEVGGVQPVSIQAFASSSTLHGLAHIFSYERLSLKRALWAL<br>CFLGSLAVLLCVCTERVQYYFCYHHVTKLDEVAASQLTFPAVTLCNL<br>NEFRFSQVSKNDLYHAGELLALLNNRYEIPDTQMADEKQLEILQDKA<br>NFRSFKPKPFNMREFYDRAGHDIRDMLLSCHFRGEACSAEDFKVVFTR<br>HDGALAYDEPIYLSDGNIINIGEFVDKFFKKYKNSIKKEDNGFGWIDIG<br>NENIYIKSFNKLSLIIEDKRILRVWRKKYSGKLIKITTKNRREITLTHDHP<br>VYISKTGEVLEINAEMVKVGDYIYIPKNNT*<br><br>Nucleotide sequence:<br>ATGGAACTCAAGACCGAGGAAGAGGAAGTCGGCGGCGTTCAGCCA<br>GTGTCCATTCAGGCCTTTGCCAGCAGCAGCACACTGCACGGACTGG<br>CCCACATCTTCAGCTACGAGAGACTGAGCCTGAAGCGGGCCCTGTG<br>GGCTCTGTGTTTTCTGGGATCTCTGGCCGTGCTGCTGTGCGTGTGTA<br>CAGAGAGAGTGCAGTACTACTTCTGCTACCACCACGTGACCAAGCT<br>GGATGAGGTGGCCGCTTCTCAGCTGACATTCCCTGCCGTGACACTG<br>TGCAACCTGAACGAGTTCCGGTTCTCCCAGGTGTCCAAGAACGACC<br>TGTATCACGCCGGCGAACTGCTGGCCCTGCTGAACAACAGATACGA<br>GATCCCCGACACACAGATGGCCGACGAGAAGCAGCTGGAAATTCT<br>GCAGGACAAGGCCAACTTCAGAAGCTTCAAGCCCAAGCCGTTCAA<br>CATGCGCGAGTTCTACGATAGAGCCGGCCACGACATCCGGGACAT<br>GCTGCTGAGCTGTCACTTTAGAGGCGAGGCCTGTAGCGCCGAGGAC<br>TTCAAGGTGGTGTTCACCAGACACGATGGAGCTCTGGCTTATGATG<br>AACCAATTTATCTGTCTGATGGAAATATTATTAATATTGGAGAGTT<br>CGTAGACAAGTTCTTCAAGAAGTACAAGAACTCCATCAAGAAAGA<br>GGATAATGGATTTGGATGGATTGATATTGGAAATGAAAATATTTAT<br>ATTAAATCTTTTAATAAACTGTCTCTGATTATTGAAGATAAAAGAA<br>TTCTGAGAGTGTGGAGAAAAAAATATTCTGGAAAACTGATTAAAA<br>TTACAACAAAAAATAGAAGAGAAATTACACTGACACACGATCACC<br>CAGTGTATATTTCTAAAACAGGAGAAGTGCTGGAAATTAATGCTGA<br>AATGGTGAAAGTGGGAGATTATATTTATATTCCAAAAAATAATACA<br>TGA |
| | Peptide X:<br>*Mja*KlbA$^{C}$-mASIC1a_(194-213)-*Ssp*DnaB$^{M86\text{-}N}$-linker-ER retention sequence<br><br>+1+2+3<br>C Y T > C S G<br>(for *Mja*KlbA$^{C}$) | Amino acid sequence:<br>MSSGSSINLDEVIKVETVDYNGHIYDLTVEDNHTYIAGKNEGFAVSNC<br>SGFNSGQDGRPRLKTMKGGTGCATCAGCGGCGACAGCCTGATTTCT<br>CTGGCCSSGESKDEL*<br><br>Nucleotide sequence:<br>ATGAGCTCTGGCTCTAGCATTAATCTGGATGAAGTGATTAAAGTGG<br>AAACAGTGGATTATAATGGACACATTTATGATCTGACAGTGGAAG<br>ATAATCACACATATATTGCTGGAAAAAATGAAGGATTTGCTGTGTC<br>TAATTGCTCTGGATTCAATAGCGGCCAGGACGGCAGACCCCGGCTG<br>AAAACAATGAAGGGCGGCTGCATCAGCGGCGACAGCCTGATTTCT<br>CTGGCCAGCTCTGGCGAGAGCAAGGACGAACTGTAA |
| | C-fragment:<br>*Ssp*DnaB$^{M86_C}$-mASIC1a_(214-526) | No sequence optimization required |
| *Npu*DnaE-*Ssp*DnaB$^{M86}$ | N-Fragment:<br>mASIC1a_(1-193)-*Npu*DnaE$^{N}$ | No sequence optimization required |

| | | |
|---|---|---|
| | Peptide X:<br>*Npu*DnaE$^{C}$-mASIC1a_(194-213)-*Ssp*DnaB$^{M86\text{-}N}$-linker-ER retention sequence<br><br>+1+2+3<br>C Y T > C F N<br>(for *Npu*DnaE$^{C}$) | Amino acid sequence:<br>MDHNFALKNGFIASNCFNFNSGQDGRPRLKTMKGGCISGDSLISLASSGESKDEL*<br><br>Nucleotide sequence:<br>ATGGATCATAACTTTGCGCTGAAAAACGGCTTTATTGCGAGCAACTGCTTCAACTTCAATAGCGGCCAGGACGGCAGACCCCGGCTGAAAACAATGAAGGGCGGCTGCATCAGCGGCGACAGCCTGATTTCTCTGGCCAGCTCTGGCGAGAGCAAGGACGAACTGTAA |
| | C-fragment:<br>*Ssp*DnaB$^{M86_C}$-mASIC1a_(214-526) | No sequence optimization required |
| gp41-1-*Ssp*DnaX | N-Fragment:<br>mASIC1a_(1-179)-gp41-1$^{N}$ | No sequence optimization required |
| | Peptide X:<br>gp41-1$^{C}$-mASIC1a_(180-213)-*Ssp*DnaX$^{N}$-linker-ER retention sequence<br><br>+1+2<br>A E > S S<br>(for gp41-1$^{C}$) | Amino acid sequence:<br>MMLKKILKIEELDERELIDIEVSGNHLFYANDILTHNSSSDFKVVFTRYGKCYTFNSGQDGRPRLKTMKGGCLTGDSQVLTRSSGESKDEL*<br><br>Nucleotide sequence:<br>ATGCTGAAAAAAATTCTGAAAATTGAAGAACTGGATGAACGCGAACTGATTGATATTGAAGTGAGCGGCAACCATCTGTTTTATGCGAACGATATTCTGACCCATAACAGCAGCAGCGACTTCAAGGTGGTGTTCACCAGATACGGCAAGTGCTACACCTTCAATAGCGGCCAGGACGGCAGACCCCGGCTGAAAACAATGAAGGGCGGCTGCCTGACCGGCGATAGCCAGGTGCTGACCCGCAGCTCTGGCGAGAGCAAGGACGAACTGTAA |
| | C-fragment:<br>*Ssp*DnaX$^{C}$-mASIC1a_(214-526)<br><br>+1+2+3<br>C G N > C P T<br>(for *Ssp*DnaX$^{C}$) | Amino acid sequence:<br>METDTLLLWVLLLWVPGSTGDYPYDVPDYAGSAGSAAGSGEFNGLMSIDNPQIKGREVLSYNETLQQWEYKKVLRWLDRGEKQTLSIKTKNSTVRCTANHLIRTEQGWTRAENITPGMKILSPAPQWHTNFEEVESVTKGQVEKVYDLEVEDNHNFVANGLLVHNCPTGLEIMLDIQQDEYLPVWGETDETSFEAGIKVQIHSQDEPPFIDQLGFGVAPGFQTFVSCQEQRLIYLPSPWGTCNAVTMDSDFFDSYSITACRIDCETRYLVENCNCRMVHMPGDAPYCTPEQYKECADPALDFLVEKDQEYCVCEMPCNLTRYGKELSMVKIPSKASAKYLAKKFNKSEQYIGENILVLDIFFEVLNYETIEQKKAYEIAGLLGDIGGQMGLFIGASILTVLELFDYAYEVIKHRLCRRGKCQKEAKRNSADKGVALSLDDVKRHNPCESLRGHPAGMTYAANILPHHPARGTFEDFTC*<br><br>Nucleotide sequence:<br>ATGGAAACGGACACCCTGCTGCTGTGGGTGCTGTTGTTGTGGGTGCCAGGCAGCACAGGCGACTACCCTTACGATGTGCCTGATTACGCCGGCAGCGCTGGATCTGCTGCTGGAAGCGGAGAGTTTAACGGCCTGATGAGCATTGATAACCCGCAGATTAAAGGCCGCGAAGTGCTGAGCTATAACGAAACCCTGCAGCAGTGGGAATATAAAAAAGTGCTGCGCTGGCTGGATCGCGGCGAAAAACAGACCCTGAGCATTAAAACCAAAAACAGCACCGTGCGCTGCACCGCGAACCATCTGATTCGCACCGAACAGGGCTGGACCCGCGCGGAAAACATTACCCCGGGCATGAAAATTCTGAGCCCGGCGCCGCAGTGGCATACCAACTTTGAAGAAGTGGAAAGCGTGACCAAAGGCCAGGTGGAAAAAGTGTATGATCTGGAAGTGGAAGATAACCATAACTTTGTGGCGAACGGCCTGCTGGTGCATAACTGCCCTACAGGCCTGGAAATCATGCTGGACATTCAGCAGGACGAGTACCTGCCTGTGTGGGGCGAGACAGACGAGACATCTTTTGAGGCCGGCATCAAGGTGCAGATCCACAGCCAGGATGAGCCTCCATTCATCGACCAGCTCGGCTTTGGAGTGGCCCCTGGCTTTCAGACCTTCGTGTCCTGCCAAGAGCAGCGGCTGATCTACCTGCCTTCTCCTTGGGGCACCTGTAACGCCGTGACCATGGACAGCGATTTCTTCGACAGCTACAGCATCACCGCCTGCCGGATCGACTGCGAGACAAGATACCTGGTGGAAAACTGCAACTGCCGGATGGTGCACATGCCTGGCGACGCCCCTTACTGTACACCCGAGCAGTACAAAGAGTGCGCCGATCCTGCTCTGGACTTCCTGGTTGAGAAGGACCAAGAGTACTGCGTGTGCGAGATGCCCTGCAACCT |

| | | |
|---|---|---|
| | | GACCAGATACGGCAAAGAACTGAGCATGGTCAAGATCCCCAGCAA<br>GGCCTCTGCCAAGTACCTGGCCAAGAAGTTCAACAAGAGCGAGCA<br>GTATATCGGCGAGAACATCCTGGTGCTGGATATCTTCTTCGAGGTG<br>CTGAACTACGAGACAATCGAGCAGAAGAAGGCCTACGAGATCGCC<br>GGCCTGCTGGGAGATATTGGCGGACAGATGGGCCTGTTTATCGGCG<br>CCAGCATCCTGACCGTGCTGGAACTGTTCGACTACGCCTACGAAGT<br>GATCAAGCACCGGCTGTGCAGACGGGGCAAGTGTCAGAAAGAGGC<br>CAAGAGAAACAGCGCCGACAAAGGCGTGGCCCTGAGCCTGGATGA<br>TGTGAAGAGACACAACCCCTGCGAGAGCCTGAGAGGACATCCTGC<br>CGGAATGACCTACGCCGCCAACATTCTGCCTCATCACCCTGCCAGA<br>GGCACCTTCGAGGACTTCACCTGTTGA |
| gp41-1-CatInt | N-Fragment:<br>mASIC1a_(1-179)-<br>gp41-1$^{N}$ | No sequence optimization required |
| | Peptide X:<br>gp41-1$^{C}$-<br>mASIC1a_(180-<br>213)-CatInt$^{N}$-linker-<br>ER retention<br>sequence<br><br>+1+2<br>A E > S S<br>(for gp41-1$^{C}$)<br><br>-2 -1<br>G G > F E<br>(for CatInt$^{N}$) | Amino acid sequence:<br>MLKKILKIEELDERELIDIEVSGNHLFYANDILTHNSSSDFKVVFTRYG<br>KCYTFNSGQDGRPRLKTMKFECLSGDTMIEILDDDGIIQKISMEDLYQR<br>LASSGESKDEL*<br><br>Nucleotide sequence:<br>ATGCTGAAAAAAATTCTGAAAATTGAAGAACTGGATGAACGCGAA<br>CTGATTGATATTGAAGTGAGCGGCAACCATCTGTTTTATGCGAACG<br>ATATTCTGACCCATAACAGCAGCAGCGACTTCAAGGTGGTGTTCAC<br>CAGATACGGCAAGTGCTACACCTTCAATAGCGGCCAGGACGGCAG<br>ACCCCGGCTGAAAACAATGAAGTTTGAATGCCTGAGCGGCGATAC<br>CATGATTGAAATTCTGGATGATGATGGCATTATTCAGAAAATTAGC<br>ATGGAAGATCTGTATCAGCGCCTGGCGAGCTCTGGCGAGAGCAAG<br>GACGAACTGTAA |
| | C-fragment:<br>CatInt$^{C}$-<br>mASIC1a_(214-<br>526)<br><br>+1+2+3<br>C G N > C E F<br>(for CatInt$^{C}$) | Amino acid sequence:<br>METDTLLLWVLLLWVPGSTGDYPYDVPDYAGSAGSAAGSGEFMFKL<br>NTKNIKVLTPSGFKSFSGIQKVYKPFYHHIIFDDGSEIKCSDNHSFGKDK<br>IKASTIKVGDYLQGKKVLYNEIVEEGIYLYDLLNVGEDNLYYTNGIVS<br>HNCEFGLEIMLDIQQDEYLPVWGETDETSFEAGIKVQIHSQDEPPFIDQ<br>LGFGVAPGFQTFVSCQEQRLIYLPSPWGTCNAVTMDSDFFDSYSITAC<br>RIDCETRYLVENCNCRMVHMPGDAPYCTPEQYKECADPALDFLVEKD<br>QEYCVCEMPCNLTRYGKELSMVKIPSKASAKYLAKKFNKSEQYIGENI<br>LVLDIFFEVLNYETIEQKKAYEIAGLLGDIGGQMGLFIGASILTVLELFD<br>YAYEVIKHRLCRRGKCQKEAKRNSADKGVALSLDDVKRHNPCESLR<br>GHPAGMTYAANILPHHPARGTFEDFTC*<br><br>Nucleotide sequence:<br>ATGGAAACGGACACCCTGCTGCTGTGGGTGCTGTTGTTGTGGGTGC<br>CAGGCAGCACAGGCGACTACCCTTACGATGTGCCTGATTACGCCGG<br>CAGCGCTGGATCTGCTGCTGGAAGCGGAGAGTTTATGTTTAAACTG<br>AACACCAAAAACATTAAAGTGCTGACCCCGAGCGGCTTTAAAAGC<br>TTTAGCGGCATTCAGAAAGTGTATAAACCGTTTTATCATCATATTAT<br>TTTTGATGATGGCAGCGAAATTAAATGCAGCGATAACCATAGCTTT<br>GGCAAAGATAAAATTAAAGCGAGCACCATTAAAGTGGGCGATTAT<br>CTGCAGGGCAAAAAAGTGCTGTATAACGAAATTGTGGAAGAAGGC<br>ATTTATCTGTATGATCTGCTGAACGTGGGCGAAGATAACCTGTATT<br>ATACCAACGGCATTGTGAGCCATAACTGCGGCAACGGCCTGGAAA<br>TCATGCTGGACATTCAGCAGGACGAGTACCTGCCTGTGTGGGGCGA<br>GACAGACGAGACATCTTTTGAGGCCGGCATCAAGGTGCAGATCCA<br>CAGCCAGGATGAGCCTCCATTCATCGACCAGCTCGGCTTTGGAGTG<br>GCCCCTGGCTTTCAGACCTTCGTGTCCTGCCAAGAGCAGCGGCTGA<br>TCTACCTGCCTTCTCCTTGGGGCACCTGTAACGCCGTGACCATGGA<br>CAGCGATTTCTTCGACAGCTACAGCATCACCGCCTGCCGGATCGAC<br>TGCGAGACAAGATACCTGGTGGAAAACTGCAACTGCCGGATGGTG<br>CACATGCCTGGCGACGCCCCTTACTGTACACCCGAGCAGTACAAAG<br>AGTGCGCCGATCCTGCTCTGGACTTCCTGGTTGAGAAGGACCAAGA<br>GTACTGCGTGTGCGAGATGCCCTGCAACCTGACCAGATACGGCAA<br>AGAACTGAGCATGGTCAAGATCCCCAGCAAGGCCTCTGCCAAGTA |

| | | |
|---|---|---|
| | | CCTGGCCAAGAAGTTCAACAAGAGCGAGCAGTATATCGGCGAGAA<br>CATCCTGGTGCTGGATATCTTCTTCGAGGTGCTGAACTACGAGACA<br>ATCGAGCAGAAGAAGGCCTACGAGATCGCCGGCCTGCTGGGAGAT<br>ATTGGCGGACAGATGGGCCTGTTTATCGGCGCCAGCATCCTGACCG<br>TGCTGGAACTGTTCGACTACGCCTACGAAGTGATCAAGCACCGGCT<br>GTGCAGACGGGGCAAGTGTCAGAAAGAGGCCAAGAGAAACAGCG<br>CCGACAAAGGCGTGGCCCTGAGCCTGGATGATGTGAAGAGACACA<br>ACCCCTGCGAGAGCCTGAGAGGACATCCTGCCGGAATGACCTACG<br>CCGCCAACATTCTGCCTCATCACCCTGCCAGAGGCACCTTCGAGGA<br>CTTCACCTGTTGA |
| *Mja*KlbA-VidaL | N-Fragment:<br>mASIC1a_(1-193)-<br>*Mja*KlbA$^{N}$<br><br>-3 -2 -1<br>Y G K > H D G<br>(for *Mja*KlbA$^{N}$) | Same as in *Mja*KlbA- *Ssp*DnaB$^{M86}$ split intein pair |
| | Peptide X:<br>*Mja*KlbA$^{C}$-<br>mASIC1a_(194-<br>213)-VidaL$^{N}$-linker-<br>ER retention<br>sequence<br>+1+2+3<br>C Y T > C S G<br>(for *Mja*KlbA$^{C}$)<br><br>-2<br>G > S<br>(for VidaL$^{N}$) | Amino acid sequence:<br>MSSGSSINLDEVIKVETVDYNGHIYDLTVEDNHTYIAGKNEGFAVSNC<br>SGFNSGQDGRPRLKTMKSGESGALPKEAVVQIRLTKKGSSGESKDEL*<br><br>Nucleotide sequence:<br>ATGAGCTCTGGCTCTAGCATTAATCTGGATGAAGTGATTAAAGTGG<br>AAACAGTGGATTATAATGGACACATTTATGATCTGACAGTGGAAG<br>ATAATCACACATATATTGCTGGAAAAAATGAAGGATTTGCTGTGTC<br>TAATGCTCTGGATTCAATAGCGGCCAGGACGGCAGACCCCGGCTG<br>AAAACAATGAAGAGCGGCGAATCTGGAGCTCTGCCAAAAGAAGCT<br>GTGGTGCAGATTAGACTGACAAAAAAAGGAAGCTCTGGCGAGAGC<br>AAGGACGAACTGTAA |
| | C-fragment:<br>VidaL$^{C}$-<br>mASIC1a_(214-<br>526)<br><br>+3<br>N > K<br>(VidaL$^{C}$) | Amino acid sequence:<br>METDTLLLWVLLLWVPGSTGDYPYDVPDYAGSAGSAAGSGEFMIEEK<br>KVTVQELRELYLSGEYTIEIDTPDGYQTIGKWFDKGVLSMVRVATATY<br>ETVCAFNHMIQLADNTWVQACELDVGVDIQTAAGIQPVMLVEDTSD<br>AECYDFEVMHPNHRYYGDGIVSHNSGKGLEIMLDIQQDEYLPVWGET<br>DETSFEAGIKVQIHSQDEPPFIDQLGFGVAPGFQTFVSCQEQRLIYLPSP<br>WGTCNAVTMDSDFFDSYSITACRIDCETRYLVENCNCRMVHMPGDAP<br>YCTPEQYKECADPALDFLVEKDQEYCVCEMPCNLTRYGKELSMVKIP<br>SKASAKYLAKKFNKSEQYIGENILVLDIFFEVLNYETIEQKKAYEIAGL<br>LGDIGGQMGLFIGASILTVLELFDYAYEVIKHRLCRRGKCQKEAKRNS<br>ADKGVALSLDDVKRHNPCESLRGHPAGMTYAANILPHHPARGTFEDF<br>TC*<br><br>Nucleotide sequence:<br>ATGGAAACGGACACCCTGCTGCTGTGGGTGCTGTTGTTGTGGGTGC<br>CAGGCAGCACAGGCGACTACCCTTACGATGTGCCTGATTACGCCGG<br>CAGCGCTGGATCTGCTGCTGGAAGCGGAGAGTTTATGATTGAAGA<br>AAAAAAAGTGACAGTGCAGGAACTGAGAGAACTGTATCTGTCTGG<br>AGAATATACAATTGAAATTGATACACCAGATGGATATCAGACAATT<br>GGAAAATGGTTTGATAAAGGAGTGCTGTCTATGGTGAGAGTGGCT<br>ACAGCTACATATGAAACAGTGTGTGCTTTTAATCACATGATTCAGC<br>TGGCTGATAATACATGGGTGCAGGCTTGTGAACTGGATGTGGGAGT<br>GGATATTCAGACAGCTGCTGGAATTCAGCCAGTGATGCTGGTGGAA<br>GATACATCTGATGCTGAATGTTATGATTTTGAAGTGATGCACCCAA<br>ATCACAGATATTATGGAGATGGAATTGTGTCTCACAATTCTGGAAA<br>AGGCCTGGAAATCATGCTGGACATTCAGCAGGACGAGTACCTGCCT<br>GTGTGGGGCGAGACAGACGAGACATCTTTTGAGGCCGGCATCAAG<br>GTGCAGATCCACAGCCAGGATGAGCCTCCATTCATCGACCAGCTCG<br>GCTTTGGAGTGGCCCCTGGCTTTCAGACCTTCGTGTCCTGCCAAGA<br>GCAGCGGCTGATCTACCTGCCTTCTCCTTGGGGCACCTGTAACGCC |

| | | |
|---|---|---|
| | | GTGACCATGGACAGCGATTTCTTCGACAGCTACAGCATCACCGCCT<br>GCCGGATCGACTGCGAGACAAGATACCTGGTGGAAAACTGCAACT<br>GCCGGATGGTGCACATGCCTGGCGACGCCCCTTACTGTACACCCGA<br>GCAGTACAAAGAGTGCGCCGATCCTGCTCTGGACTTCCTGGTTGAG<br>AAGGACCAAGAGTACTGCGTGTGCGAGATGCCCTGCAACCTGACC<br>AGATACGGCAAAGAACTGAGCATGGTCAAGATCCCCAGCAAGGCC<br>TCTGCCAAGTACCTGGCCAAGAAGTTCAACAAGAGCGAGCAGTAT<br>ATCGGCGAGAACATCCTGGTGCTGGATATCTTCTTCGAGGTGCTGA<br>ACTACGAGACAATCGAGCAGAAGAAGGCCTACGAGATCGCCGGCC<br>TGCTGGGAGATATTGGCGGACAGATGGGCCTGTTTATCGGCGCCAG<br>CATCCTGACCGTGCTGGAACTGTTCGACTACGCCTACGAAGTGATC<br>AAGCACCGGCTGTGCAGACGGGGCAAGTGTCAGAAAGAGGCCAAG<br>AGAAACAGCGCCGACAAAGGCGTGGCCCTGAGCCTGGATGATGTG<br>AAGAGACACAACCCCTGCGAGAGCCTGAGAGGACATCCTGCCGGA<br>ATGACCTACGCCGCCAACATTCTGCCTCATCACCCTGCCAGAGGCA<br>CCTTCGAGGACTTCACCTGTTGA |
| NrdJ1-*Ssp*DnaX | N-Fragment:<br>mASIC1a_(1-179)-<br>NrdJ1$^{N}$<br><br>-3-2-1<br>E A C > N P C | Same as in NrdJ1- *Ssp*DnaB$^{M86}$ split intein pair |
| | Peptide X:<br>NrdJ1$^{C}$-<br>mASIC1a_(180-<br>213)-*Ssp*DnaX$^{N}$-<br>linker-ER retention<br>sequence<br><br>+1+2+3<br>S A E > S E I<br>(for NrdJ1$^{C}$) | Amino acid sequence:<br>MEAKTYIGKLKSRKIVSNEDTYDIQTSTHNFFANDILVHNSEIDFKVVF<br>TRYGKCYTFNSGQDGRPRLKTMKGGCLTGDSQVLTRSSGESKDEL*<br><br>Nucleotide sequence:<br>ATGGAAGCTAAAACATATATTGGAAAACTGAAATCTAGAAAAATT<br>GTGTCTAATGAAGATACATATGATATTCAGACATCTACACACAATT<br>TTTTTGCTAATGATATTCTGGTGCACAATTCTGAAATTGACTTCAAG<br>GTGGTGTTCACCAGATACGGCAAGTGCTACACCTTCAATAGCGGCC<br>AGGACGGCAGACCCCGGCTGAAAACAATGAAGGGCGGCTGCCTGA<br>CCGGCGATAGCCAGGTGCTGACCCGCAGCTCTGGCGAGAGCAAGG<br>ACGAACTGTAA |
| | C-fragment:<br>*Ssp*DnaX$^{C}$-<br>mASIC1a_(214-<br>526)<br><br>+1+2+3<br>C G N > C P T<br>(for *Ssp*DnaX$^{C}$) | Same as in gp41-1-*Ssp*DnaX split intein pair |
| IMPDH-1-<br>*Ssp*DnaX | N-Fragment:<br>mASIC1a_(1-179)-<br>IMPDH-1$^{N}$ | No sequence optimization required |
| | Peptide X:<br>IMPDH-1$^{C}$-<br>mASIC1a_(180-<br>213)-*Ssp*DnaX$^{N}$-<br>linker-ER retention<br>sequence<br><br>+1+2+3<br>S A E > S I C<br>(for IMPDH-1$^{C}$) | Amino acid sequence:<br>MKFKLKEITSIETKHYKGKVHDLTVNQDHSYNVRGTVVHNSICDFKV<br>VFTRYGKCYTFNSGQDGRPRLKTMKGGCLSGDTMIEILDDDGIIQKIS<br>MEDLYQRLASSGESKDEL*<br><br>Nucleotide sequence:<br>ATGAAATTTAAACTGAAAGAAATTACCAGCATTGAAACCAAACAT<br>TATAAAGGCAAAGTGCATGATCTGACCGTGAACCAGGATCATAGC<br>TATAACGTGCGCGGCACCGTGGTGCATAACAGCATCTGCGACTTCA<br>AGGTGGTGTTCACCAGATACGGCAAGTGCTACACCTTCAATAGCGG<br>CCAGGACGGCAGACCCCGGCTGAAAACAATGAAGGGCGGCTGCCT<br>GAGCGGCGATACCATGATTGAAATTCTGGATGATGATGGCATTATT<br>CAGAAAATTAGCATGGAAGATCTGTATCAGCGCCTGGCGAGCTCTG<br>GCGAGAGCAAGGACGAACTGTAA |
| | C-fragment: | Same as in gp41-1-*Ssp*DnaX split intein pair |

| | | |
|---|---|---|
| | *Ssp*DnaX$^{C}$-mASIC1a_(214-526)<br><br>+1+2+3<br>C G N > C P T<br>(for *Ssp*DnaX$^{C}$) | |
| NrdJ1-VidaL | N-Fragment:<br>mASIC1a_(1-179)-NrdJ1$^{N}$ | Same as in NrdJ1- *Ssp*DnaB$^{M86}$ split intein pair |
| | Peptide X:<br>NrdJ1$^{C}$-mASIC1a_(180-213)-VidaL$^{N}$-linker-ER retention sequence<br><br>+1+2+3<br>S A E > S E I<br>(for NrdJ1$^{C}$)<br><br>-2<br>G > S<br>(for VidaL$^{N}$) | Amino acid sequence:<br>MEAKTYIGKLKSRKIVSNEDTYDIQTSTHNFFANDILVHNSEIDFKVVF<br>TRYGKCYTFNSGQDGRPRLKTMKGGESGALPKEAVVQIRLTKKGSSG<br>ESKDEL*<br><br>Nucleotide sequence:<br>ATGGAAGCTAAAACATATATTGGAAAACTGAAATCTAGAAAAATT<br>GTGTCTAATGAAGATACATATGATATTCAGACATCTACACACAATT<br>TTTTTGCTAATGATATTCTGGTGCACAATTCTGAAATTGACTTCAAG<br>GTGGTGTTCACCAGATACGGCAAGTGCTACACCTTCAATAGCGGCC<br>AGGACGGCAGACCCCGGCTGAAAACAATGAAGAGCGGCGAATCTG<br>GAGCTCTGCCAAAAGAAGCTGTGGTGCAGATTAGACTGACAAAAA<br>AAGGAAGCTCTGGCGAGAGCAAGGACGAACTGTAA |
| | C-fragment:<br>VidaL$^{C}$-mASIC1a_(214-526)<br><br>+3<br>N > K<br>(VidaL$^{C}$) | Same as in *Mja*KlbA-VidaL split intein pair |
| *Ssp*GyrB-*Ssp*DnaX | N-Fragment:<br>mASIC1a_(1-179)-*Ssp*GyrB$^{N}$ | No sequence optimization required |
| | Peptide X:<br>*Ssp*GyrB$^{C}$-mASIC1a_(180-213)-*Ssp*DnaX$^{N}$-linker-ER retention sequence<br><br>+3<br>E > K<br>(for *Ssp*GyrB$^{C}$) | Amino acid sequence:<br>MGVFVHNSAKDFKVVFTRYGKCYTFNSGQDGRPRLKTMKGGCLTGD<br>SQVLTRSSGESKDEL*<br><br>Nucleotide sequence:<br>ATGGGAGTGTTTGTCCATAACAGCGCCAAGGACTTCAAGGTGGTGT<br>TCACCAGATACGGCAAGTGCTACACCTTCAATAGCGGCCAGGACG<br>GCAGACCCCGGCTGAAAACAATGAAGGGCGGCTGCCTGACCGGCG<br>ATAGCCAGGTGCTGACCCGCAGCTCTGGCGAGAGCAAGGACGAAC<br>TGTAA |
| | C-fragment:<br>*Ssp*DnaX$^{C}$-mASIC1a_(214-526)<br><br>+1+2+3<br>C G N > C P T<br>(for *Ssp*DnaX$^{C}$) | Same as in gp41-1-*Ssp*DnaX split intein pair |

#### 6. References:

1. Khoo, K.K., Galleano, I., Gasparri, F., Wieneke, R., Harms, H., Poulsen, M.H., Chua, H.C., Wulf, M., Tampe, R., and Pless, S.A. (2020). Chemical modification of proteins by insertion of synthetic peptides using tandem protein trans-splicing. Nat Commun *11*, 2284. 10.1038/s41467-020-16208-6.
2. Flood, D.T., Yan, N.L., and Dawson, P.E. (2018). Post-Translational Backbone Engineering through Selenomethionine-Mediated Incorporation of Freidinger Lactams. Angew Chem Int Ed Engl *57*, 8697-8701. 10.1002/anie.201804885.
3. Hopkins, C.E., Hernandez, G., Lee, J.P., and Tolan, D.R. (2005). Aminoethylation in model peptides reveals conditions for maximizing thiol specificity. Arch Biochem Biophys *443*, 1-10. 10.1016/j.abb.2005.08.020.
4. Lynagh, T., Flood, E., Boiteux, C., Wulf, M., Komnatnyy, V.V., Colding, J.M., Allen, T.W., and Pless, S.A. (2017). A selectivity filter at the intracellular end of the acid-sensing ion channel pore. Elife *6*. 10.7554/eLife.24630.
5. Dahan, D.S., Dibas, M.I., Petersson, E.J., Auyeung, V.C., Chanda, B., Bezanilla, F., Dougherty, D.A., and Lester, H.A. (2004). A fluorophore attached to nicotinic acetylcholine receptor beta M2 detects productive binding of agonist to the alpha delta site. Proc Natl Acad Sci U S A *101*, 10195-10200. 10.1073/pnas.0301885101.
6. Liwocha, J., Krist, D.T., van der Heden van Noort, G.J., Hansen, F.M., Truong, V.H., Karayel, O., Purser, N., Houston, D., Burton, N., Bostock, M.J., et al. (2021). Linkage-specific ubiquitin chain formation depends on a lysine hydrocarbon ruler. Nat Chem Biol *17*, 272-279. 10.1038/s41589-020-00696-0.
7. Pinto, F., Thornton, E.L., and Wang, B. (2020). An expanded library of orthogonal split inteins enables modular multi-peptide assemblies. Nat Commun *11*, 1529. 10.1038/s41467-020-15272-2.
8. Zhang, X., Liu, X.Q., and Meng, Q. (2019). Engineered Ssp DnaX inteins for protein splicing with flanking proline residues. Saudi J Biol Sci *26*, 854-859. 10.1016/j.sjbs.2017.07.010.
9. Iwai, H., Zuger, S., Jin, J., and Tam, P.H. (2006). Highly efficient protein trans-splicing by a naturally split DnaE intein from Nostoc punctiforme. FEBS Lett *580*, 1853-1858. 10.1016/j.febslet.2006.02.045.
10. Stevens, A.J., Sekar, G., Gramespacher, J.A., Cowburn, D., and Muir, T.W. (2018). An Atypical Mechanism of Split Intein Molecular Recognition and Folding. J Am Chem Soc *140*, 11791-11799. 10.1021/jacs.8b07334.
11. Burton, A.J., Haugbro, M., Parisi, E., and Muir, T.W. (2020). Live-cell protein engineering with an ultra-short split intein. Proc Natl Acad Sci U S A *117*, 12041-12049. 10.1073/pnas.2003613117.
